# *De novo* mutation rates at the single-mutation resolution in a human *HBB* gene-region associated with adaptation and genetic disease

**DOI:** 10.1101/2021.05.24.443729

**Authors:** Daniel Melamed, Yuval Nov, Assaf Malik, Michael B. Yakass, Evgeni Bolotin, Revital Shemer, Edem K. Hiadzi, Karl L. Skorecki, Adi Livnat

## Abstract

While it is known that the mutation rate varies across the genome, previous estimates were based on averaging across various numbers of positions. Here we describe a method to measure the origination rates of target mutations at target base positions and apply it to a 6-bp region in the human ***β***-globin (*HBB*) gene and to the identical, homologous ***δ***-globin (*HBD*) region in sperm cells from both African and European donors. The *HBB* region of interest (ROI) includes the site of the hemoglobin S (HbS) mutation, which protects against malaria, is common in Africa and has served as a classic example of adaptation by random mutation and natural selection. We found a significant correspondence between *de novo* mutation rates and past observations of alleles in carriers, showing that mutation rates vary substantially in a mutation-specific manner that contributes to the site frequency spectrum. We also found that the overall point mutation rate is significantly higher in Africans than Europeans in the *HBB* region studied. Finally, the rate of the 20A***→***T mutation, called the “HbS mutation” when it appears in *HBB*, is significantly higher than expected from the genome-wide average for this mutation type. Nine instances were observed in the African *HBB* ROI, where it is of adaptive significance, representing at least three independent originations, and no instances were observed in the European *HBB* ROI or in the European or African *HBD* ROI. Further studies will be needed to examine *de novo* mutation rates at the single-mutation resolution across these and other loci and organisms and to uncover the molecular mechanisms responsible.

---

It is widely known that mutation rates vary across the genome at multiple scales (*1-3*) and are affected by multiple factors, from the mutation type (*4, 5*), to the local genetic context (*2-7*) to the general location in the genome (*8-11*). Although this knowledge is highly advanced now compared to a mere decade ago (*2, 3, 12–14*), it could be further improved. In particular, rate measurements to date have all been based on averages of various kinds, such as an average across the genome (*2, 15*), across the instances of any particular motif (*3, 7*), or across the entire stretch of a gene in certain cases (*16-18*), whereas technological limitations have precluded measuring the mutation rates at particular base positions and of particular mutations at such positions. Such high-resolution knowledge of the mutation-rate variation would bear on multiple open questions in genetics and evolution—from the relative importance of mutation-rate variation to the site frequency spectrum (SFS) (*19-21*), to its importance to adaptive evolution and parallelism (*22-29*), to its contribution to recurrent genetic disease (*30-33*).

The most precise way of measuring mutation rates, free of biases due to past natural selection or random genetic drift events, is offered by *de novo* mutations, because they appear for the first time in their carrier and are not inherited (*2, 34*). They are usually detected by studies comparing the genomes of children and their parents (”trio studies”) (*35,36*). However, because each individual carries only a small number (e.g., several dozen in humans) of *de novo* mutations scattered across the genome, the chance of encountering any particular target mutation of interest is miniscule, rendering it impractical to measure rates of target mutations using such studies.

To overcome this barrier, we have developed a method that enables identification and counting at high accuracy ultra-rare genetic variants of choice in extremely narrow regions of interest (ROIs) within large populations of cells, such as a single target mutant in 100 million genomes. Since this method has both an error rate lower than the human mutation rate and sufficient yield for the purpose, it enables measuring the evolutionarily relevant origination rates of target mutations of choice in males by counting *de novo* mutations in human sperm cells at the single-digit resolution. Note that aside from this evolutionary application, ultra-accurate methods of mutation-detection are sought after for early detection of cancer, fetal diagnostic tests through maternal blood, early virus identification within host, and more (*37*).

As a first target for this method, we chose two sites: a 6-bp region spanning 3 codons within the human *β*-globin (*HBB*) gene that is of great importance for adaptation and hematologic disease, and the identical, homologous region within the *δ*-globin (*HBD*) gene. The former region includes, among others, the site of the HbS mutation. The most iconic balanced polymorphism mutation (*38-43*), the HbS mutation is an A to T transversion (GAG*→*GTG, Glu*→*Val) in codon 6 of *HBB* causing sickle-cell anemia in homozygotes (*38*) and providing substantial protection against severe malaria in heterozygotes (*40, 44–46*). Malaria, in turn, has been a leading cause of human morbidity and mortality, often causing more than a million deaths per year in recent past, with Africa bearing the brunt of the disease burden (*47*), and thus has been the strongest known agent of selection in humans in recent history (*46*). Besides the HbS mutation, many other mutations, both point mutations and indels, are also known at this site, many of which are involved in hematologic illness (*48, 49*). In contrast to *HBB*, mutations in *HBD* have a more limited effect and are not thought to confer malarial resistance, as the *HBD*’s lower expression levels make it account for less than 3% of the circulating red blood cell hemoglobin in adults (*50*). While the population prevalence of the *HBB* mutations, whether beneficial or detrimental, is normally attributed to natural selection, so far it has not been possible to examine to what degree, if at all, mutational phenomena may also be relevant to their prevalence. Here, we sought to characterize the rates of mutations, including the HbS mutation, in the *HBB* and *HBD* ROIs in sperm samples of both African and European donors.

Using the method described below, we examined a total of more than half a billion gene fragments individually for the origination patterns of *de novo* mutations in the two ROIs. As described below, results show that mutation rates vary substantially between specific mutations, between the two ROIs and between the two populations. A correspondence is obtained at an ultra-high resolution between *de novo* rates and prevalence of alleles in populations, underscoring the importance of mutation-specific rates to the SFS. The overall point mutation rate in the *HBB* ROI is significantly higher than expected and is significantly higher in the African than in the European population. The HbS mutation has a rate significantly higher than expected from the genome-wide average for its type, even after adjusting for the local genetic context. Consistent with the significant population-level difference in the overall point mutation rates, specifically in our samples we observed the HbS mutation *de novo* more frequently in the African than in the European samples. These results and more establish the ability to measure and demonstrate the importance of measuring mutation rate variation at the single-mutation resolution.

## Methods

After extracting DNA from the sperm of the donors (SI Appendix, Table S1), each sample is enriched for the mutation of interest and nearby mutations. Specifically for the target sites, the restriction enzyme Bsu36I is used, which cleaves the wild-type (WT) sequence CCTGAGG at positions 16–22 of *HBB* and the homologous positions of *HBD* while leaving the HbS mutant and other mutants in these positions intact. This step reduces substantially both the probability of false positives, i.e., a PCR or sequencing error appearing as a mutation, and the sequencing cost (Fig 1, SI Appendix, Figs. S1–S4).

**Figure 1:**
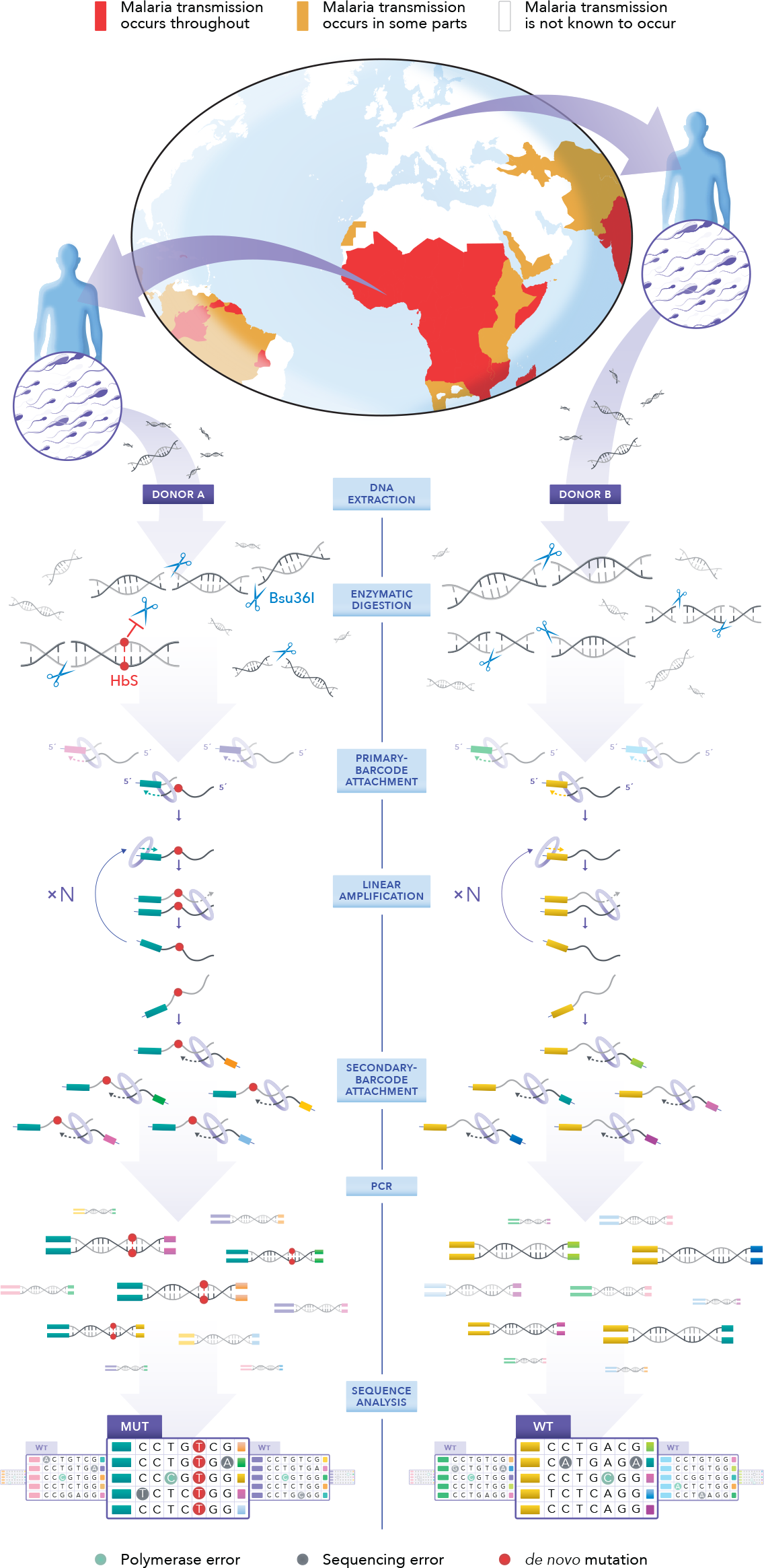
Experiment overview. Sperm samples are obtained from world regions with high or low malaria infection burden (malaria impact map adjusted from the CDC map; ref. 74). Whole-genome DNA is extracted and an amount equivalent to 60–80 million sperm cells per donor is subjected to Bsu36I digestion. Bsu36I cleaves the DNA at multiple sites, including the *HBB* and *HBD* ROIs, which carry a specific recognition sequence. The HbS mutation blocks Bsu36I digestion and is thus enriched over the wild-type (WT). A primary barcode is added directly to each antisense DNA strand that carries the *HBB* or *HBD* ROI via a DNA-polymerase– assisted fill-in reaction. Since each barcode consists of a random sequence of nucleotides, each of the numerous target fragments has its own, unique barcode. Multiple single-strand copies are each generated directly from each uniquely barcoded target fragment by linear amplification. A secondary barcode composed of a random sequence of nucleotides is added to the other end of each of these copies by a single primer-extension reaction. Thus, only full-length fragments (i.e., mutant or WT ROI sequences that evaded Bsu36I digestion) carry both the primary and the secondary barcodes and can be amplified by PCR for next-generation sequencing (NGS). At the sequence analysis step, sequencing reads are grouped based on their shared primary-barcode sequences. Sporadic DNA-polymerase errors and sequencing errors are unlikely to repeat in multiple copies and are removed. *De novo* mutations, such as the HbS mutation, are easily identified by their appearance in multiple reads from distinct linear-amplification events. For a complete description of the library preparation protocol, which includes additional steps, see SI Appendix supplementary Figs. S1–S3.

**Table 1:**
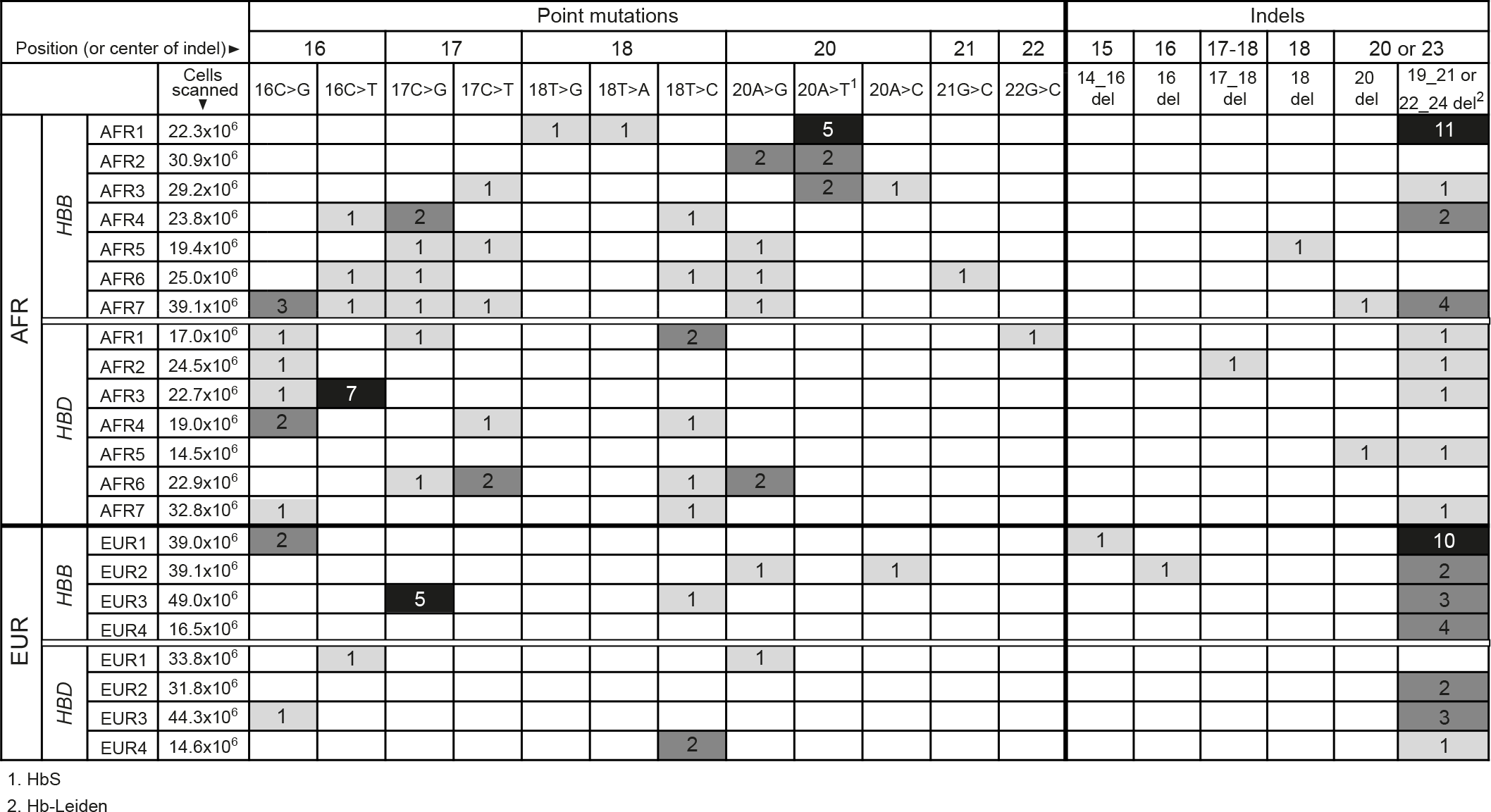
HBB and HBD ROI mutation counts. Counts of *de novo* mutations identified by MEMDS in DNA from 11 sperm samples, 7 from African (AFR) and 4 from European (EUR). The numbers next to the donor labels refer to the calculated number of haploid individual genomes scanned by MEMDS. Some of the mutations have been observed before in carriers and have common names when they appear in *HBB*. These are 16C→G, Hb-Gorwihl; 16C→T, Hb-Tyne; 17C→G, Hb-Warwickshire; 17C→T, Hb-Aix-les-Bains; 20A→G, Hb-Lavagna; 20A→T, HbS; 20A→C, Hb-G-Makassar; 22G→C, Hb Bellevue III and 19 21del or 22 24del, Hb-Leiden. Note that Hb-Leiden can result from deletion of either positions 19–21 or positions 22–24, which include the same GAG sequence, both of which can be enriched and captured by MEMDS.

Next, unique barcodes are attached to the DNA fragments in order to reduce error by consensus sequencing of copies originating from the same original fragment. To this end, we build on the Maximum Depth Sequencing method (MDS) (*51*), which allows one to focus on a narrow region of interest (ROI) and which attaches the barcodes directly to the cleaved end of one of the two strands of each original target DNA fragment via a DNA-polymerase–assisted extension reaction (*51*) (Fig. 1 and SI Appendix, Fig. S1). Additionally, multiple innovations are introduced that increase sequencing accuracy, handle the large amounts of DNA required and accurately measure the Bsu36I enrichment factor per sample as needed for the mutation rate calculation (SI Appendix, Figs. S1–S5). We refer to this method as Mutation Enrichment followed by upscaled Maximum Depth Sequencing—MEMDS (for a complete protocol, see Text S1–S9 and Materials and Methods in SI Appendix).

Finally, following sequence analysis (SI Appendix, Figs. S6–S10 and Table S2) the number of appearances of any mutation that confers resistance to the restriction enzyme is counted and divided by the calculated number of cells analyzed, providing the evolutionarily relevant *de novo* origination rate for each specific mutation in males per donor and per group of donors (SI Appendix, Figs. S11–S13 and Table S3). Following previous literature, we ignore G*→*T and C*→*T mutations in the barcoded strand (C*→*A and G*→*A in the sequenced strand) because they are thought to reflect not lasting mutations but the experimental disruption of an ongoing *in vivo* process of base damage and repair as well as *in vitro* mutations due to guanine oxidation and cytosine deamination (*51, 52*) (SI Appendix, Text S8 and Figs. S12–S14). In addition, we exclude C*→*A, the complement of G*→*T, due to its association with the latter and its frequent appearance in the data (SI Appendix, Text S8 and Fig. S12). Following normal loss of material of *∼*65%, true positives of non G*→*T, C*→*T and C*→*A mutations are identified with a false positive rate (error rate) *<* 2.5 × 10*^−^*^9^ per base (Fig. 2). Overall, MEMDS surpasses recent cutting-edge methods in both accuracy and yield (Fig. 2 and SI Appendix, Fig. S11).

**Figure 2:**
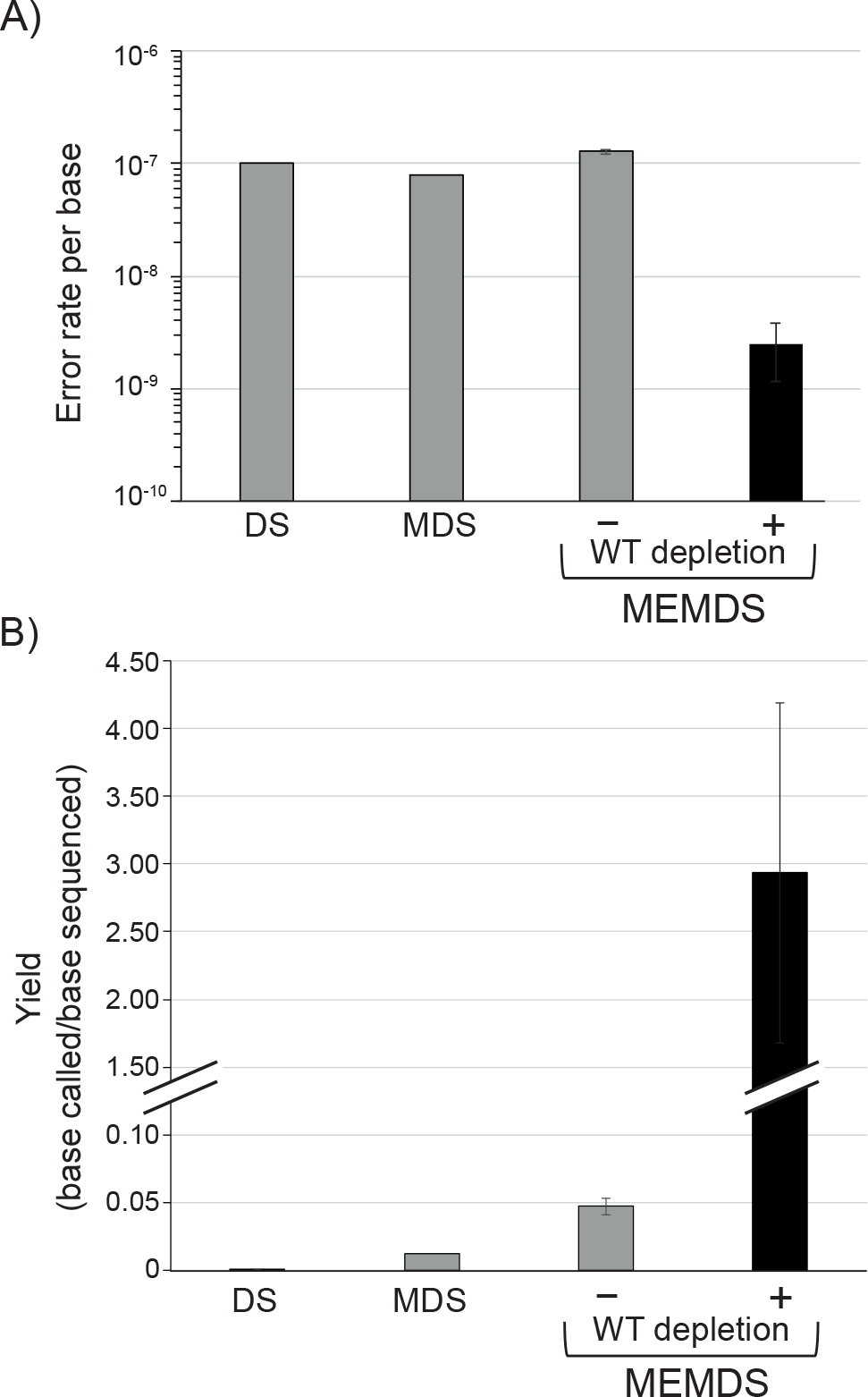
Accuracy and yield of MEMDS in comparison to current cutting-edge methods. A) Under a highly conservative estimate, MEMDS increases accuracy by at least 40-folds in comparison to Duplex Sequencing (DS) (*75*) and Maximum-Depth Sequencing (MDS) (*51*). MEMDS increases yield per sequenced base (i.e., the number of MEMDS confirmed bases divided by the number of paired-end sequenced bases) by orders of magnitude over both DS and MDS (*51,75*). Notice that in MEMDS, the yield can be higher than 1 because the mutation-enrichment factor is accurately calculated (see SI Appendix, Text S2) and the base identity is known for the ROI sequences that were digested and removed from the final sequencing libraries (they have the RE-1 recognition sequence).

We obtained sperm from 12 donors. Since one of the samples was a mixture from two African donors with a total number of cells similar to the other African samples, we consider it here as a single sample of mixed African origins, bringing the total to 11 samples, 7 from African and 4 from European donors (SI Appendix, Table S1).

## Results

The numbers of cells scanned and *de novo* mutations observed per person are shown in Table 1.

### Average per-ROI mutation rates

The average per-base point mutation rates in the *HBB* and *HBD* ROIs are 3.3 × 10*^−^*^8^ and 2.79 × 10*^−^*^8^ respectively, significantly higher by *∼*2.6-fold (*P <* 2 × 10*^−^*^8^, 95% CI 2.4 × 10*^−^*^8^–4.4 × 10*^−^*^8^) and *∼*2.2-fold (*P <* 6.7 × 10*^−^*^5^, 95% CI 1.9 × 10*^−^*^8^–4 × 10*^−^*^8^, two-sided binomial exact test) than 1.25 × 10*^−^*^8^, which we use as an estimate of the genome-wide per-base per-generation point-mutation rate (SI Appendix, Text S10). The average indel rates in these ROIs were 1.1 × 10*^−^*^8^ and 4.3 × 10*^−^*^9^ respectively, significantly higher by *∼*9-fold (*P <* 4.3 × 10*^−^*^25^, 95% CI 8 × 10*^−^*^9^–1.5 × 10*^−^*^8^) and *∼*3.4-fold (*P <* 1.8 × 10*^−^*^4^, 95% CI 2.3 × 10*^−^*^9^–7.3 × 10*^−^*^9^; two-sided binomial exact test) than the expected ^1^⁄10 of the point mutation rate (SI Appendix, Text S10). The average point mutation rate of the *HBB* ROI is not significantly higher than that of the *HBD* ROI (*P* = 0.49, two-sided Fisher exact test), and the average indel rate of the former is significantly higher by *∼*2.6-fold than that of the latter (*P* = 0.0015, OR 95% CI 1.42 – 5.01; two-sided Fisher exact test).

### Basic characteristics of mutation-rate variation

The variance in the rates of *de novo* point mutations is higher than expected from the genome-wide average (GWA) rates of these mutations (e.g., refs. *53, 54*) and their relative rates are different than expected from the GWA rates (*P <* 10*^−^*^6^ in an omnibus multinomial test, adjusted for the excluded mutations, compared to the rates of ref. 2), even when adjusting the latter for the 3-mer, 5-mer and 7-mer nucleotide contexts (*P <* 10*^−^*^5^ in all cases, compared to the rates of ref. 3). The overall *de novo* rates of the 6 observed deletion types are highly non-uniform (*P <* 10*^−^*^6^, multi-sample proportion test).

### Correspondence between *de novo* rates and observations of alleles in carriers

The HbS and Hb-Leiden mutations have both been notably observed on multiple different genetic backgrounds in human populations, the former particularly in Africans (*45, 48, 49*). Here, they are the point mutation of highest *de novo* rate in the African *HBB* ROI and the deletion mutation of highest *de novo* rate in any gene and ethnicity. Furthermore, of the 23 potential deletions of up to size 3 that are observable by our method per ROI, only 5 deletions have been reported on the HbVar database—a large collection of hemoglobin variants (*48, 49*)—all in *HBB*; and of these deletion types, a significantly higher fraction is observed here *de novo* compared to deletion types not reported on HbVar (SI Appendix, Text S11). Pooling together both the *HBB* and *HBD* ROIs given the similarity of *de novo* indel types observed between them, this effect is significant both with (*P* = 0.0078, OR 95% CI 2.17–818.08, two-sided Fisher exact test) and without (*P* = 0.024, OR 95% CI 1.44–653.93, two-sided Fisher exact test) the Hb-Leiden mutation, showing that the correspondence between *de novo* rates and alleles in populations extends beyond the HbS and Hb-Leiden mutations. The same analysis cannot be repeated for the point mutations, because of the synonymous vs. non-synonymous mutation confound and the smaller number of observable mutation types (SI Appendix, Text S11). Importantly, the correspondence observed could not have been predicted from the mutations’ GWA rates, also when adjusting for the genetic context (SI Appendix, Text S10–S11).

### Between-population comparisons

The per-person overall point mutation rates in the *HBB* ROI are significantly higher in the African than in the European group both with (*P* = 0.0061) and without (*P* = 0.043, two-sided Wilcoxon rank sum test) counting the HbS mutation. This excludes individual- or sample-level variation alone as accounting for this overall point-mutation rate difference and establishes a significant population-level difference between the groups. Pooling together cells from different donors within each population shows that the overall point-mutation rate in the *HBB* ROI is significantly (2.57×) higher in the African than in the European donors (*P <* 0.006, OR 95% CI 1.27–5.49, two-sided Fisher exact test). In the *HBD* ROI, the number of mutations is not high enough to establish a significant population-level difference as opposed to individual- or sample-level variation. In contrast to the *HBB* overall point mutation rate, the overall indel rate does not vary significantly between these groups in either ROI (P=0.35 and P=1, respectively, two-sided Fisher exact test).

### Rates of the Hb-Leiden mutation

The 3 bp in-frame deletion variant of either codon 6 or codon 7 that is called Hb-Leiden when it occurs in *HBB* recurs noticeably more often than other mutations (comparing its per-person rates to those of all other deletions combined, *P <* 0.0005, two-sided Wilcoxon rank sum test). Pooled across individuals, it appears at rates of 1.11 × 10*^−^*^7^ and 3.96 × 10*^−^*^8^ in the *HBB* and *HBD* ROIs respectively, *∼*88.86 and *∼*31.66 higher than the 1.25×10*^−^*^9^ estimate (*P* = 4.04×10*^−^*^58^, 95% CI 7.82×10*^−^*^8^–1.53×10*^−^*^7^; and *P* = 1.62×10*^−^*^13^, 95% CI 1.98 × 10*^−^*^8^–7.08 × 10*^−^*^8^), where the pooled *HBB* rate is significantly (*∼*2.81×) higher than the *HBD* rate (*P* = 0.002, OR 95% CI 1.40–5.63, two-sided Fisher exact test).

### Rates of the HbS mutation

The 20A*→*T mutation called “the HbS mutation” when it appears in the *HBB* ROI appears 9 times in the African *HBB* ROI and no times in the other cases combined (the European *HBB* ROI and the European and African *HBD* ROIs) (*P* = 0.023, 95% CI 1.5077–Inf; two-sided Fisher exact test classifying each individual and gene case as having [*>* 0] or not having [= 0] *de novo* 20A*→*T in sperm and comparing the fractions of these classes between the groups). The rate of the HbS mutation in the overall group (Africans and Europeans combined)—2.7 × 10*^−^*^8^—is 19.6 times higher (*P <* 2 × 10*^−^*^9^, rate 95% CI 1.24 × 10^08^–5.13 × 10*^−^*^8^) than expected from the GWA for this mutation type, and its rate in the African group specifically—4.74 × 10*^−^*^8^—is *∼*35× higher than expected from its GWA (*P* = 1.2 × 10*^−^*^11^, rate 95% CI 2.17 × 10*^−^*^8^–9.0 × 10*^−^*^8^; two-sided binomial exact test). In the African group, it is the mutation that deviates most (Table S4) from its GWA among the 12 observable point mutations, where its *de novo* rate varies significantly across samples (*P* = 0.0025, multi-sample proportion test), from 0 to 2.24 × 10*^−^*^7^ (the latter rate being *∼*163× faster than expected; *P* = 2.23 × 10*^−^*^10^, 95% CI 7.27 × 10*^−^*^8^–5.23 × 10*^−^*^7^, two-sided binomial exact test). Note that the evolutionarily relevant mutation rate depends on the fraction of the mutation in sperm, not on whether it repeats because of independent originations or due to an early appearance followed by duplications during spermatogenesis. That being said, the minimal number of independent originations of the HbS mutation is 3, and the corresponding minimal rate of independent occurrence of the HbS mutation in the sperm samples (a rate lower than the actual evolutionarily relevant mutation rate observed) is 9.01 × 10*^−^*^9^. This rate is *∼*6.5× higher than the genome-wide evolutionarily relevant mutation rate for this mutation type (*P* = 0.011, 95% rate CI 1.86 × 10*^−^*^9^–2.63 × 10*^−^*^8^, two-sided binomial exact test).

## Discussion

The data exposes an ultra-high resolution correspondence between *de novo* mutation rates and past observations of alleles in carriers (refs. 45, 48, 49, Results and SI Appendix, Text S11), suggesting that these rates contribute to the prevalence of these mutations in populations. This correspondence could not have been predicted from the GWA rates of these mutation types even when adjusting for the local genetic context (SI Appendix, Text S10–S11). Consideration of the deletions observed clarifies this point. While past literature featured a single microdeletion rate decreasing with size (*18, 55, 56*), sized-based rate-variation cannot explain the correspondence aforementioned obtained for same-sized deletions, the higher rate of the Hb-Leiden mutation compared to the smaller deletions or the extent of rate-variation observed. Thus, the correspondence aforementioned, together with the fact that the rates of some mutations (e.g., those of the HbS and Hb-Leiden mutations) deviate much more than others from these mutations’ GWA rates show that mutation-specific rates vary not only in the case of large rearrangement mutations (*57, 58*) but also in the cases of point mutations and microindels. This rate variation could not have been seen from average-based measures (*18, 56*) and establishes the relevance of mutation-specific point mutation and microindel rates to the site frequency spectrum (SFS) (*19-21*).

The significantly higher overall point mutation rate in Africans than in Europeans in the *HBB* ROI cannot be attributed to individual- or sample-level variance alone and thus represents a population-level difference between the groups. This difference in an extremely narrow region spanning 3 codons of great importance for adaptation and genetic disease is at least two orders of magnitude larger than previous differences in GWA mutation rates between continental groups (*53, 54*). Both the correspondence between mutation-specific *de novo* rates and observations of alleles in carriers and the large difference in the overall point mutation rate between populations in a narrow region establish the importance of measuring mutation-rate variation at an ultra-high resolution.

Potential contributions to mutation rates from gross-level biological or environmental factors such as age or pesticides cannot sufficiently explain the results. First, the two populations are similar in ages (Appendix, Table S1). Second, any mutation-specific effect, like the correspondence between *de novo* rates and observations of alleles in carriers, cannot be explained by such macro-level factors. Third, the overall point mutation rate difference between the populations is also unlikely to be explained by them, because if on their own such macro-level factors affected the ROIs, they should have affected the entire genome similarly, yet GWA differences in point mutation rates between continental groups are incomparably smaller than the ROI-specific differences observed here (*53,54*); and if macro-level factors affect mutation rates in interaction with mutation-, locus-, individual- and/or population-specific factors, then such specific factors must be assumed in any case. Thus, rather than suggesting involvement of macro-level factors, the data suggests a complex picture of mutation rates involving mutation-specific influences.

Although the replication of mutations during spermatogenesis (clonal dependence) may contribute to the data, in practice it does not explain the significant results. First, due to the statistical test used, the significance of the continental difference in the overall point mutation rates in *HBB* is impervious to any sample-level variation, including clonal dependence. Second, the correspondence between mutation rates and observations of alleles in carriers cannot be driven by it. Any presence of clonal dependence would only make it more difficult to obtain significance for such patterns and in that sense is conservative to such results. The significance of these patterns can only be driven by independent originations of the mutations. These independent originations are most suitably explained by mutation-specific rates being influenced by genetic and/or epigenetic factors (*59, 60*).

Note that the prevalence of a mutation of heterozygote advantage in a population and of reading-frame conservation in a coding sequence have generally been considered to be outcomes of selection. However, here, both the HbS mutation, which provides strong malaria protection in heterozygotes, and the Hb-Leiden mutation, which is an in-frame deletion, are frequent not because of selection but because of frequent *de novo* origination. Indeed, that the rate of the in-frame Hb-Leiden mutation is much higher than that of all other observed deletions, which are frameshift deletions, demonstrates reading-frame conservation that is not due to selection (*19*), but due to mutational phenomena. This observation provides a concrete example of “mutational conservation”—evolutionary conservation due to mutational reasons, which, if occurs more broadly, could offer an explanation for the puzzling observation of reading-frame conservation bias in pseudogenes (*61*).

The fact that the genetic sequences at and adjacent to the ROIs are identical across individuals, populations, and the two genes, yet the mutation rates vary significantly between these entities, suggests that what affects these mutation rates in the germline includes more than this local DNA sequence and in that sense is complex (*59,60*). These results are interestingly consistent with the observation that the variation of the mutation rates across loci is partly cryptic (not explained by the local DNA context) (*1, 62*), especially in the case of A*↔*T transversions (*62*), which include the HbS mutation-type (A*→*T). Combining the multiple insights discussed, the results suggest that mutation rates are both mutation-specific and influenced in a complex manner by the genetic and/or epigenetic background (*59, 60*).

The *HBB* region spanning 3 codons is of particular importance for adaptation and genetic disease: it is the site of mutations that provide strong protection against malaria (HbS and HbC, the latter not observable by our method) and/or mutations that increase the risk for hematologic disease (*45, 48, 49*). Thus, it is of interest that the overall point mutation rate in this region is significantly higher than expected, and that it is significantly higher in the African than in the European population. These facts stand to add to previous, pioneering experimental work on the relevance of locus-specific mutation rates to adaptive evolution and its repeatability (*23, 24, 26–29*). It has been shown, for example, that high rates of deletion of a pelvic enhancer of the Pitx1 gene are responsible for parallel and likely adaptive loss of the pelvic hindfin in freshwater sticklebacks (*26*), that mutationally frequent losses or gains of tandem duplicates of agrp2 exon 3 may have been repeatedly involved in color pattern diversification of cichlids (*27*), and that hypermutability of a HoxA13 duplicate was associated with its neofunctionalization in zebrafish and related taxa (*23*).

The results underscore the importance of mapping the mutation-rate variation at an ultra-high resolution. It is beyond this fact that several observations on the HbS mutation can be mentioned. First, under the assumption that the HbS rate is the same between the continental groups, it is significantly higher by nearly *∼* 20-fold than expected from the GWA for this mutation type, in both Africans and Europeans. Any amount of hypothetical clonal dependence does not change this estimate of the observed evolutionarily relevant mutation rate, because the latter does not depend on the cause of the recurrence of the mutation in the sperm. Even the observed minimal rate of independent HbS originations in sperm is still significantly larger by × than the evolutionarily relevant GWA rate for this mutation type. Consideration of the local genetic context does not change this conclusion (SI Appendix, Text S10). Thus, while the classical account of the HbS case relied only on selection, not on origination rate, as a part of the explanation, even under the most conservative assumptions the overall HbS mutation rate is notably higher than expected.

Second, given the continental difference in the overall point mutation rate between the groups, it would be surprising if the HbS mutation specifically does not show a continental effect. Consistent with this, in our samples, using the methodology described, we observe no instances of it in Europeans but 9 instances of it in total in Africans, amounting to a rate *∼*35× higher than expected from the GWA of this mutation type in the latter. Further consistent with a continental difference in the HbS mutation rate, it fits with the broader correspondence between *de novo* rates and observations of alleles in populations that HbS is most frequent in Africans and in some other populations in the Asian malaria belt (*45*) and appears *de novo* in our African but not in our European samples, while Hb-Leiden has been observed across the globe (*48, 49*) and appears *de novo* in both our African and European samples.

Third, in the African *HBB* ROI, out of 12 observable point mutations, the HbS mutation has the rate that deviates most from the corresponding GWA rate (Table S4).

Fourth, it is striking that despite at least three independent occurrences of the HbS mutation in the *HBB* ROI, not a single case of the equivalent 20A*→*T mutation in the *HBD* ROI was observed in any donor, African or European. Accordingly, we note that the binary test establishing the significantly higher concentration of the 20A*→*T mutation in the African *HBB* ROI as opposed to all other cases (the European *HBB* ROI or the *HBD* ROIs), impervious to any individual- or sample-level variance including clonal dependence, suggests that the 20A*→*T mutation arises more frequently where it is of adaptive significance than where it is not, though data does not suffice to tell whether this effect is due to a population-level difference or due to a locus-based difference or both.

Knowing that the HbS mutation is advantageous in heterozygotes under malarial pressure, how shall we interpret these results? One possibility is that, for a reason unrelated to adaptation, some individuals have a genomic fragility in *HBB* that generates the HbS mutation at a high rate. Accordingly, it is a coincidence that HbS provides protection against malaria, even more so if the fragility applies more to Africans.

Another possibility is modifier theory. However, it is hard to account for the results with it, because it does not predict an increase in the rate of specific DNA mutations at specific base positions in a sexual species (*1, 63, 64*) nor the complex genetic and/or epigenetic influences on such mutation rates suggested by the current data. On the contrary, the “reduction principle,” the first-order principle in modifier theory, underscores the general difficulty of accounting for increased mutation rates (*65, 66*).

Third, a recent proposal holds that mutation-specific origination rates are influenced by the complex genetic and epigenetic background and that long-term adaptive evolution involves joint evolution of adaptations and the complex spectrum of mutation-specific origination rates (*59, 60*). Among else, this proposal predicts the existence of genetic relatedness in mutational tendencies, the ability of mutation-specific rates to come to reflect past selection pressures, and the possibility that the HbS mutation originates more rapidly in African populations that have long experienced intense malarial selection pressure (*59, 60*). This proposal does not consider the HbS mutation rate to be adaptive as previously interpreted (*67, 68*). Rather, it raises the possibility that DNA or RNA regulation or modification processes that are themselves outcomes of evolution, such as RNA editing (*69*), epigenetic modifications (*70*) and other processes may lead directly to their own replacement and simplification via DNA mutations that arise in the course of evolution from these processes’ molecular mechanistic nature, thus mechanistically linking regulatory activity with structural mutational changes. We argue, for example, that A*→*I RNA editing can often be an early, transitory stage of adaptation (cf. ref. 71) that is followed in evolutionary time by A*→*G DNA changes in the edited positions arising under a higher A*→*G mutation rate caused by or sharing a cause with the editing (cf. ref. 72). This replacement hypothesis is an extension of the hypothesis that “genes that are used together are fused together”—that two genes whose products interact closely are more likely to undergo a fusion mutation in the course of evolution for mechanistic reasons (*60*). It suggests that a mutation like the HbS one may not arise as a random mutation that begins a process of adaptation but rather may arise as a later stage in an ongoing evolutionary process where adaptations and mutation rates jointly evolve (*59, 60*).

According to this last possibility, a missing element in studies on the potential connection between mutation origination and adaptation may have been that they looked for an immediate, Lamarckian adaptive mutational response to an environmental change during the lifetime of the organism (*67, 73*) rather than a response following multiple generations under selection as in the present study, where Africans have experienced intense malarial selection pressure for many generations. Unlike previous methods that could explore only diffuse relationships between long-term selection pressures and the evolution of GWA mutation rates, the present method offers the refined ability needed to explore such relationships, if they exist, at the mutation-specific resolution. This method likely applies across loci and organisms, given an RE suitable for the mutations of interest. Therefore, some of the most important tasks now are to examine the high-resolution mutation rate variation across other loci of interest and to explore the molecular mechanisms responsible.

## Conclusions

The origination rates of target mutations at target base positions of choice can now be measured provided a suitable RE for the task. The measurement is highly accurate in the *HBB* and *HBD* ROIs, exposing a broad correspondence between origination rates and observations of alleles in populations. Results show that mutation rates are mutation-specific and vary across positions, genes and populations more than expected from previous studies and in a manner largely unexplained by the local genetic context. The African and European populations studied differ markedly in their overall point mutation rates in the *HBB* ROI—an extremely narrow region of great importance for adaptation and genetic disease. A concrete example is given showing reading frame and thus amino acid sequence conservation that is not due to natural selection but due to mutational conservation. Observations on the HbS mutation rate show it to be significantly higher than expected from the genome-wide average for this mutation type, even when considering the local genetic context. It also appears to originate more where it is of adaptive significance, although the significance of this effect cannot yet be attributed to a continental or a locus effect or both. However, the higher overall point mutation rate in Africans in the extremely narrow *HBB* region studied, the correspondence between *de novo* mutation rates and observations of alleles in carriers, and the concentration of HbS mutation observations in our specific samples in the African donors suggest that the HbS mutation rate is higher in Africans than Europeans. A hypothesis is proposed connecting long-term selection pressures and mutation-specific rates. Future research is needed to increase our knowledge on the HbS mutation rate and to explore the extent of mutation-rate variation across these and other loci of interest, the dependence of mutation on the complex genetic and epigenetic background and the molecular mechanisms responsible.

## Acknowledgments

We thank Marc Feldman for comments on a previous draft, Rami Reshef for infrastructural resources, Mary Otoo and Joshua Adoboe for help with sample collection, Sara Zelig, Alan Templeton and Nick Pippenger for technical comments, and Kim Weaver for extensive help.

## Author contributions

DM and AL invented the method and designed the studies; DM performed all experiments except for RS’s; RS processed the EUR4 sample; YN created software tools for data analysis; AM created the computational pipeline for mutation calling; EB improved the pipeline; YN and AL provided statistical tools; MY, EH, KS and AL obtained IRB and Helsinki approvals; MY and EH collected samples; DM, YN, EB, AM and AL analyzed the results; DM and AL drafted the paper; DM, YN, KS and AL revised the draft; KS provided general advice; KS and AL acquired funding; AL conceived of the project and the replacement hypothesis and supervised the project.

## Data and materials availability

Raw data will be deposited in NCBI dbGaP. Software is available at https://github.com/livnat-lab/HBB HBD. All other data required for evaluating the conclusions of this manuscript is available either in the main text or in the supplementary materials.

## Supplementary Information

### 1. The MEMDS method

#### 1.1. Current high-accuracy sequencing methods

While Next Generation Sequencing (NGS) technology has tremendously improved the cost and scale of DNA sequencing, the detection of extremely rare genetic variants remains a major challenge. This unresolved problem is due to both DNA polymerase errors that are introduced during sample preparation and sequencing errors made by the NGS machinery. DNA polymerase error rates range between *∼*1 × 10*^−^*^4^ per base for Taq polymerase to *∼*1 × 10*^−^*^6^ per base for various high-fidelity DNA polymerases (*1–3*), and error rates of the commonly used NGS platforms range between 10*^−^*^2^-10*^−^*^3^ per sequenced base (*4*), with some computational efforts being able to enhance sequencing accuracy 10–100 fold (*5, 6*). These abilities allow for the detection of some sub-populations of sequences within a highly homogenous sample. However, given that the average per-base point mutation rate across the human genome is *∼*10*^−^*^8^ (see supplementary section 10), entailing on average *∼*10^8^ wild-type copies per mutant, the rates above would lead to numerous false positives per one true base mutation. Thus, detecting a single instance of a particular *de novo* mutation in a particular gene has been practically impossible so far. Moreover, even in the absence of errors, obtaining enough reads of such a mutation by sequencing alone would have entailed an exorbitant sequencing cost.

In recent years, a few experimental approaches have been developed that substantially reduce the noise generated by both DNA polymerase and NGS errors (*7–16*). One key idea has been to attach a unique molecular tag or “barcode” to each DNA fragment at the first PCR cycle during the amplification step. After library preparation and standard NGS sequencing, reads that share the same barcode are recognized as having been derived from the same original molecule. Since those reads should be identical, the differences between them are considered to be errors introduced during PCR and/or NGS sequencing and are filtered out at the analysis stage (*8*). This filtration step removes many of the DNA polymerase and NGS errors that occur after barcode attachment.

Importantly, however, the standard way by which the barcode has been added to the target DNA is by being included as a part of a target-specific primer that is extended by a single elongation reaction, generating a sequence subsequently to be amplified using an external pair of primers. A major disadvantage of this standard method is that any replication error introduced by the DNA polymerase during the critical, initial copying of the original DNA molecule is transferred to all downstream copies during the PCR reaction and cannot be filtered out by the regular barcoding-and-consensus-sequencing approach.

To overcome this problem, a few methods have been developed. In Duplex Sequencing (DS) (*13*), double-stranded barcodes are attached to both ends of a sheared DNA segment by ligation. This operation makes it possible, at the sequence analysis step, to group together the two sets of copies from the two strands of each original double-stranded DNA molecule based on their barcode complementarity. A consensus sequence is first constructed for each strand of an original double-stranded DNA molecule from its set of copies, and then the two consensus sequences are compared to each other. The identity of a base at each position is approved only if the two consensus sequences show a perfect match. This approach allows the DS method to capture errors that occur at any amplification and sequencing step, including the generation of the first copy of each original molecule, and to reach an error rate below *∼*2.5×10*^−^*^6^ when applied to the M13mp2 bacteriophage DNA (*13*) and an error rate of 1×10*^−^*^7^– 5 × 10*^−^*^8^ according to unpublished data (*17*). However, while ideal for genomic regions at the 1Mb scale, used on smaller regions, the hybridization-based capture that it uses to enable the attachment of duplex barcodes by ligation to double-stranded molecules would entail extremely low yields, which in turn would make it impossible to reduce the size of the region of interest (ROI) and increase the sequencing depth in a manner nearly cost-effective enough as to focus on a particular mutation of interest.

An alternative method that avoids errors in the first copying step, Maximum Depth Sequencing (MDS) (*16*), is based on single-strand sequencing. However, instead of generating a barcoded copy of each original DNA molecule by extending a barcoded primer, it adds the barcode to the original target DNA molecule itself by cleavage of the target molecule near the ROI followed by a fill-in reaction that extends the target-DNA strand using a barcoding oligo as a template. Next, linear amplification is performed to obtain multiple copies of the target molecule, each generated directly from the original, now barcoded, single-stranded target DNA molecule, and the preparation of this library is finally completed by a standard exponential PCR. Like DS, MDS consensus sequences reach all the way to the very original target DNA molecules, without an intervening, uncontrolled copying step. However, in contrast to DS, MDS can potentially minimize the ROI to a single base in the genome. Since MDS recovers the sequence information from only one of the two DNA strands, however, it cannot correct certain particular kinds of error due to DNA damage or base misincorporation by the cellular DNA polymerase that affect the target DNA strand, in contrast to DS. Yet, eliminating these highly frequent, known types of errors from the mutation rate calculation resulted in a tested MDS error rate of about 1 × 10*^−^*^7^ while using Phusion DNA polymerase, and a suggested theoretical error rate of less than 5 × 10*^−^*^8^ if Q5 DNA polymerase is used (*16*).

#### 1.2. MEMDS boosts both mutation detection accuracy and yield

While both DS and MDS reach extraordinary levels of precision, their error rates and sequence coverage demands still pose serious difficulties in detecting particular mutations occurring at or near the human genome-wide average mutation rate. In particular, note that natural mutation variants constitute a tiny fraction of the target DNA molecules, while the vast majority of the target DNA consists of a common, non-mutated sequence, which we refer to as the “wild-type” sequence. This fact has two negative consequences. First, since each wild-type molecule is one that can be mistakenly read as a mutation, the wild type is a ubiquitous source of false-positives. Second, since our goal is to detect mutations, most of the sequencing capacity and costs are devoted to sequence copies that are of little interest. Therefore, removing as many wild-type sequences as possible prior to DNA sequencing, *while measuring the extent of that removal*, would greatly improve both accuracy and sequencing efforts.

We present here a method, named MEMDS (Mutation Enrichment followed by upscaled Maximum Depth Sequencing), which uses principles of MDS for barcoding, as described above, but reaches a notably higher accuracy at a much smaller cost while focusing on detecting mutations in a very narrow ROI. MEMDS enriches the sample for mutations in the ROI prior to library preparation by removing a large fraction of non-mutated variants. In addition, it includes various steps that *a*) further enable the processing of the very large initial amounts of genomic DNA, as required for identifying *de novo* mutations in humans; *b*) enhance accuracy by using routinely and in an informed manner a dual barcoding system and other measures guarantying the authenticity of target DNA molecules; and *c*) accurately quantify the fraction of non-mutated variants removed, which is necessary in order to obtain the denominator for the calculation of *de novo* mutation rates. Using MEMDS, we achieve an error rate of at least 2.5 × 10*^−^*^9^ per base after removing the high-frequency G*→*T, C*→*T and C*→*A mutations (see supplementary section 8) and a recovery rate of *∼*35% of the input target sequences due to normal loss of material. With this recovery rate, for example, starting with 3 instances of a particular mutation in 300 million cells, 1 mutation in 100 million cells on average could be identified and reported. Thus, the recovery rate only affects the cost of sampling, and does not affect the cost or the accuracy of sequencing.

#### 1.3. The MEMDS method outline

The MEMDS method involves two workflows that are run in parallel. One enriches for mutations at the ROI—in this work using restriction enzyme digestion, though alternatives like CRISPR-editing (*18*) could also be used depending on the types of mutations that are sought after (point mutations, indels) and the improvement in site-recognition specificity (*19*). The other workflow is used for computing the enrichment fold, and hence the exact number of wild-type ROI sequences that were removed from the ROI pool. The protocol outlined below and in Figure S1 describes the workflow for the enrichment of mutated ROIs. This workflow is identical to the one applied for computing the enrichment fold, with the exception that the restriction enzyme used for enrichment (Fig. S1, step 1) is omitted in the latter. For a detailed explanation of the complete experimental design involving the two workflows, see supplementary section 2.

##### Step 1: Enzymatic digestion of genomic DNA

The genomic DNA is digested by two restriction enzymes. The first (RE-1) digests the wild-type sequence at a certain site that is several residues long and that constitutes the region of interest (ROI). Namely, the experiment is designed by choosing an ROI and an RE-1 so that the recognition site for RE-1 matches the wild-type sequence at the ROI. As a result, sequences with no mutations are efficiently digested, while variants with mutations that hamper site-recognition by RE-1 are protected from cleavage. These mutations are therefore enriched in the pool of uncleaved sequences (for calculating the exact number of wild-type ROIs that have been removed by RE-1, see the complete experimental design in supplementary section 2 and Fig. S2). The second restriction enzyme (RE-2) is used to cleave the DNA near the ROI. The choice of a suitable RE-2 is dependent on the availability of an adequate recognition site far enough from the RE-1 site to allow for an efficient annealing of a primary-barcode oligo (oligo A) between the two sites, yet short enough to meet the read-length limits of the chosen NGS sequencing platform. To satisfy these conditions, the RE-2 site may be selected to be either upstream or downstream of the ROI, a choice which will determine which of the two DNA strands will be barcoded and analyzed.

##### Step 2: Primary barcode attachment

Following digestion, the DNA is subjected to single-strand extension using a high-fidelity DNA polymerase and a single oligonucleotide (oligo A). Oligo A anneals with its 3’ part to the sequence between the RE-2 site and the RE-1 site and acts as a template for extension of the target-DNA strand. This extension reaction introduces three sequence features directly into the target strand: *a*) a segment of four bases that serves as a sample-identifier sequence to secure the sample in the event of a rare contamination by DNA libraries from other samples; *b*) 14 randomized bases that create a primary barcode unique to each particular DNA fragment; and *c*) an Illumina P5-primer sequence. In order to prevent the oligonucleotide itself from being extended while using an already barcoded target strand as template in the subsequent linear amplification step, an inverted-dT modification is included at the 3’ terminus of oligo A that blocks the DNA polymerase and prevents the extension of oligo A during the process. To account for the event that some oligo A molecules escaped the inverted-dT modification during their synthesis by the manufacturer, a single-base insertion is planted in the oligo A sequence that anneals to the genomic strand, so that undesired extensions of rare, unblocked oligos could be easily detected at the sequence analysis step for their inclusion of this single-base insertion, and removed.

##### Step 3: Linear amplification of barcoded ROI products

The genomic ROI is linearly amplified by 15 cycles using a high-fidelity DNA polymerase and a single primer (oligo B) that anneals to the Illumina P5-primer sequence. Oligo B contains the complete Illumina-adapter sequence, and carries five phosphorothioate bonds (PS) at its 5’ edge. This step results in up to 15 single-stranded copies of each barcoded ROI (each copy having been generated directly from the same barcoded original DNA molecule), protected by the phosphorothioate bonds from 5’-exonuclease activity.

##### Step 4: Degradation by 5’-exonucleases

The linear amplification products are treated with a mixture of 5’-exonucleases, which degrade both single and double-stranded DNA with or without phosphate groups at their 5’ termini, from the 5’ edge to the 3’ edge of each strand. The linearly-amplified ROI copies are protected from this exonuclease activity due to the multiple PS bonds at their 5’ edges. This step removes the majority of the genomic DNA, including most of the ROI digestion products, and simplifies the rest of the experimental workflow by allowing the next reactions to be carried in a small number of tubes rather than in 96-wells plates as well as by eliminating sequences that could potentially promote the generation of unwanted byproducts in the subsequent amplification steps.

##### Step 5: Secondary barcode attachment

The DNA from the 5’-exonuclease reaction is subjected to a single primer-extension reaction, using a secondary-barcode primer (oligo C) that anneals 3’ to the ROI site and extends by a single cycle using a high-fidelity polymerase. The secondary-barcode primer also carries three features: *a*) a segment of four bases that serves as a sample-identifier sequence; *b*) five randomized bases that create a secondary barcode generally unique to each member within a group of copies (copies sharing the same original DNA molecule); and *c*) an Illumina P7-primer sequence. This step produces a complementary strand for each of the 15 copies (or less) generated per target-DNA molecule during the linear amplification step. Each of these complementary strands carries the same primary-barcode sequence and a unique secondary-barcode sequence.

##### Step 6: Degradation by a 3’-exonuclease

To prevent recurrent labeling by secondary-barcode primers in subsequent amplification reactions, a 3’-exonuclease that degrades single-stranded DNA from the 3’ edge to the 5’ edge of the molecule is added immediately after the secondary barcode attachment to eliminate free, unbound primers. The double-stranded molecules that just completed the secondary barcode extension reaction are protected from this degradation. The 3’-exonuclease is added together with a known amount of relabeling-control primer (oligo D). This control primer is identical in sequence to the secondary-barcode primers except for the sample-identifier and the secondary-barcode features that are replaced by a known sequence. Therefore, in the event of incomplete degradation by the 3’-exonuclease, the amount of NGS reads with an oligo D sequence signature serves as a proxy for the frequency of relabeling by the secondary-barcode primer.

##### Step 7: Amplicon generation by PCR for next generation sequencing

PCR amplification of the purified DNA is carried using primers E and F, which add Illumina index and adapter sequences to the 3’ edge of the amplicon (as described in the materials and methods section, we break this step into two PCR reactions to preserve some of the first PCR product as a backup). Importantly, RE-1 digestion products that were not eliminated until this step will not be amplified, as only complete segments that were not digested by RE-1 have the two primer annealing sites.

##### Step 8: Analysis of sequenced data

NGS reads are grouped into families based on their primary-barcode sequences. Thus, each family is made of a collection of sequences originated from linearly amplified copies of a single target-DNA strand, belonging to a single gene. Each read in a family is aligned against a reference sequence specific to the donor and mutations with a high-quality sequencing score are noted. Three criteria are then used in combination to select for true mutations: *a*) the number of reads in the family (i.e., family size); *b*) the number of secondary barcodes associated with a particular mutation (i.e., BC2 count); and *c*), the fraction of the particular mutation in the family (i.e., mutation frequency). Mutation candidates that pass the combined cutoff criteria are designated true, *de novo* mutations. The total number of target wild-type sequences screened, which consist of *a*) target wild-type sequences that were digested by RE-1 and removed from the final DNA libraries, and *b*) target wild-type sequences that evaded RE-1 digestion and were included in the sequenced DNA libraries, is calculated from the sequencing outputs of the RE-1–treated and the RE-1–untreated samples (see supplementary section 2 and Fig. S2 for a detailed description of this methodology). Finally, from the mutation count and the total number of cells scanned, we calculate the per-locus, per-mutation *de novo* mutation rate for mutations of interest in the ROI.

### 2. Experimental design

#### 2.1 Calculating the number of RE-1 digested sequences

The evolutionarily relevant *de novo* mutation rate of a mutation in a sample is the number of target sequences in the sample identified as carrying that mutation divided by the total number of target sequences scanned by the MEMDS procedure. The number of target sequences scanned by the MEMDS procedure includes two sets of molecules: *a*) target sequences identified at the sequence analysis step; and *b*) target sequences removed by the enrichment step described in section 1.3 (Fig. S1, step 1). Therefore, to calculate the mutation rate, one must be able to determine how many target wild-type sequences have been removed by RE-1 digestion as opposed to having been removed by general loss of genetic material during the MEMDS procedure. The fold-reduction in target wild-type sequences, also referred to here as the RE-1 enrichment fold for RE-1–resistant mutations, multiplied by the number of target wild-type sequences identified at the sequence analysis step, yields the number of target sequences scanned by MEMDS. This number, in turn, serves as the denominator in the mutation rate calculation.

Toward this end, we have established an experimental design that uses the NGS output to obtain a precise measurement of the RE-1 enrichment fold. This experimental design avoids errors due to impreciseness of input DNA concentration measurements and variation in DNA loss and in performance of the MEMDS steps across samples.

We start with two tubes: a genomic- and a mock-DNA tube (Fig S2). The genomic-DNA tube includes the DNA extracted from the human sperm sample. In people who are not carriers of HbS or other mutations in the ROI, this tube contains mostly wild-type target sequences, which are sensitive to digestion by RE-1, and are denoted *S*. The other tube is a mock-DNA tube containing copies of an artificial sequence, denoted *R*, that are resistant to RE-1 digestion and easily distinguishable from natural mutants at the sequence analysis step. From the genomic-DNA tube, we transfer an amount of material into an “RE-1–treated” tube (Fig S2), whose material will undergo the full protocol including RE-1 digestion, and another amount into an “RE-1–untreated” tube, whose material will undergo the same steps except for digestion by RE-1. Likewise, from the mock-DNA tube, we transfer an amount of material into the RE-1–treated tube and another amount into the RE-1–untreated tube. The principle underlying this design is that the relative amounts transferred from a tube can be known through their volume measurements alone and, given these measurements, the RE-1 enrichment fold can be obtained by comparing the ratios between the numbers of sensitive and resistant DNA molecules following each treatment (as shown formally below), where those amounts are precisely known from the sequence analysis step.

Specifically, let the concentrations of *S* in the genomic-DNA tube and of *R* in the mock-DNA tube be [*S*] and [*R*], respectively. From the genomic-DNA tube we move a volume *V_Se_* to the “RE-1–treated” tube and a volume *V_Sc_* to the “RE-1–untreated” tube. From the mock-DNA tube we move a volume *V_Re_* to the RE-1–treated tube and a volume *V_Rc_* to the RE-1–untreated tube. Let *L_e_* represent the fold loss of material (whether sensitive or resistant) due to normal loss in the experimental condition, and *L_c_* represent the fold loss of material due to normal loss in the control condition. Finally, let *E* be the RE-1 enrichment factor (i.e., 1*/E* is the fold reduction in sensitive molecules in the RE-1–treated tube due to RE-1 digestion). At the final, sequence analysis step, we can precisely count the number of sensitive (i.e., wild-type) molecules called in the RE-1–treated condition, 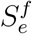; the number of artificial, resistant molecules called in the RE-1–treated condition, 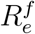; the number of sensitive (i.e., wild-type) molecules called in the RE-1–untreated condition, 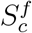; and the number of resistant molecules called in the RE-1–untreated condition, 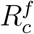. These quantities can be written as follows:

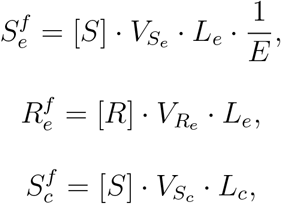

and

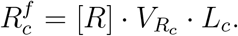

Therefore, we can obtain *E* by using the following formula:

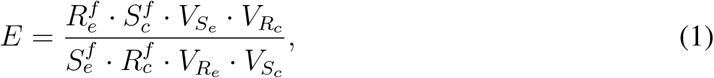

where all terms on the right-hand side are precisely known.

Given *E*, the amount of wild-type molecules scanned by the procedure, *W*, namely molecules either removed by RE-1 (and are therefore wild-type) or identified as wild-type at the sequence analysis step, is

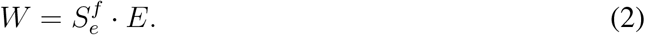

Thus, under the following assumptions, we do not need to know [*S*], [*R*] or the amount of material lost during the runs of the two parallel protocols in order to know the RE-1 enrichment fold: *a*) the solutions can be kept sufficiently homogeneous for the purposes of drawing volumes of similar concentrations from them (we ensure this by a thorough mixing before drawing); *b*) volumes, at the range used, can be measured easily and accurately (as is the case); *c*) the normal loss of material during the run of the protocol (i.e., loss that is not due to RE-1 digestion) within any one treatment of a sample (RE-1–treated or untreated) does not substantially differ between sensitive and resistant molecules (as we confirm by observation; see supplementary section 7).

Finally, suppose that mutations of *n* different types have been identified in the ROI at the sequence analysis step (mutations that confer resistance to digestion by RE-1). Let *M_i_* be the number of instances of mutation of type *i* ∈ { 1, 2, …, *n*} identified at that step. The rate of mutation *i* is then

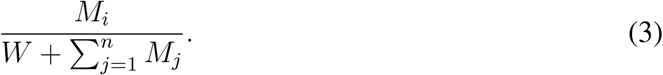

Since the sum in the denominator is negligible compared to *W*, it suffices to calculate the rate of mutation *i* as

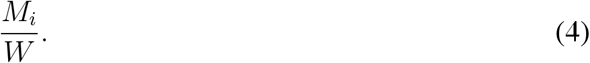

#### 2.2 Practical considerations of amounts used for the treated and untreated samples

If sequencing were costless and unlimited in capacity, one could have started the MEMDS processing of both the experimental and control samples with the same mix of genomic-DNA and mock-DNA sequences. Yet, while small amounts of mock-sequences could be easily identified in the RE-1–treated sample due to their enrichment, using the same amounts in the RE-1–untreated sample where no enrichment for RE-1– resistant variants is carried out would require a large sequencing effort to trace them among the vast majority of wild-type sequences. On the other hand, using large amounts of mock-DNA would improve their sequence recovery in the RE-1–untreated sample but would consume NGS capacity at the expense of sperm sample sequences in the RE-1–treated sample. Similarly, while large amounts of genomic DNA can be used in the RE-1–treated sample due to the removal of many wild-type ROIs by RE-1 from the final sequencing input, using the same amount of genomic DNA in the RE-1–untreated sample will require a massive sequencing effort.

Therefore, to match the experimental design to the NGS coverage limitations, we carry out the following routine. From a single human-sperm DNA source we transfer a volume equivalent to *∼*60–80 million haploid cells to the RE-1–treated tube, and a volume equivalent to exactly 5% of the initial amount taken for the RE-1–treated tube (i.e., *∼*3–4 million haploid cells, respectively) to the RE-1–untreated tube. For each ROI to be analyzed, we use a mix of two linearized plasmids as the mock-DNA sample. These plasmids carry all the ROI-flanking sequences that are necessary for processing by the MEMDS protocol, and each is designed to carry a unique stretch of mutations at the ROI that distinguishes it from the wild type, from natural mutants, and from the other plasmid (we use multiple mutations to make it practically impossible for the plasmid to be indistinguishable from natural mutants). Using the same plasmid-mix source tube, we add a volume equivalent to 7,500 copies from each linearized plasmid to the RE-1–treated tube (thus creating a genome:plasmid ratio close to 10,000:1) and 45,000 copies from each plasmid to the RE-1–untreated tube (creating a genome:plasmid ratio close to 100:1). Importantly, and as discussed above, the relative volumes drawn from any one source tube, not the absolute amounts of genomic and plasmid DNA, are the values that matter for the RE-1 enrichment fold calculation (Eq. 1).

### 3. *HBB* and *HBD* sequence features utilized by the MEMDS method

Different mutations in *HBB* that protect against malaria are known to have occurred and to have spread in human populations multiple times (*20, 21*). HbS, the most notable mutation variant associated with malarial resistance, is a single base substitution (20A*→*T) in codon 6 of the *HBB* coding sequence that causes a Glutamate to Valine change (*22–24*). Some other point mutations and short deletions near the HbS site are also known to confer malarial resistance (*25, 26*). *δ*-globin, encoded by the *HBD* gene, is expressed in adulthood together with *HBB* (*27*). These two paralogues exhibit a high degree of homology, showing 80% identity in coding sequence and 93% identity in amino acid sequence. However, mutations in *HBD* are not considered to be protective against malaria, probably due to its low expression levels compared to *HBB*, which accounts for less than 3% of the hemoglobin in adults (*28*).

The *HBB* and *HBD* gene sequences that were selected for processing by the MEMDS method encompass 114 bases from exon 1, ranging from 32 nucleotides upstream of the mRNA translation start site to 81 nucleotides into the protein coding sequence (Fig. S3). This region is highly conserved between the two genes, which differ in only eight of the 114 bases. The region of interest (ROI) is a palindromic sequence found between positions 16-22 of the coding sequence, which forms the recognition site for the restriction enzyme Bsu36I (CCTNAGG) both in *HBB* and *HBD*. Since Bsu36I can tolerate any of the four possible nucleotides at the central position of its recognition sequence, the ROI is limited to six of the seven nucleotides of this palindromic sequence. Therefore, Bsu36I serves as RE-1, which digest non-mutated (wild type) ROI sequences and enriches for *HBB*- and *HBD*-ROI mutation variants (Fig. S1, step 1). The second restriction enzyme, HpyCH4III, which serves as RE-2 for the primary-barcode attachment, digests the *HBB* and *HBD* gene segments at its recognition site (ACNGT), 45 bases upstream of the 5’ edge of the Bsu36I restriction site. The identity of the “N” base at the center of the HpyCH4III site is of central importance, as after digestion by HpyCH4III this base is found at the 3’ terminus of the antisense strand that extends to incorporate the primary barcode via a fill-in reaction (Fig. S1, step 2). Since *HBB* and *HBD* carry a different nucleotide at this “N” position of the HpyCH4III recognition site, the primary-barcode oligo (oligo A) that initiates the fill-in reaction carried a randomized base at that position, matching either one of the two complementary bases to allow for similar efficiencies of primary-barcode synthesis for the two genes.

A region of 30 bases between the Bsu36I and the HpyCH4III sites is used as the annealing site for the primary-barcode oligo, and a region of 28 bases starting 60 bases downstream to the 3’ edge of the Bsu36I restriction site serves as the annealing site for the secondary-barcode primer (oligo C, Fig. S1, step 5). This leaves a sequence of 15 bases upstream of the ROI site and 32 bases downstream of this site that are untouched by any primer and are amplified together with the enriched ROI elements. We use the differences between the *HBB* and *HBD* ROI 3’-flanking sequences to define NGS reads as belonging to either *HBB* or *HBD* during the sequence analysis step.

An advantage of the fact that, for each donor, the *HBB* and *HBD* ROIs are processed by MEMDS simultaneously and side by side in the same reaction by the same oligos, with the consequence that the genes are only separated by their unique and small sequence differences at the computational stage, is that any mutational patterns arising in one gene and not in the other cannot be assigned to methodological artifacts. Such artifacts would have been expected to manifest themselves in both genes.

### 4. *In vitro* analysis of the effects of restriction-site mutations on Bsu36I activity

To study the *de novo* origination rate of the HbS mutation in *HBB* and of the parallel A*→*T mutation in *HBD*, as well as the rates of other mutations in their vicinity, we applied the MEMDS method to the *HBB* and *HBD* genes from human-sperm DNA, exploiting the fact that codon 6 in both genes comprises a part of the recognition site for the Bsu36I restriction enzyme (RE-1) (Fig. S3). However, the enrichment of any mutation within the ROI site depends on the efficient blockage of Bsu36I digestion, and therefore we tested the ability of all single-base substitutions to effectively block Bsu36I digestion. For this purpose, we have applied the deep mutational scanning approach (*29, 30*) to generate a synthetic-DNA library of *HBB* segments carrying all possible single-base substitutions in the Bsu36I site and its flanking sequences. After incubating the library for 20 hours either with or without Bsu36I, next-generation sequencing of the full-length products that were recovered from each sample allowed us to count each mutation variant and to calculate the fold difference in its frequency between the two samples, which serves as a proxy to the degree of resistance to Bsu36I cleavage. In accordance with the known consensus sequence for the Bsu36I site (CCTNAGG), we found that while the central base can tolerate any type of substitution, any single point mutation in the remaining 6 bases of the Bsu36I site is resistant to digestion (Fig. S4). The degree of resistance is similar to that of a variant that carries substitutions in all of the seven bases that constitute the Bsu36I site (the same set of mutations found in ALP13, which is one of the two artificial ROIs used to determine the Bsu36I-enrichment factor). Therefore, natural single-base substitutions in Bsu36I sites are effective substrates for enrichment by Bsu36I.

### 5. Generating *HBB* and *HBD* sequence datasets

We applied the MEMDS protocol to 7 sperm-DNA samples from donors of African ancestry (AFR1-7) and four samples from donors of European ancestry (EUR1-4) (see Table S1 for detail). As described in supplementary section 2, from each sample we aliquoted genomic DNA in an amount equivalent to 60-80 million sperm cells into one tube (referred to as “Bsu36I-treated”) and an amount equivalent to 5% of the cells (3-4 million sperm cells, respectively) into a second tube (referred to as “Bsu36I-untreated”). Each of the two reaction tubes was supplemented by a known amount of plasmid mixture that carries artificial Bsu36I-resistant *HBB* and *HBD* sequences. The Bsu36I-treated sample was treated with Bsu36I and HpyCH4III, and the Bsu36I-untreated sample was treated with HpyCH4III only. With the exception of the digestion step, the two samples were processed identically by the complete MEMDS procedure and sequenced.

Following standard quality filtration and merging of overlapping paired-end reads, reads were validated for carrying the 14-mer primary-barcode and the 5-mer secondary-barcode features, as well as the unique 5’ and 3’ sample-identifier sequences.

Control-guanine insertions designed to report for primary-barcode indirect labeling (see Fig S1 step 2, and supplementary section 13 for oligo A features) were found to be present in *∼*1/9,000 reads for the Bsu36I-treated samples and *∼*1/28,000 reads for the Bsu36I-untreated samples (Figure S5A), implying an efficient 3’inverted-dT blockage of the primary-barcode oligo. Yet, the observed difference in the fraction of reads with control-guanine insertions between the Bsu36I-treated and untreated samples suggests that the large amount of treated DNA in the former (leading, for example, to longer preparation times for some of the MEMDS step) and/or residual effects of Bsu36I digestion products may account for the elevated frequencies of indirect-labeling in the former.

After removing sequences with the control-guanine insertions, reads were sorted into separate *HBB* and *HBD* datasets based on their match to unique sequence features of each gene (see Materials and Methods and Figure S3 for the exact sorting parameters). Consequently, each sperm sample produced four major datasets consisting of separate *HBB* and *HBD* sequencing pools for each of the Bsu36I-treated and untreated samples. Each read was then aligned against the donor’s reference sequence and the presence of mutations and their types were noted per position. Next, reads were grouped into families based on their primary barcode sequences, where within each family, reads shared the same primary barcode and represented multiple copies of the same original target-DNA molecule, and each secondary barcode represented one of the *≤*15 linearly amplified copies of that target molecule. Only families that passed the criteria discussed in the next section were selected for mutation-detection analysis.

### 6. Filtering families for mutation detection analysis

Three major parameters affect the level of accuracy by which a primary-barcode family is considered as being originated from either a wild-type or a mutated target DNA molecule: *a*) the number of reads belonging to a primary-barcode family (i.e. family size); *b*) the fraction of reads in the family having the same nucleotide (either a wild type or a mutation) in a given position (mutation frequency); and *c*) the number of secondary barcodes in a primary-barcode family associated with either a wild-type base or a particular mutation (i.e., BC2 count).

For all donors and treatments, most primary-barcode families contained multiple reads (Fig. S6). Yet, as previously reported for the MDS method (*16*), many families were represented by single reads. It is likely that many of these single-read families represent genuine labeling events that did not accumulate enough reads during the amplification steps, and are thus excluded from analysis and are a part of the general loss of material. Additionally, we found that between 20% and more than 50% of the single-read families had a primary-barcode sequence that deviated by a Hamming distance of one from one of the primary-barcode sequences of a family with multiple reads (Fig. S6), suggesting that each of these single-read families is likely the result of a single-base error in the primary barcode of one of the reads in a multiple-reads family acquired during library preparation or NGS steps. Supporting this inference, a far smaller percentage of the primary barcodes of families with multiple reads were found to be at a Hamming distance of one away from other primary-barcode families.

In addition to eliminating the barcode-error artifact, increasing the family size reduces the influence that sequence errors have on the final consensus sequence. Under the most stringent assumption that all the mutations appearing in sequences from the Bsu36I-untreated samples and in the ROI-flanking sequences from the Bsu36I-treated samples are due to NGS or PCR errors, gradually increasing the family-size cutoff reduced the acceptance rate of these false-positive mutations for both samples (Fig. S7A). We have selected a minimum required family size of four reads for further mutation detection analysis, as increasing the family size cutoff beyond 4 reads did not noticeably improve mutation detection accuracy but continued to reduce the number of recovered families (Fig. S7A).

Increasing the mutation frequency cutoff, i.e., the minimal fraction of reads in a family carrying a particular nucleotide in a particular position allowing us to accept that nucleotide, reduced the fraction of false positive mutant families already when using low cutoff values, suggesting that the source of these mutations are late PCR errors or NGS errors that appear in small fractions within families (Fig. S7B). We have selected a mutation-frequency cutoff of 0.7 (i.e., at least 70% of the family members carried either a wild-type base or a particular mutation at a given position), which provided a good balance between the number of mutations that were filtered out and the number of recovered families.

For each family, the number of unique secondary barcodes that were added after the linear amplification step and before the PCR amplification step corresponds to the number of unique linearly amplified copies of the original DNA molecule. Therefore, requiring multiple secondary barcodes allows us to reduce the error rate by ensuring that reads from distinct linear amplification events are used in the analysis. For the families with the highest read counts, we found that usually 4-5 of the unique secondary barcodes were more frequent than the remaining secondary barcodes, suggesting that while some of the linearly amplified copies of each ROI were PCR amplified more efficiently than others their repertoire was diverse enough and not over dominated by a single linearly amplified copy (Fig. S8). We found negligible amounts of families with more than 15 unique secondary barcodes, which matches the maximal number of linearly amplified copies and supports the authenticity of these barcodes. Our control for secondary-barcode relabeling suggests that such an event occurs once every 250-350 reads (Figure S5B). Since both the originally labeled and the erroneously relabeled copies need to be sampled and included in the same family for their secondary barcodes to be miscounted twice, the negative effect of this event should be even smaller. Limiting mutation calling by requiring a minimum of two secondary barcodes associated with a particular nucleotide in a particular position as a condition for that nucleotide to be accepted (whether it is a wild type or a mutation), in addition to the family size cutoff, improved accuracy with a minimal effect on the number of recovered families (Fig. S7C). Thus, besides the major contribution of mutation enrichment by restriction-enzyme digestion to mutation detection accuracy, setting up a secondary-barcode count cutoff as a regular part of the MEMDS procedure add further precision in mutation calling in comparison to the MDS method (*16*).

Based on the above considerations we have selected the following combined threshold criteria: primary-barcode families with at least four reads, a minimal within-family mutation-frequency cutoff of 70%, and the association of at least two secondary barcodes with each base. The flowchart of the algorithm we developed for base calling is provided in Figure S9. Importantly, we use the same criteria for calling a wild-type family or a mutant family, thus eliminating any computational bias that would have been associated with different treatments of wild type and mutant and could affect the calculation of *de novo* mutation rates. In those rare events where neither the wild-type nucleotide nor a particular mutation in a *≥*4-read family meet the mutation-frequency cutoff or the secondary-barcode count cutoff conditions at a certain position, the nucleotide identity at that position is declared ambiguous and the family is rejected from further analysis.

The numbers of families that were rejected or approved by these three cutoff criteria are shown for each library in Table S2. We also removed from further analysis *HBB* and *HBD* families that shared the same primary-barcode sequences, which point to *HBB*/*HBD* chimeric artifacts that were generated during library amplification (we discuss these events more thoroughly in supplementary section 9). Datasheets S1–S44 describe the properties of each primary-barcode family that passed the combined cutoff criteria.

### 7. MEMDS performance measures: Enrichment factors, numbers of genomes scanned, mutation recovery rate and error rate

In order to determine the origination rate of a particular mutation at a particular site, one must divide the number of sampled target sequences carrying that mutation by the total number of sampled target sequences. For the Bsu36I-untreated samples, the total number of target sequences sampled is derived solely from the number of families that are present in the sequencing output and that have passed the combined cutoff criteria. For the Bsu36I-treated samples, however, the total number of target sequences sampled (the number of genomes scanned by MEMDS) must include also the number of target sequences that have been eliminated due to Bsu36I digestion. We derive this number using the method described in supplementary section 2. To recapitulate, we divide the ratio between the number of artificial Bsu36I-resistant families and the number of wild-type families that result from applying the MEMDS procedure to the input mixture of the Bsu36I-treated sample by the analogous ratio from the untreated sample, while correcting for the different volumes drawn for practical considerations from different source tubes, to obtain the Bsu36I-enrichment factor. The number of scanned wild-type target sequences (i.e., the number of target sequences that had been removed by Bsu36I digestion plus the number of target sequences that escaped Bsu36I digestion and formed wild-type families that passed the cutoff criteria) is then calculated by multiplying the number of wild-type families that passed the cutoff criteria by the Bsu36I-enrichment factor.

On average, about 13% of the input *HBB* and *HBD* wild-type ROIs were recovered in the Bsu36I-untreated samples (Fig. S10). A similar recovery rate of 15% was observed for the artificial ROIs in these samples, suggesting that both the genomic and the plasmid variants are processed similarly by this MEMDS workflow. Notably, the relatively low recovery rate of both target molecules is likely due to the overload input DNA, as no restriction enzyme-based depletion of wild-type ROI sequences takes place in these samples. However, these recovery rates satisfy the main purpose of the Bsu36I-untreated samples, which is to set the artificial ROI/genomic ROI ratio in the absence of Bsu36I enrichment (see supplementary section 2).

Following Bsu36I treatment, the wild-type ROI levels dropped to an average of *∼*0.25% of their input levels, while the artificial ROI levels were at an average of *∼*40% of their input levels, with *HBB* recovery being slightly higher than the recovery of *HBD* (Fig. S10). Given that point mutations at the ROI block Bsu36I digestion and are enriched similarly to the artificial sequences (Fig. S4), this percent recovery of the artificial ROIs suggests that at least a third of Bsu36I-resistant mutations that were present in the input sperm DNA were recovered in the Bsu36I-treated samples by the MEMDS procedure.

For each donor, the Bsu36I enrichment factors obtained from the artificial ROI/genomic ROI ratios showed a high degree of consistency between *HBB* and *HBD*, which reflects a similar activity of Bsu36I on both genes (Table S3). However, these enrichment factors displayed some variation across donors, ranging from a 64-fold enrichment to a 340-fold enrichment, likely associated with either differences in Bsu36I activity due to batch effects or differences in the integrity of the sperm DNA (namely, the fraction of *HBB* and *HBD* ROI segments that were in a double-stranded state). In particular, the average enrichment factor of the four European samples (278.0 *±*47.8) was about 2.6-fold higher than the average enrichment factor of the seven African samples (107.3 *±*35.0). This difference in the average enrichment factor, which represents a Bsu36I digestion of 99.6% of the of the European *HBB* and *HBD* ROI sequences compared to 99.1% of the African ROI sequences, could arise due to the differences in semen composition between the two groups of donors that affect the double-strand state of the genomic DNA during its extraction. We found that the enrichments of the highly frequent G*→*T and C*→*T substitutions in the target DNA strand (C*→*A and G*→*A in the sequenced strand, respectively) at the ROIs exhibit differences between the eleven donors that followed the same direction as the differences between the Bsu36I-enrichment factors, which further supports the Bsu36I-enrichment-factor calculation by our approach (see supplementary section 8). Thus, given an average enrichment factor of *∼*170-fold, the enrichment step of MEMDS alone boosts both the sequence coverage in the search for the target *de novo* mutations and the accuracy of mutation-detection by more than two orders of magnitude in comparison to the mutation rate in the Bsu36I-untreated samples.

Based on the calculated enrichment factors, the total number of wild-type *HBB* and *HBD* target sequences that were screened by Bsu36I (i.e. in the Bsu36I-treated samples) reaches about 300 million for each gene (Table S3). These numbers represent an average recovery rate of slightly more than 35% for wild-type target sequences, which is highly similar to the recovery rate of the artificial ROIs, further supporting the similar processing of both the plasmid and the genomic variants by the MEMDS procedure.

To calculate the MEMDS error rate, all G*→*T, C*→*T and C*→*A mutations in the target-DNA strand were removed from analysis due to the reasons discussed in supplementary section Under the most stringent assumption that all remaining mutations found at the ROI and ROI-flanking sequences in the Bsu36I-untreated samples and at the ROI-flanking sequence in the Bsu36I-treated samples arose due to PCR or NGS errors, the total per-base error rates for the Bsu36I-untreated and treated samples across all genes and donors in these unenriched sequences was 1.3 × 10*^−^*^7^ and 3.1 × 10*^−^*^7^ per base, respectively (Fig. S11A) (error rate calculation for an individual gene from a single donor would be less accurate due to low counts of non G*→*T, C*→*T and C*→*A mutations in these sequences). This 2.4-fold difference between the two error rates could result from a secondary influence of the large amount of treated DNA in the former, residual effects of Bsu36I-digestion products, or from moderate enrichment of mutations outside the Bsu36I-recognition site that may display weak inhibitory effects on Bsu36I. Taking a conservative approach, we have selected the error rate at the ROI-flanking sequences in the Bsu36I-treated samples as our base line for the calculation of the MEMDS error rate. Therefore, to determine the MEMDS error rate, namely the error rate within the 6 bases of the ROI site in the Bsu36I-treated samples, for each donor and each of the two genes, we divided the error rate obtained for the ROI-flanking sequences of *HBB* and *HBD* in the Bsu36I-treated samples (2.9 × 10*^−^*^7^ and 3.3 × 10*^−^*^7^, respectively) by their matching enrichment factors, reaching an average per-base error rate of 2.3 × 10*^−^*^9^ (*±*1.2 × 10*^−^*^9^) and 2.6 × 10*^−^*^9^ (*±*1.4 × 10*^−^*^9^) for *HBB* and *HBD*, respectively, and an average of 2.5 × 10*^−^*^9^ (*±*1.3 × 10*^−^*^9^) for both genes (Fig. S11B). Together with MEMDS’s substantial reduction in sequencing cost, this error rate enables the identification of specific *de-novo* mutations at particular bases of interest that originate at rates even lower than the whole genome average mutation rate in humans.

### 8. G***→***T, C***→***T and C***→***A mutations

With respect to the ROI-flanking sequences both the Bsu36I-treated and the Bsu36I-untreated samples displayed similar mutation patterns (Figures S12 and S13) with Pearson’s correlation coefficients of 0.934 for *HBB* and 0.878 for *HBD*, for the mutations depicted in Figure S13. Inspecting this mutational spectrum revealed high frequencies of single-base substitutions of three types, two of which were C*→*A and G*→*A, with average rates of *∼*2.4 × 10*^−^*^6^ (*±*1.4 × 10*^−^*^6^) and *∼*4.2 × 10*^−^*^6^ (*±*2.2 × 10*^−^*^9^), resp., across both genes, treatments and donors. Since the consensus sequences are composed of reads of *HBB* and *HBD* at the sense orientation, these mutations are the reciprocals of the G*→*T and C*→*T mutations, respectively, that were present in the target, antisense DNA strand.

A major cause of G*→*T mutations is DNA damage occurring both endogenously under normal metabolic conditions (*31, 32*) and during DNA extraction and NGS preparation procedures (*33–35*). Reactive oxygen species (ROS) that arise as by-products of normal aerobic metabolism or due to the high temperatures used during DNA purification and PCR amplification steps can damage the genomic DNA by oxidizing guanine to 8-oxoguanine (8-oxoG), which in turn can pair up with adenine (8-oxoG:A) and promote a G:C*→*T:A mutation (*36*). C*→*T mutations occur either naturally or *in vitro* by heat-induced hydrolytic deamination of either cytosine or 5-methylcytosine (5-meC) that generate uracil or thymine, respectively (*37,38*). These bases can then pair up with adenine and facilitate a C:G*→*T:A transition (*39*).

In the *HBB* and *HBD* target antisense strands, G*→*T substitutions constituted *∼*27% and *∼*35% of the mutations found across Bsu36I-untreated and treated samples, respectively, and C*→*T substitutions were *∼*67% and *∼*56% of the mutations across the same samples, resp. Compared to these high rates, we found the rates of the reciprocal substitutions in the same strands to be much lower: 4% and 6 for C*→*A, and less than 0.5% for G*→*A for each treatment (Figure S12). As in previous studies that used one of the two DNA strands to explore *de novo* mutations, we take these imbalanced frequencies to indicate that the formation of 8-oxoG and deaminated cytosines (or 5-meC) occurs in the DNA either in vivo or *in vitro* before the library amplification step, while the subsequent completion to full G:C*→*T:A and C:G*→*T:A substitutions, respectively, occurs during library amplification and not before then (*13, 33, 34*). Such DNA damages occurring *in vitro* before the library amplification step and/or representing the snapshot image of a disrupted ongoing process of base-damage and repair *in vivo* could result in mutational reads only when they occur in the target, antisense strand and not when they occur in the sequenced, sense strand, explaining the target-strand imbalance mentioned above.

Examining the mutation distribution along *HBB* and *HBD* sequences across all donors and treatments (Fig. S13) reveals that both the G*→*T and C*→*T substitutions in the target, antisense strand were enriched at the ROI site, suggesting that both these types of DNA damages were formed at the target strand either *in vivo* or *in vitro* prior to the enzymatic digestion step (Fig. S1, step 1) and conferred Bsu36I resistance. Indeed, 8-oxoG modifications placed at restriction sites or near them have been shown to interfere with the activity of multiple restriction enzymes (*40–44*). Similarly, generation of T:G or U:G mismatches that result from the deamination of cytosine or 5-meC have also been shown to inhibit enzymatic digestion (*45, 46*).

Importantly, while the frequency of the C*→*A mutation (the reciprocal of G*→*T) in the target strand was much lower than that of the G*→*T mutation, it was still noticeably higher than those of all other point mutations besides G*→*T and C*→*T, with an average rate of *∼*3.9 × 10*^−^*^7^ (Fig. S12). This observation implies that some of the guanine-oxidative damages could have affected the DNA sense strand (the non-target strand) early enough during library amplification and thus were able to produce mutations that were approved by the combined cutoff criteria.

The high frequency of the G*→*T and C*→*T substitutions in the target, antisense strand at the ROI site in the Bsu36I-treated and untreated samples allowed us to calculate their enrichment fold in a manner entirely independent from the enrichment-fold calculation based on the artificial sequences described in supplementary section 2 (albeit more limited and less accurate than the latter as these mutations were either too infrequent or absent from the ROI of every Bsu36I-untreated sample). We found the enrichment of these substitutions to follow the same trend as the Bsu36I-enrichment factors, i.e., samples with higher enrichment factor values calculated from the artificial sequences showed increased G*→*T and C*→*T enrichments at the ROI site in comparison to samples with lower enrichment factor values (Fig. S14). In absolute terms, G*→*T and C*→*T enrichment values were lower than their matching enrichment factors calculated from the artificial sequences, likely due to 8-oxoG damages providing only incomplete resistance to Bsu36I digestion (*44*) and/or continuous DNA damage occurring after the restriction enzyme digestion and affecting uncut segments before the linear amplification step in both the Bsu36I-treated and untreated samples.

In addition to the enrichment of G*→*T and C*→*T mutations at the ROI site, we found also a G*→*T enriched mutation at position 14 of *HBB* and *HBD*, two residues away from the Bsu36I site (Fig. S13). Indeed, 8-oxoG has been shown to affect neighboring bases and to compromise enzymatic digestion when placed near a restriction site (*44, 47*). Our finding that a complete G:C*→*T:A mutation (i.e., the mutation is fixed in both strands) at position 14 has no effect on Bsu36I digestion (Fig. S4) further supports the effect of a single-stranded change such as 8-oxoG on Bsu36I digestion.

Given their high frequencies and unbalanced distribution between the two strands that disqualify G*→*T, C*→*T and C*→*A substitutions in the target DNA strand as true, *de novo* mutations occurring in sperm cells, we excluded them (i.e., their sequencing output reciprocals C*→*A, G*→*A and G*→*T) from the calculation of mutation rates. By comparison, Jee et al. (*16*) removed the mutations C*→*T, C*→*A and A*→*G from the MDS analysis of bacterial gene segments, suggesting that their high frequencies and strand occupancy bias reflect a snapshot image of base misincorporation and repair processes in the bacterial cells (*16*).

### 9. Repeated *de novo* mutations are not due to chimeric duplication events

As shown in Table 1 of the main text, certain mutations were found to be enriched in the ROI of both *HBB* and *HBD* of the four donors by the Bsu36I treatment and are considered as true, *de novo* mutations. Of the overall 49 single-base substitutions that were found for all genes and donors, 14 occurred repeatedly in the same gene and in the same donor. Of the six enriched deletion mutations that were found, the Hb-Leiden mutation (a deletion of either codon 6 or codon 7, which results in the same sequence) occurred repeatedly in *HBB* in seven of the 11 samples and in *HBD* in two of the 11 samples. Methodological artifacts cannot explain the correspondence that we see between de novo origination rates and observations of alleles in populations, as described in the main text and section 11 SI Appendix. That being said, we additionally confirmed independently that these repetitions are not due to duplications of mutant families by artifactual chimera that are generated during library preparation.

Chimeric sequences arising during PCR amplification are a common source of NGS sequencing artifacts, ranging from a few percent to nearly half of the sequences in individual libraries (*48–50*). A chimeric sequence can be generated during PCR due to low processivity of the DNA polymerase or insufficient elongation time that produce an incomplete DNA strand. Such a strand can anneal in one of the following cycles to a full-length strand of a second allele or a paralogue gene and complete its extension, thus creating an Allele1/Allele2, or a Gene1/Gene2 chimeric product in addition to the PCR products of the two alleles, or genes, respectively. Therefore, a similar mechanism involving the interaction between an incomplete strand of a mutation variant and a full-length strand of a wild-type variant could theoretically result in the duplication of the mutation variant. Specifically, if a mutation-carrying strand ends prematurely and loses its primary barcode, when serving as a mega primer in the following PCR cycle it will acquire a new barcode and could potentially lead to a second family that carries the mutation, where, in the unlikely event that such a family passes the combined filtration criteria, that second family is a false positive.

Since *HBB*/*HBD* chimeras in our experiment are identifiable, as they carry both *HBB*- and *HBD*-specific markers on different sides of the chimeric breakpoint (exemplified, for instance, by the relatively high frequency of *HBB* 9C*→*T or *HBD* 9T*→*C mutations), we used the *HBB*/*HBD* chimeras to estimate the probability that two separate families (each with its own primary barcode) that carry the same mutation actually arose from one family due to a chimeric event and thus represent a double-counting of the mutation. Specifically, since *HBB* and *HBD* share a high sequence identity, *HBB*/*HBD* chimeric artifacts could be generated as explained, by extending an incomplete strand of one paralog while using the full-length strand of the other paralog as a template during library preparation. Thus the extended strand acquires the primary barcode of the template strand. As *HBB* and *HBD* reads are sorted into distinct sequence analysis pools based on their unique sequence markers, both the chimeric family and the “template” family were identified by their shared primary-barcode sequences and removed from further analysis (supplementary section 6). For AFR1, AFR2 and AFR3, who exhibited multiple instances of the HbS mutation, about 1% or less of the Bsu36I-treated families that passed our combined filtration criteria were identified as potential *HBB*/*HBD* chimeras (Table S2). Since each chimeric event between a mutant strand and a wild-type strand can result in either a wild type or a mutant duplication, the fraction of observed mutant families that constitutes artifactual duplicates of other mutant families during PCR is at most half the fraction of chimerism, namely *<* 0.5%. This per-mutant probability of artificial duplication is unable to account for the recurrence of mutations in our data, as not a single mutation is expected in the dataset to be a false positive due to double counting. In particular, it is unable to explain the repetition of HbS and Hb-Leiden in the data.

### 10. Genome-wide average point mutation, indel and A***→***T rates

The human genome-wide point mutation rate per base per generation is generally considered to be close to 1 × 10*^−^*^8^ (*51*). Most recent estimates fall within the range 1–1.5 × 10*^−^*^8^ (*52–56*). Thus, we use the midpoint of this range, 1.25 × 10*^−^*^8^, as a reference point for the sake of comparisons. Studies of the whole-exome mutation rate per base per generation average a bit higher, around 1.5 × 10*^−^*^8^ (*57*). However, while many of these studies are based on individuals with a given disease, whole-exome mutation rates from healthy individuals or neutral sites have reported rates closer to the 1.25 × 10*^−^*^8^ reference point (*53, 57*). Either way, whether using 1.25 or the relatively high 1.5 as a reference point, no significance assignment reported in the main text is affected. Furthermore, since the human genome-wide per base per generation indel rate is more than 10× smaller than the human genome-wide per base per generation point mutation rate (*58–60*), we use 1.25 × 10*^−^*^9^ as a slightly conservative reference point for the latter.

To obtain a per-base point mutation rate across an ROI that can be compared to previous measures of the genome-wide average per-base point mutation rate, we take into account the fact that 12 out of 18 of the possible point mutations across a single instance of the ROI are observable due to G*→*T, C*→*T and C*→*A exclusion. The effective average per-base point mutation rate observed, *µ*^ROI^, is then obtained as follows:

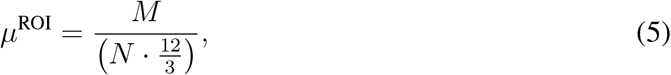

where *M* is the total number of point mutations observed across the ROI and *N* is the total number of families analyzed, namely the primary barcode families that have passed the combined cutoff criteria. In other words, the total number of point mutations observed is divided by the maximal number of point mutations that could have been observed, where the latter is divided by 3 to obtain a per-base rate that can be compared to previous measures of the genome-wide point mutation rate per-base, since 3 mutations are possible per base. This simple calculation is suitable for the purpose of testing whether the average point mutation rates observed across the ROIs are higher than previously measured genome-wide point mutation rates, because here the ROI rate is inferred from 9 out of 12 possible point mutation types, where the average rate of the excluded mutation types is expected *a priori* to be no lower than the average rate of the included ones based on previous knowledge of mutation rates per type (*55, 61*).

The advantage of this method of comparing the ROI average to the genome-wide average per-base point mutation rate is that it takes into account the particular sequence at the ROI. Alternatively, taking the ROI per-base point mutation rate that is due to the 9 observable point mutation types only and comparing it to its genome-wide equivalent would require a complex adjustment of the genome-wide measure, whereas the goal here is merely to provide a general-sense comparison to a well known figure.

To obtain a per-base indel rate across an ROI, we divide the total number of indel events (in our samples, only deletions have been observed) by the total number of bases examined across primary barcode families by MEMDS. Here, a complication arises from the fact that we can observe not only indels that are entirely contained within the ROI but also indels that partly overlap with the ROI, as those too are captured and enriched by MEMDS. A simple way of addressing this fact is to expand the number of base positions examined to include all positions between the farthest upstream and downstream breakpoints observed in the dataset, namely between position 14 and 24 (a stretch of 11 positions). The denominator of the indel rate calculation is then the total number of families observed multiplied by 11. For testing whether the observed indel rate is higher than expected from the genome-wide average, this simple method is slightly conservative, because indels that overlap with the region between positions 14 and 24 but not with the ROI (e.g., a 12 14 deletion) are potentially possible but not observable and do not contribute toward the indel count.

To explain the calculation of the genome-wide average rates for the observable point-mutation types, take the A*→*T transversion for example. Based on a subset of *de novo* mutations with phasing information, the A:T*→*T:A transversion accounts for *∼*6.5% of the total of point mutations across the human genome in males (*55*), while the A:T content across the human genome is *∼*59% (*62*). Therefore, the average A*→*T mutation rate per adenine base per generation in males can be estimated as follows:

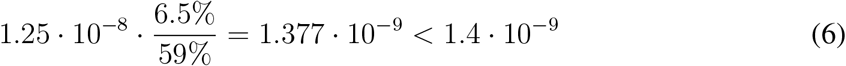

Using a similar calculation with data on the relative frequencies of Extremely Rare Variants (ERVs) (*61*) allows us to obtain the A*→*T mutation rates in the 3-mer, 5-mer and 7-mer contexts relevant to the *HBB* and *HBD* ROIs, namely the GAG, TGAGG and CTGAGGA contexts, respectively (using the supporting materials of ref. 61), which are approximately 1.3, 1.2 and 0.9 ×10*^−^*^9^, respectively. In the same way, we obtain the genome-wide average rates, with or without the extended contexts, for the other observable point-mutations (Table S4). Thus, the local sequence up to the 7-mer context does not explain the high *de novo* origination rate of the A*→*T mutation in the *HBB* ROI in the African samples, consistent with the implication of complex genetic influences on the mutation-specific mutation origination rate (see main text). The same is true for the other point mutations that deviate from their expected genome-wide average rates (Table S4).

### 11. The correspondence between *de novo* rates and observations of alleles in populations

12 point mutation types in each ROI are observable by our method, all of which have been observed to occur *de novo* in at least one ethnicity and ROI. In addition, 23 deletions of up to size 3 are observable in each ROI (taking into account that size 2 and 3 observable deletions reach beyond the boundaries of the ROI, and that size 1 deletions at position 19 are not observable).

The expected rate of indels decreases with indel size (*59*). Thus, because of the rarity of indels size *>*3, capping the analysis at this size provides for conservative *P* values, as including more indels would have only increased the fraction of indels that both have not been reported on HbVar and that are not observed *de novo*. (Indeed, the next deletion reported on HbVar that could have been observed by our method, 20 45del, is of size 26). Insertions are also relatively rare and neither were reported before in the ROIs in HbVar nor are observed here *de novo*, and thus their exclusion is also conservative for our statistical tests.

Of the point mutations, 8 have been reported on HbVar (16*→*CG, 16C*→*T, 17C*→*G, 17C*→*T, 20A*→*G, 20A*→*T, 20A*→*C, and 22G*→*C), and of the 23 observable indels of up to size 3, 5 have been reported on HbVar: 16delC, 17 18delCT, 18 19delTG, 19 21delGAG or the equivalent 22 24delGAG (the Hb-Leiden mutation), and 20delA, all in *HBB* (Table S4). HbVar is an online database that gathers reports of all human hemoglobin variants from the literature and is arguably the largest source of information on this topic (*25, 26*).

The *de novo* rates of deletion types reported on HbVar are significantly higher than the rates of deletion types not reported on HbVar, both combining the two ROIs (*P* = 0.0033, two-sided permutation test) and for the *HBB* (*P* = 0.029) and *HBD* ROIs separately (*P* = 0.0056). However, because the Hb-Leiden mutation has an exceptionally high *de novo* rate among the deletions, in order to find out whether this mirroring between deletion *de novo* rates and reports of these mutations on HbVar is driven by the Hb-Leiden mutation alone or extends also to other mutations, we compare the fraction of deletion types reported on HbVar that have been observed *de novo* at least once in our data to the same fraction among deletion types not reported on HbVar. The former fraction is significantly larger than the latter both when combining the two ROIs (*P* = 0.0078, two-sided Fisher exact test) and for each ROI separately (*HBB*: *P* = 0.048, *HBD*: *P* = 0.0056, two-sided Fisher exact test). While these tests already show that the results are not driven only by the high Hb-Leiden *de novo* rate, when combining the two ROIs due to the similarity of indel types observed between them, the results remain significant also when excluding the Hb-Leiden mutation completely (*P* = 0.026 and *P* = 0.024, two-sided Fisher exact test), further demonstrating that the mirroring between these mutation’s *de novo* rates and observations of them in populations extends beyond the Hb-Leiden mutation. Indeed, note that of the 6 indel types observed by us *de novo* in any ROI and ethnicity, 4 are included in the 5 deletions reported on HbVar (16delC, 17 18delCT, 19 21delGAG or 22 24delGAG, and 20delA), and of the deletions reported on HbVar, only one has not been observed *de novo* (18 19delTG). In accord with this observation, most of the deletion mutations we observe can already be seen to be taking part in the mirroring effect mentioned above.

For the point mutations, a much smaller list of observable types is available, all of which have been observed *de novo* at least once. In addition, 3 out of the 4 point mutations not reported on HbVar are synonymous (18T*→*G, 18T*→*A and 18T*→*C), for which reason one could not have expected them to be reported on HbVar to begin with. Therefore, it is not possible to apply the mirroring analysis above to the point mutations.

Examining instead the frequencies of alleles in the ROIs reported in population genetic data from gnomAD exome sequencing (*63*) provided by Ensembl (release 102) (*64*), where only sparse data is available, we find that only 3 point mutations in total were reported at non-zero frequencies, all in *HBB*. These are the HbS mutation, 16C*→*G and 16C*→*T. Two are of high and one is of middle *de novo* rate in our data. We also find that the Hb-Leiden mutation, notably the most frequent *de novo* mutation also in the *HBD* ROI, is the only variant of non-zero frequency reported on gnomAD in the *HBD* ROI.

We conclude that the correspondence between *de novo* rates and observations of alleles in populations applies to both types of mutation and extends beyond the HbS and Hb-Leiden mutations. Further adding to this analysis, while HbS is common mostly in Africa and in some populations in the Asian malaria belt (*20*), it appears *de novo* in our African but not in our European donors, and while the Hb-Leiden mutation has been reported across the globe (*25,26*), it appears *de novo* in both our African and European donors.

The correspondence above-mentioned could not have been predicted from the genome-wide average (GWA) rates of the mutations involved. In particular, different indel GWA rates are not assigned to different indel types as defined here but to different indel sizes (the smaller the more frequent; 59), a minor effect which only further contrasts with the fact that the Hb-Leiden deletion (our largest) has the highest *de novo* rate. In addition, the HbS mutation is an A*→*T transversion, the least frequent point mutation type on average.

### 12. Materials and methods

All of the oligos for the sperm DNA library preparation were ordered from IDT with standard desalting purity, unless otherwise mentioned. All enzymes were obtained from New England Biolabs (NEB). PCR purification and agarose gel extraction were carried out with QIAGEN kits.

#### 12.1. Preventing cross-contamination between samples

Because of the ultra-high accuracy of the method, preventing cross-contamination between samples is of the utmost importance. DNA was extracted from semen and prepared for next generation sequencing one donor at a time. Ultra-pure water (DNase and RNase-free) was used for all applications, from semen DNA extraction to library preparation. When possible, reagents were purchased in ready-to-use solutions and used in aliquots that were disposed after each library-preparation cycle together with other dry disposable materials. Non-disposable instruments were cleaned and when possible sterilized according to the manufacturer’s instruction. Working surfaces were cleaned by labZAPTM (A&A Biotechnology) to remove traces of nucleic acid, and the molecular biology work was confined to UV-sterilized cabinets. A spatial separation between pre- and post-PCR procedures was maintained by performing the pre-PCR steps in one lab and the post-PCR steps in a second, remote lab. A unidirectional flow of materials was maintained so that samples, reagents or other items were allowed to move from the pre-PCR lab to the post-PCR lab but never in the opposite direction. Plasmids were constructed in a separate room, mixed and diluted to a working concentration and only then transferred to the pre-PCR room. Other than this final plasmid mix, no reagents or other materials were allowed to move from the plasmid room to the pre-PCR lab area. Additionally, each library was generated with two unique identifier sequences that were added during the primary and the secondary barcode labeling steps, so that in the event that any amplicon sequence from a previous analysis unrelated to the present analysis has infiltrated the sample during one of the library preparation steps, it could be eliminated during the sequence analysis step.

#### 12.2. Spike-in plasmids preparation

Four puc19-based plasmids were generated. Two (ALP13 and ALP17) were designed to carry the *HBB* genomic segment from position -203 to +223 relative to the mRNA translation start site, with the Bsu36I restriction site CCTGAGG replaced with TTATGTT and ACGAGAC, respectively; and two others (ALP16 and ALP18) were designed to carry the *HBD* genomic segment from position -59 to +220 relative to the mRNA translation start site, with the Bsu36I-restriction site replaced with TTATGTT and ACGAGAC, respectively. To prepare the spike-in mixture, the four plasmids were linearized by BamHI, mixed in equal amounts and diluted to 10 femtograms/µl for the AFR1, AFR3, AFR5, AFR6, AFR7, EUR3 and EUR4 samples and 5 femtograms/µl for all other samples.

#### 12.3. Collection of sperm samples

Semen samples from Africans were collected in the Assisted Conception Unit of the Lister Hospital & Fertility Centre in Accra, Ghana following clinical standards, and semen samples from Europeans were purchased from Fairfax, a large US cryobank, with the approvals of the Institutional Review Board of the Noguchi Memorial Institute for Medical Research (NMIMR-IRB 081/16-17) at the University of Ghana, Legon, and of the Rambam Health Care Center Helsinki Committee (0312-16-RMB) and the Israel Ministry of Health (20188768). Donors with a history of cancer or infertility or with high fever in the last 3 months prior to donation were excluded. Informed consent was obtained from all participants and personal identifying information was removed and replaced with codes at the source. The samples were shipped in dry ice or liquid nitrogen according to availability to the University of Haifa for analysis.

#### 12.4. DNA extraction from sperm cells

This DNA isolation protocol is a modified version of the method described by Weyrich (*65*). A semen sample from a single donor was divided into 500µl aliquots in multiple screw-capped tubes. The sperm aliquots were washed twice with 70% ethanol to remove seminal plasma. The remaining cells were rotated overnight at 50*^◦^*C in a 700µl lysis buffer (50 mM Tris-HCl pH 8.0, 100 mM NaCl2, 50 mM EDTA, 1% SDS) containing 0.5% Triton X-100 (Fisher BioReagents BP151-100), 50 mM Tris(2-carboxyethyl)phosphine hydrochloride (TCEP; Sigma 646547) and 1.75 mg/mL Proteinase K (Fisher BioReagents BP1700-100). Lysates were centrifuged at 21,000 x g for 10 minutes at room temperature and supernatants were united in a single tube. DNA purification from the cleared lysate was carried out using QIAGEN Blood & Cell Culture DNA Maxi Kit (13362). Specifically, 5 mL lysate were supplemented by 15 mL of buffer G2 (800 mM guanidine hydrochloride, 30 mM Tris-HCl pH 8.0, 30 mM EDTA pH 8.0, 5% Tween 20, 0.5% Triton X-100), vortexed thoroughly and allowed to gravity-flow through a single Genomic-tip 500/G column pre-equilibrated by 10 mL of buffer QBT (750mM NaCl, 50 mM MOPS pH 7.0, 15% isopropanol (v/v)). Resin was washed twice by 15 mL of Buffer QC (1 M NaCl, 50 mM MOPS pH 7.0, 15% isopropanol (v/v)) and elution was carried out by 15 mL of Buffer QF pre-warmed to 50*^◦^*C (1.25 M NaCl, 50 mM Tris-HCl pH 7.0, 15% isopropanol (v/v)). DNA was precipitated by adding 10.5 ml room temperature isopropanol to the elute, inverting the tube 10 times, and using a sterile tip to spool and transfer the DNA to a screw-capped tube containing 500µl of buffer EB (10 mM Tris-HCl pH 8.5). The DNA was allowed to dissolve overnight at room-temperature. For each donor, a small aliquot from the extracted DNA was PCR amplified and Sanger sequenced to verify the exact sequence of *HBB* and *HBD* regions.

#### 12.5. Enzymatic digestion

For the RE-1–treated sample, roughly 264 µg sperm DNA, equivalent to 80 million haploid cells (For AFR2 DNA amount equivalent to 60 million cells was used), were mixed with a plasmid spike-in mixture (0.2 pg for AFR1 and 0.1 pg for other donors) and equally divided in a 96-well plate. Bsu36I-HF digestion (RE-1) was carried out overnight at 37*^◦^*C according to the manufacturer’s instructions using 5 units per well. Then, each well was supplemented by 6 units of HpyCH4III and digestion continued for three more hours. For the RE-1–untreated reaction (no Bsu36I digest), 13.2 µg sperm DNA (and 9.9 µg for AFR2), representing 5% of the DNA amount used for Bsu36I digest, were mixed with 6 times the volume of plasmid spike-in mixture, aliquoted to 5 tubes and incubated overnight with 2 units SalI-HF per tube instead of Bsu36I to allow for similar conditions of DNA digestion without affecting the Bsu36I and HpyCH4III sites. Then, each well was supplemented by 6 units of HpyCH4III (RE-2) and digestion continued for three more hours. All digestion products were purified using a QIAGEN PCR purification kit.

#### 12.6. Primary-barcode labeling and linear amplification

Direct barcode labeling and linear amplification of the digested *HBB* and *HBD* strands were carried out in a single reaction in 96-well plates. Each well contained about 1µg of digested DNA, 0.1 µM primary-barcode oligo (oligo A) and 1 µM of 5’-phosphorothioate-protected primer for linear amplification (oligo B). The reaction was carried out with Q5 high-fidelity polymerase according to the manufacturer’s instructions, using the following thermocycler parameters: initial denaturation at 98*^◦^*C for 20 seconds, followed by 16 cycles of 98*^◦^*C for 5 seconds, 68*^◦^*C for 15 seconds, and 72*^◦^*C for 20 seconds. The DNA was purified using a PCR purification kit. For each donor, each of the Bsu36I-treated and untreated samples was labeled by an oligo A with a different Donor identifier-1 (ID-1) sequence, which was also not shared by samples from other donors, making it so that each donor and each condition had its own identifier sequence.

#### 12.7. 5’-exonuclease treatment

To eliminate non 5’-phosphorothioate-protected strands, 15µg DNA aliquots from the post linearly amplified product of the Bsu36I-treated sample were incubated each at 37*^◦^*C in the presence of 15 units of Lambda exonuclease, 30 units of T7 exonuclease and 90 units of RecJF exonuclease in 1x CutSmart buffer for 2.5 hours. The post linearly amplified product of the Bsu36I-untreated sample was incubated at the same conditions with 10 units of Lambda exonuclease, 20 units of T7 exonuclease and 60 units of RecJF exonuclease. The DNA was purified using a QIAGEN PCR purification kit.

#### 12.8. Secondary-barcode labeling and 3’-exonuclease treatment

The DNA was aliquoted into a 96-well plate (1 µg per well). A single primer extension reaction was carried out using 0.5 µM of the secondary-barcode primer (oligo C) and Q5 high-fidelity polymerase according to manufacturer’s instructions. The following thermocycler parameters were used: initial denaturation at 98*^◦^*C for 20 seconds, followed by a single cycle of 98*^◦^*C for 5 seconds, 68*^◦^*C for 15 seconds, and 72*^◦^*C for 40 seconds. To remove excess oligo C, immediately after the thermocycler temperature dropped to 16*^◦^*C, 20 units of thermolabile Exo I were added directly to each well together with the relabeling control primer (oligo D) in a known amount equivalent to 0.66% of the secondary-barcode primer. After incubation of one hour at 37*^◦^*C, the thermolabile Exo I was heat-inactivated by one minute at 80*^◦^*C and the DNA was purified using a PCR purification kit. For each donor, each of the Bsu36I-treated and untreated samples was labeled by an oligo C with a different Donor identifier-2 sequence (ID-2), which was also not shared by samples from other donors, making it so that each donor and each condition had its own identifier-2 sequence.

#### 12.9. PCR amplification and sequencing

The first PCR reaction of the dual-barcode labeled product was carried out using oligo E and oligo F1 as primers and Q5 high-fidelity polymerase, according to manufacturer’s instructions. The following thermocycler parameters were used: initial denaturation at 98*^◦^*C for 30 seconds, followed by 10 cycles of 98*^◦^*C for 5 seconds, 72*^◦^*C for 15 seconds, 72*^◦^*C for 30 seconds, and a final extension at 72*^◦^*C for 30 seconds. Amplification products were purified using a PCR purification kit. The second PCR reaction was carried out using 25% of the first PCR product as template, the amplification primers E and F2, and Q5 high-fidelity polymerase according to the manufacturer’s instructions (different F2 primers were used in order to add a unique Illumina index sequence to each Bsu36I-treated and untreated sample). The following thermocycler parameters were used: initial denaturation at 98*^◦^*C for 30 seconds, followed by 24 cycles (with the exception of EUR4 sample that was amplified by 17 cycles) of 98*^◦^*C for 5 seconds, 70*^◦^*C for 15 seconds, 72*^◦^*C for 30 seconds, and a final extension at 72*^◦^*C for 1 minute. PCR products were agarose-gel purified using QIAGEN gel extraction kit, and further concentrated by a DNA clean & concentrator kit (Zymo Research). DNA libraries prepared from the Bsu36I-treated and untreated samples of the same donor were mixed in equal amounts and paired-end sequenced with 20% PhiX by Illumina MiSeq 300 cycles kit (V2) at the Technion Genome Center (TGC). For each donor two or three MiSeq runs were performed to reach a minimum of 10 million reads per treatment (specifically, all but AFR5 and EUR3 were sequenced two times) and the resulting fastq sequences were joined prior to the sequence analysis step.

#### 12.10. Sequence analysis

Illumina paired-end (PE) reads were merged via Pear using the default model for the detection of significantly aligned regions and Phred score corrections. Merged sequences were trimmed from Illumina adapters using Cutadapt, and quality-filtered by Trimmomatic using a sliding window size of 3 and a Phred quality threshold of 30. Quality filtered sequences were trimmed to remove the 5’ edge up to position 18, a sequence which includes the 14 bases of the primary barcode and the 4 bases of ID-1, while adding this information to the read’s header. Only sequences with the correct ID-1 and first three bases of *HBB* or *HBD* sequences were maintained. Similarly, sequences were trimmed from 9 bp at their 3’ edge, which include the 5 bases of the secondary barcode and the 4 bases of ID-2, while adding this information to the read’s header. Only sequences with the correct ID-2 were maintained. Trimmed sequences were sorted to *HBB* or *HBD* sequence pools, based on the occupying bases at positions 33-38 of the coding sequence (CGTTAC for *HBB* and TGTCAA for *HBD*), allowing one mismatch and frameshifts of up to -3 or +3. Successfully sorted sequences were mapped to either the *HBB* or *HBD* reference sequences (obtained by sanger sequencing aliquots from the matching donor samples) using BWA (parameters -M -t), and high-quality mutations (Phred score *≥* 28) were noted. Reads were grouped by their primary barcodes to ‘families’ and processed according to the workflow depicted in Figure S9.

#### 12.11. Bsu36I single-base substitution sensitivity assay

Nineteen oligos (BSU 1-19) carrying the first 37 bases of *HBB*, each with a randomized nucleotide at a single position within the seven bases of the Bsu36I recognition site or at one of the six bases that flank this site from either side were mixed with a similar oligo with all of the seven bases of the Bsu36I recognition site replaced by the sequence TTATGTT (Bsu36IR). This oligos mixture was PCR-amplified for 25 cycles using Q5 DNA polymerase and primers that match the Illumina adapter sequences flanking the *HBB* region in each oligo (Primers BSU F1 and BSU R1). 150 ng of this PCR product were subjected to 20 hours incubation at 37*^◦^*C with or without 5 units Bsu36I. Digestion products were purified, re-amplified (Primers BSU F2 and BSU R2) and paired-end sequenced by Illumina MiSeq. To calculate Bsu36I sensitivity, mutation variant reads were counted and the frequency of each variant in the Bsu36I-treated sample was divided by its frequency in the Bsu36I-untreated sample. All ratios were normalized to the ratio of the Bsu36IR variant that was considered to be 100% resistant to Bsu36I.

### 13. DNA oligos

#### 13.1. Oligos for Sperm DNA library preparation

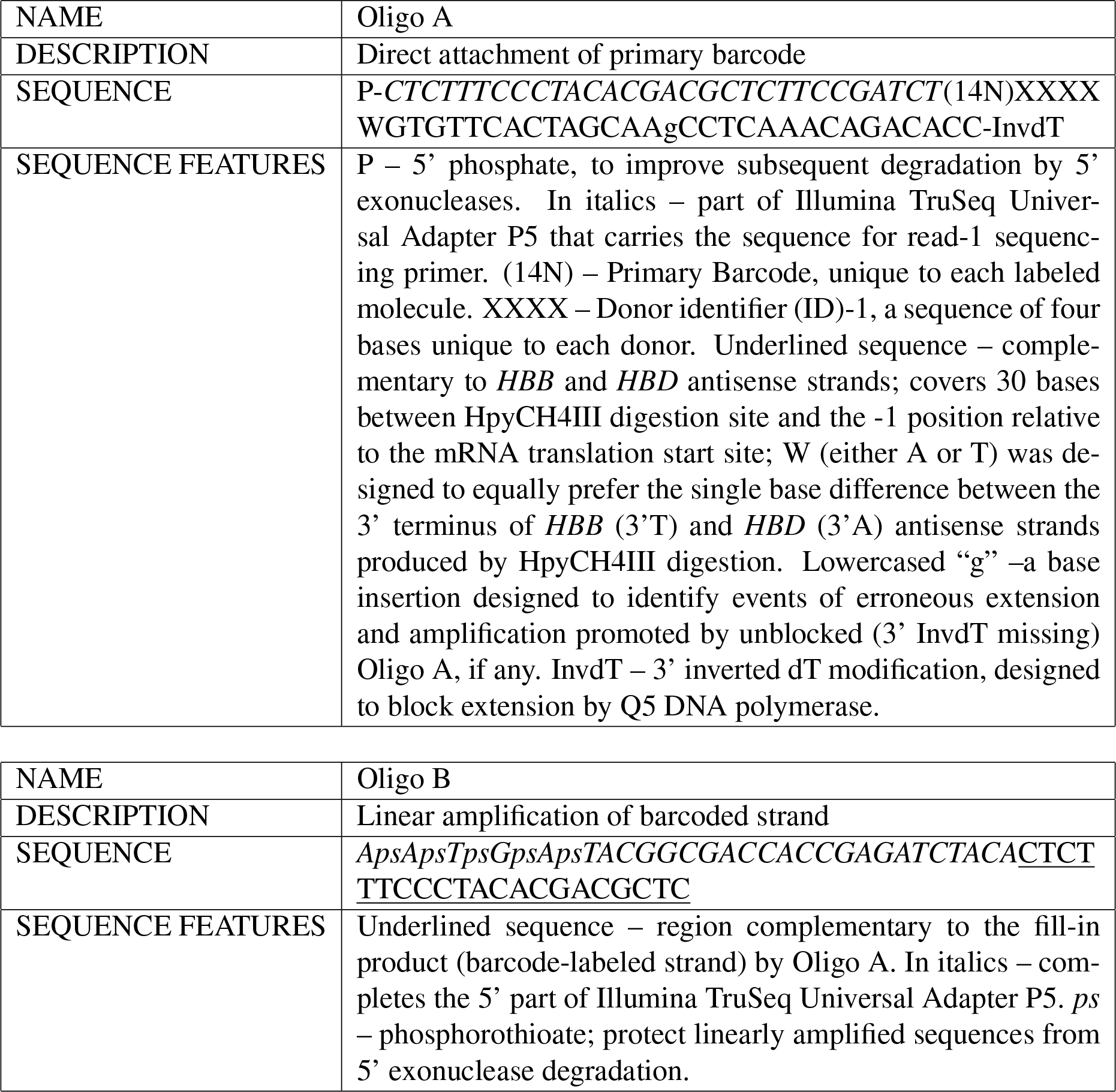

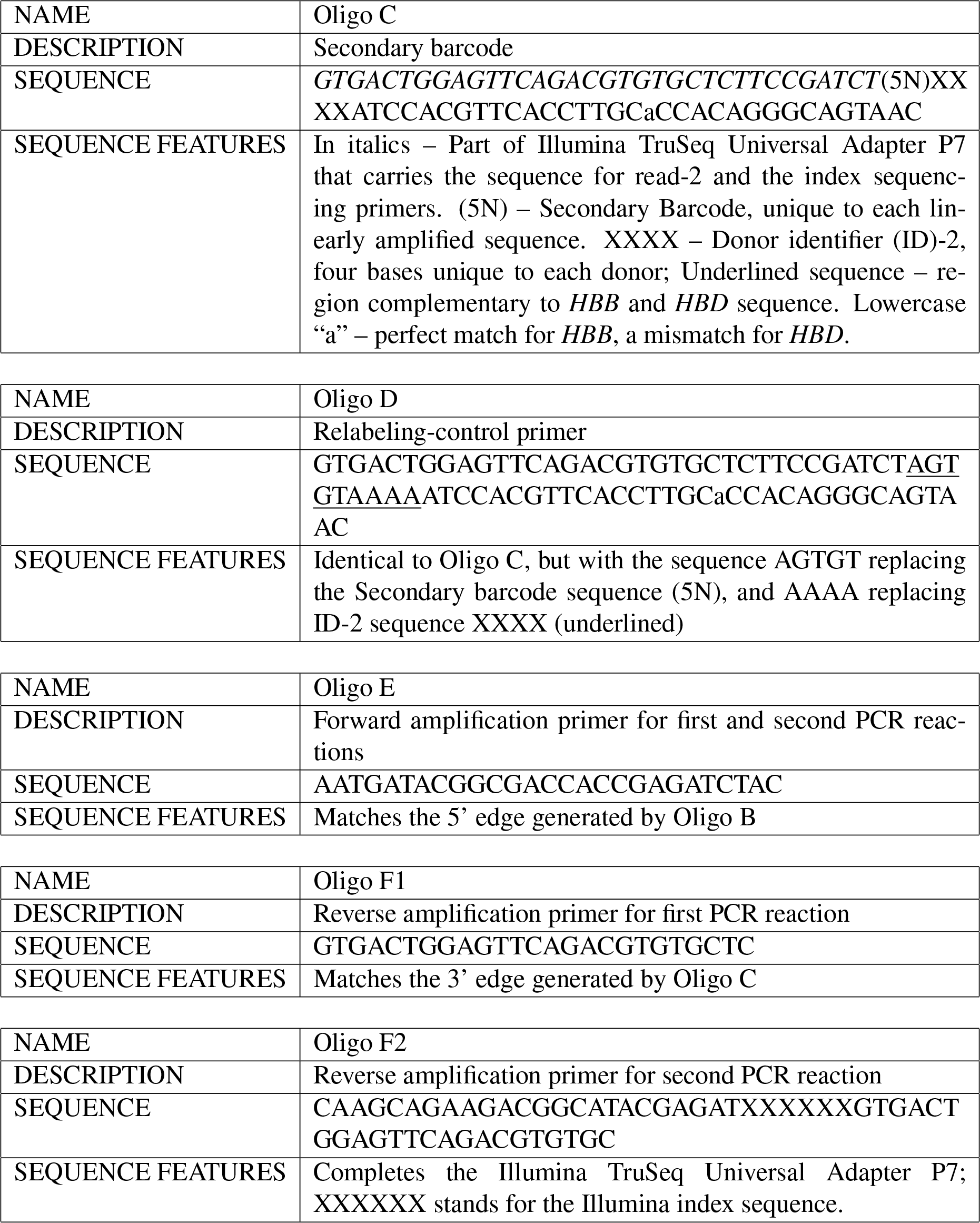

#### 13.2. Oligos for Bsu36I-mutation sensitivity test

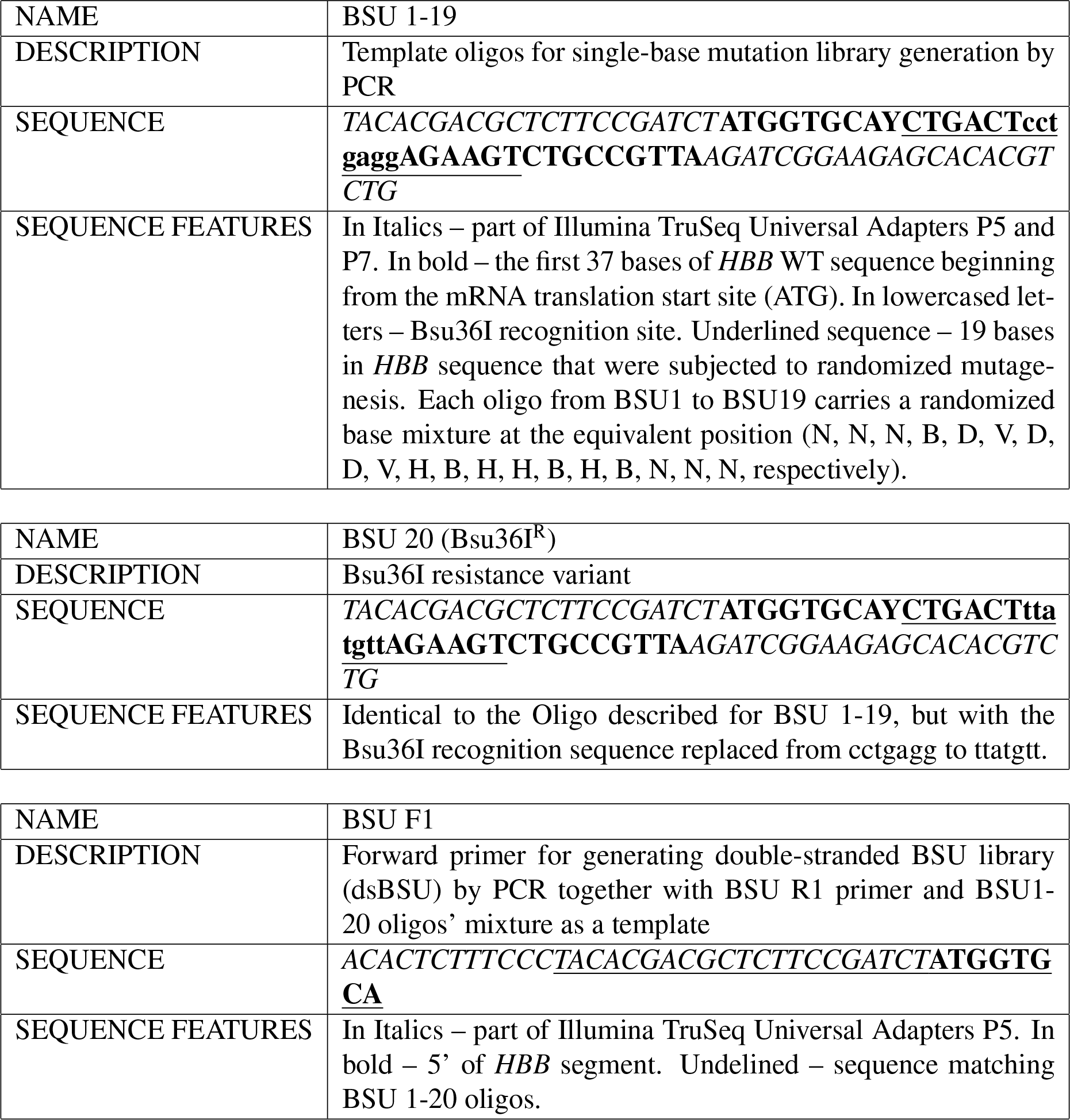

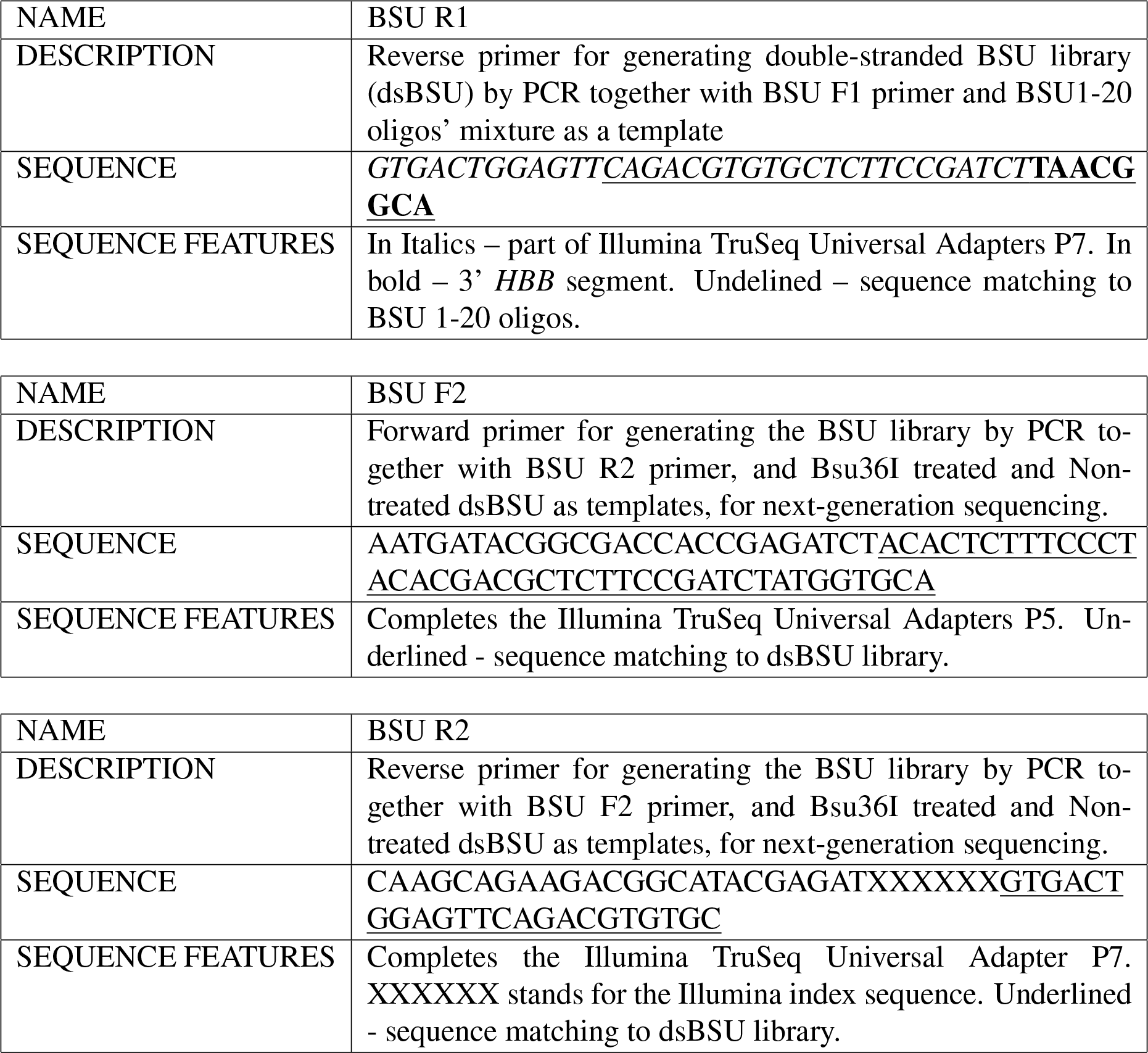

### 14. Supplementary datasheets

The below supplementary datasheet files accompany this manuscript:

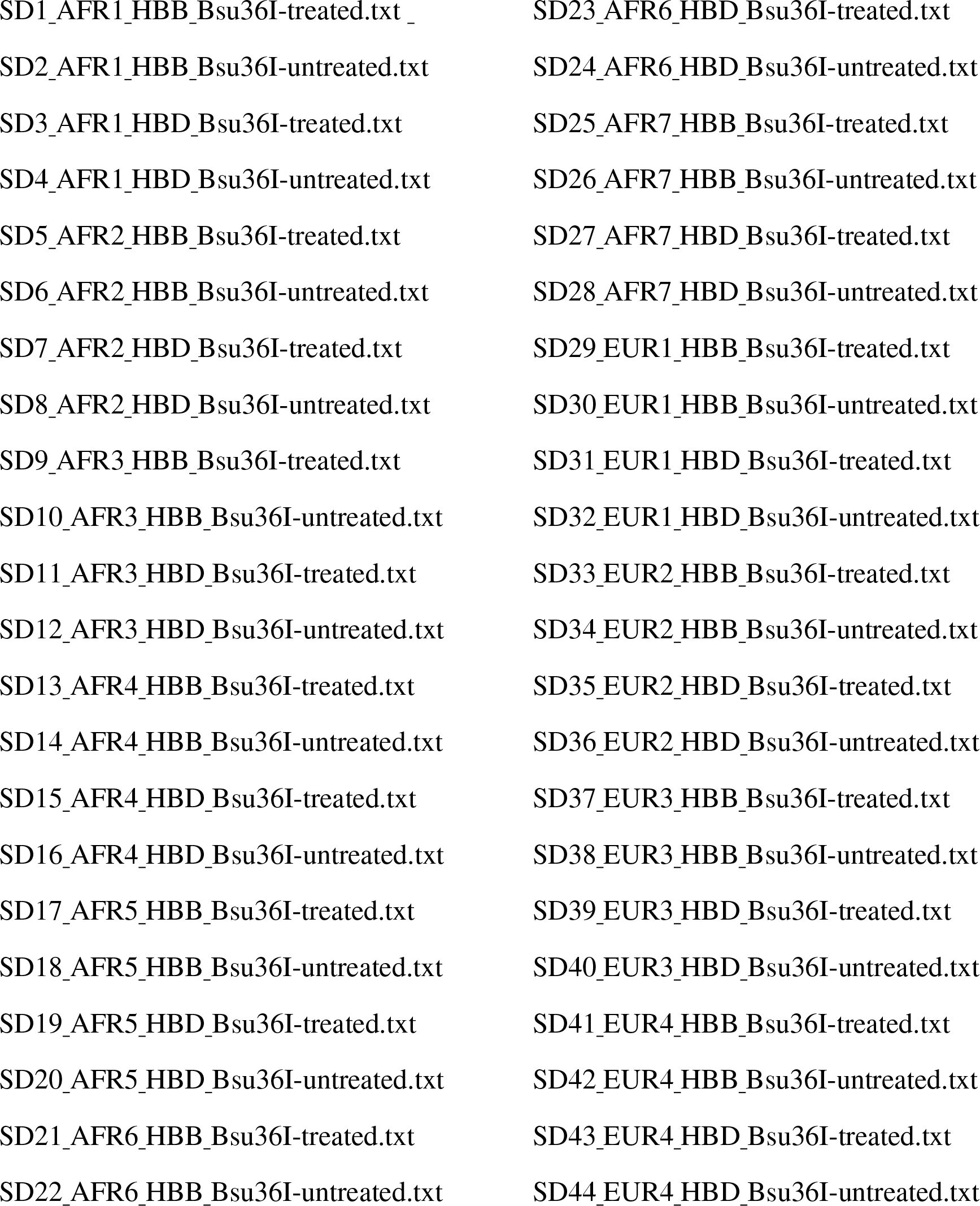

### Description

These files describe features of primary-barcode families that were approved for analysis according to the following cutoff criteria:

A. Family size *≥* 4
B. Mutation frequency *≥* 0.7
C. Secondary barcode *≥* 2

For each primary-barcode family, the data below are shown:

1. Primary barcode: 14 bases of a randomized-barcode sequence that tag a single gene (*HBB* or *HBD*) from a single sperm cell.
2. Read consensus: The consensus sequence obtained after using the combined cutoff criteria shown above. “WT” stands for a wild-type sequence in the ROI and the ROI flanking sequences (i.e., no mutation passed the mutation-frequency and secondarybarcode cutoff criteria in these primary-barcode families). When a mutation is specified, its identity is described by its position relative to the first sequenced base in the read after the sample identifier-1 (ID-1) sequence, followed by the identity of the original base at that position and the identity of the substituting base (for example, 50AT stands for an A*→*T substitution at position 50). Mutation identifiers with a hyphen located at the position of the original base or the substituting base describe insertion or deletion mutations respectively. When multiple mutations in a single primary-barcode family are approved by the combined cutoff criteria, the mutation identifiers are separated by semicolons.
3. HBB consensus (or HBD consensus): The same as for the Read consensus, but with the mutation position in the mutation identifier adjusted to the mRNA-translation start site.
4. Mutation frequency: The fraction of reads that carry the consensus mutation in the primary-barcode family. Due to the applied mutation-frequency cutoff, only mutations (or WT bases) with a frequency of at least 0.7 are shown. When multiple mutations in a single primary-barcode family are approved by the combined cutoff criteria, the mutation frequencies are separated by semicolons. For the WT consensus sequences, a mutation-frequency value represents the fraction of reads in the primarybarcode family with no mutations. PCR or sequencing errors may reduce the fraction of complete WT sequences in a primary barcode family, sometimes below the mutation frequency cutoff of 0.7. Yet, since the mutation-frequency cutoff is applied for a single position at a time, the frequency of the WT base in each of these mutated positions exceeded the mutation-frequency cutoff.
5. Mutation count: The number of reads that carry the consensus mutation in a primarybarcode family (the numerator for the mutation frequency calculation). When multiple mutations in a single primary-barcode family are approved by the combined cutoff criteria, the mutation counts are separated by semicolons. For the WT consensus sequences, a mutation-count value represents the number of reads in the primarybarcode family with no mutations.
6. Total count: The number of reads in a primary-barcode family (the denominator for the mutation frequency calculation). Due to the applied family-size cutoff, only families with more than 4 reads are shown.
7. Number of mutations: The number of mutations in a primary-barcode family that were approved by the combined cutoff criteria.
8. Unique secondary barcodes: The number of unique-secondary barcodes that are associated with the consensus mutation in a primary-barcode family. Due to the applied secondary-barcode number cutoff, only mutations (or WT bases) with 2 or more secondary barcodes are shown. When multiple mutations in a single primarybarcode family are approved by the combined cutoff criteria, the mutation frequencies are separated by semicolons. For the WT consensus sequences, if a mutation in the primary-barcode family was disqualified by the combined cutoff criteria (i.e., PCR or sequencing errors), the number of unique-secondary barcodes associated with the WT base at that position is shown (with multiple positions with disqualified mutations producing multiple unique-secondary barcodes values for the WT consensus sequence, separated by semicolons.
9. The HBB consensus sequences 9CT;16C-;17C-;19GT;21GT;22-TT and 9CT;16CA;18T-;21-A;22GC belong to the spike-in plasmids ALP13 and ALP17 carrying the artificial ROI sequences TTATGTT and ACGAGAC, respectively, instead of the Bsu36I site CCTGAGG. The HBD consensus sequences 16C-;17C;19GT;21GT;22-TT and 16CA;18T-;21-A;22GC belong to the spike-in plasmids ALP16 and ALP18 that carries the same artificial ROI sequences. The consensus sequence 39CT;45-ATAA;47CA;49G-;50A-;51G-;52G-, appearing twice in the AFR1 HBB Bsu36I-treated, once in EUR2 HBB Bsu36I treated and once in AFR7 HBB Bsu36I treated datasets is an ALP21 plasmid (ATAACAT instead of the Bsu36I site) contaminant that was not used in this study.

---

**Figure S1:**
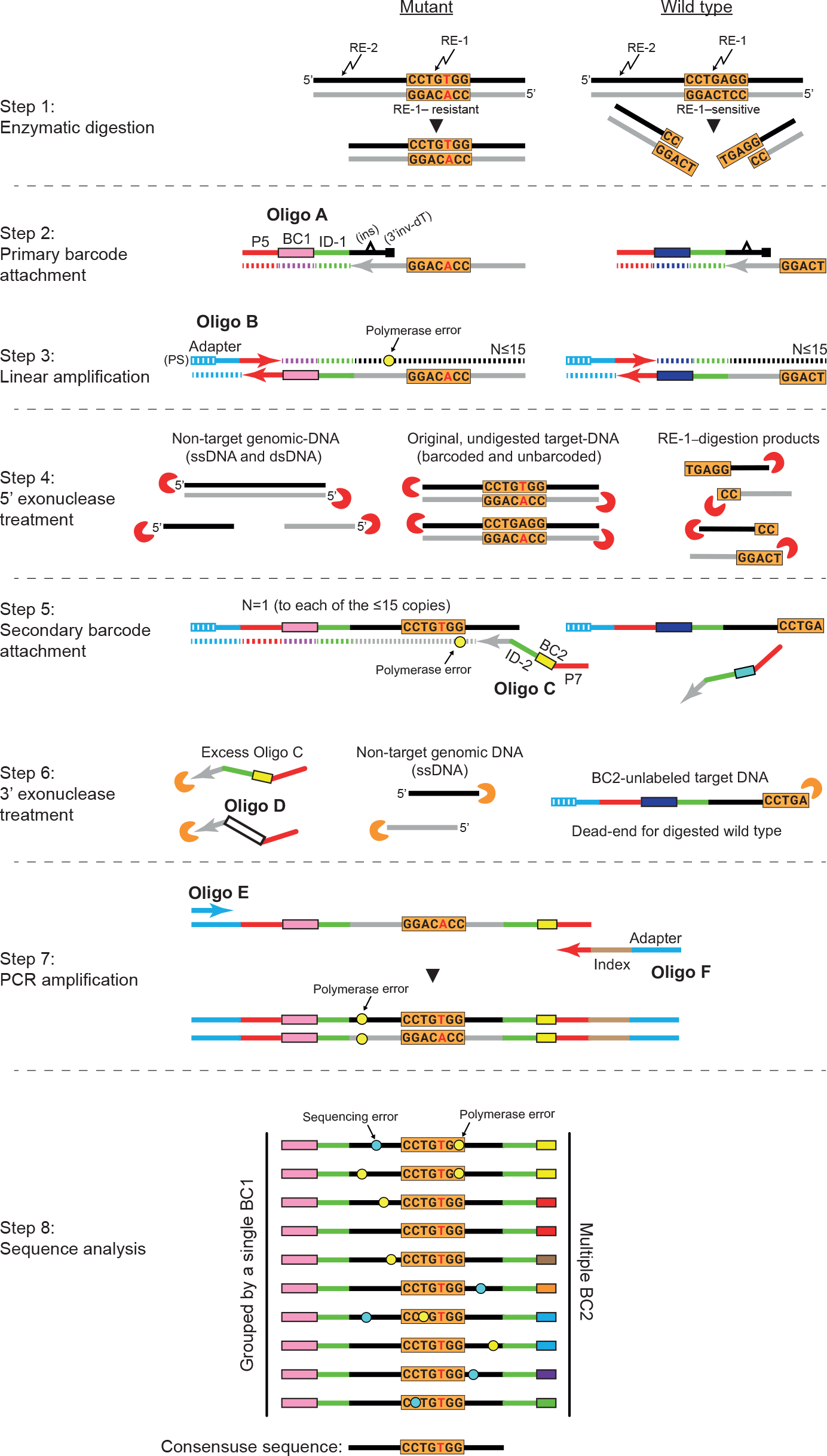
An illustration of the MEMDS method. Step 1) Enzymatic digestion of genomic DNA. Restriction Enzyme-1 (RE-1) digests a region of interest (ROI) with a wild-type sequence and is blocked by a mutation at this site (the mutation is marked by a red letter). Restriction Enzyme-2 (RE-2) digests in close proximity to the ROI. Step 2) A fill-in reaction is promoted by oligo A, which anneals to the sequence between the RE-1 and RE-2 sites and introduces directly to the target DNA strand (gray strand) a sample-identifier sequence (ID-1) common to all labeled sequences in the sample, a primary-barcode sequence (BC1) that is unique to each target DNA molecule and an Illumina-P5 sequence. Shown also are the 3’ inverted-dT (3’inv-dT) that blocks oligo A extension and the control base-insertion (ins) for the identification and removal of extension products from unblocked oligo A at the sequence analysis step. Step 3) Linear amplification of the barcoded target molecules is carried out for 15 cycles using oligo B that anneals to the P5 sequence of the target DNA strand and adds an Illumina adapter sequence with 5 phosphorothioate bonds at the 5’ edge (5’PS) of each barcoded-ROI copy. This linear amplification reaction results in 15 or less copies of each barcoded target molecule (N ≤ 15). While polymerase errors (marked by yellow circles) do occur, they are unlikely to repeat themselves at the same position in multiple copies of the same target molecule. Step 4) A mixture of 5’ exonucleases (red pacman) is added to degrade from 5’ to 3’ non-target genomic DNA including RE-1 and RE-2 digestion products. The barcoded copies are protected from this degradation step due to their 5’PS bonds. Step 5) A single-extension reaction with oligo C is carried out in order to add to each linearly amplified copy of a BC1-labeled target molecule an additional sample identifier sequence (ID-2), a unique secondary-barcode sequence (BC2), and an Illumina-P7 sequence. For each target molecule, this step results in 15 or less copies that share the same primary barcode at one end, while having a unique secondary barcode at the other end. Since oligo C anneals 3’ to the ROI sequence, any linearly amplified copy of RE-1 digestion product cannot be labeled by this oligo. Step 6) A 3’ exonuclease (orange pacman) is added immediately after the single-extension reaction to degrade from 3’ to 5’ any single-stranded DNA, including excess of oligo C, to prevent secondary-barcode relabeling during the next PCR reaction. Copies labeled by secondary barcodes are protected from this degradation step due to their double-stranded state, while non-labeled copies are single-stranded and are therefore degraded. A relabeling-control primer (oligo D), carrying a unique sequence signature, is added in known amount together with the 3’ exonuclease to assess, at the sequence analysis step, the number of oligo C relabeling in the event of incomplete degradation of oligo C by the 3’ exonuclease. Step 7) PCR amplification completes the final sequence requirements for Illumina NGS and produces a library of barcoded ROI sequences composed of enriched mutation variants as well as wild-type sequences that escaped RE-1 digestion. Step 8) Following next-generation sequencing, reads are grouped into families based on their primary-barcode sequences, so that within each family, all members have the same primary barcode, and the consensus sequence for the family is determined using three parameters: family size, mutation frequency, and the number of secondary barcodes associated with each base. This procedure allows us to eliminate PCR errors (yellow circles) and NGS errors (blue circles), which usually appear in low frequencies and are linked to single secondary barcodes, and to accept as true, *de novo* mutations only mutations that appear in multiples reads and are associated with multiple secondary barcodes, such as the “T” substitution in the figure.

**Figure S2:**
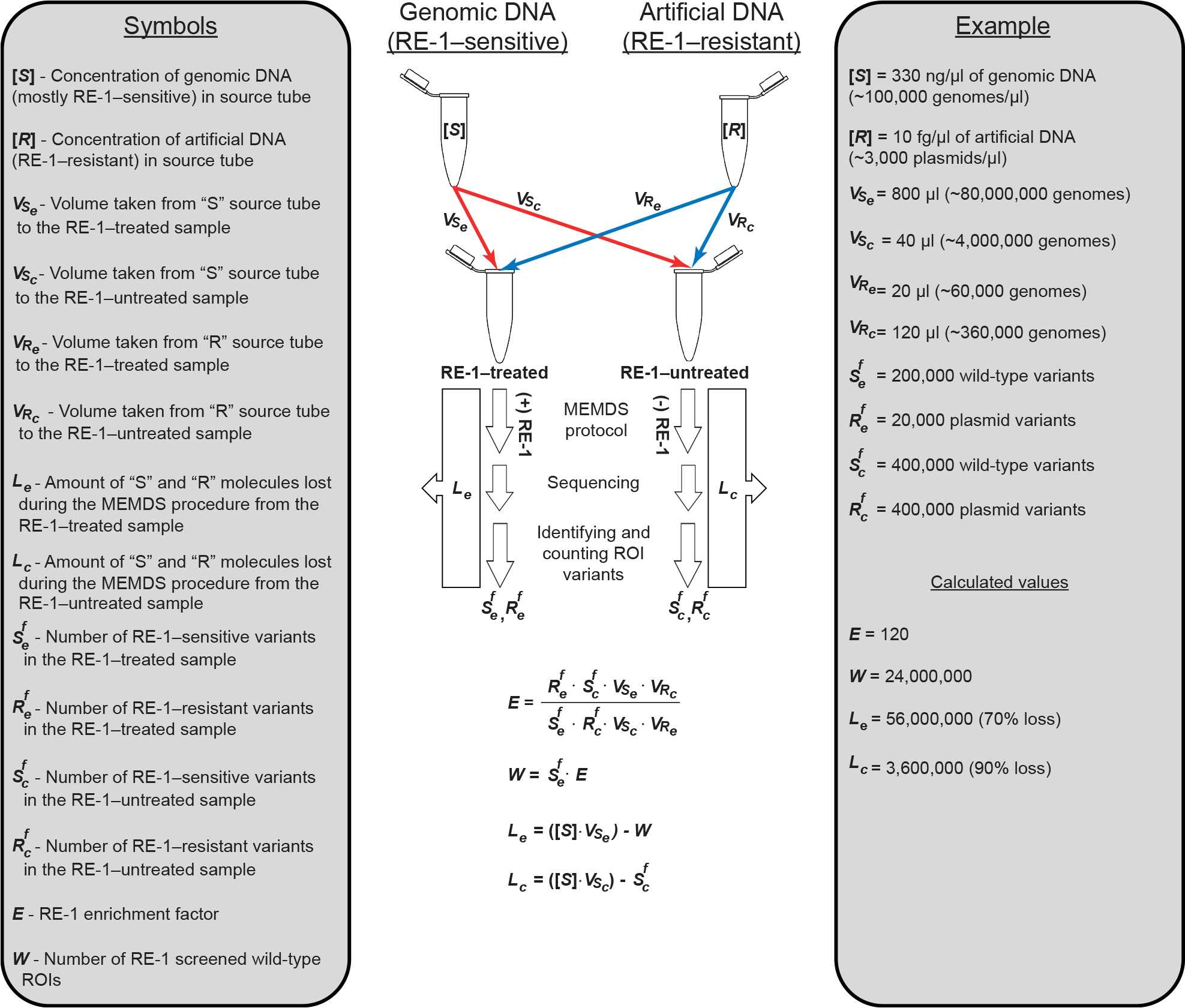
MEMDS Experimental design to calculate the RE-1–enrichment factor and the number of target DNA molecules digested by RE-1. Two tubes, one containing genomic DNA that carries mostly RE-1–sensitive ROI sequences, denoted *S*, and one containing artificial-ROI sequences resistant to RE-1 digestion, denoted *R*, are used as source tubes from which volumes are drawn in known amounts in order to create two mixtures of the two samples, designated “RE-1–treated” and “RE-1–untreated” samples (see left panel for the abbreviations used). These two samples undergo the full MEMDS protocol, with the exception that the former is treated with and the latter without RE-1. Following next-generation sequencing, variants are identified by the MEMDS computational pipeline and the numbers of RE-1–sensitive ROI variants (i.e., wild-type ROIs) and artificial RE-1–resistant ROI variants are determined for each sample (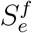 and 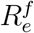 denote the numbers of sensitive and resistant variants identified for the RE-1 treated sample, and 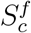 and 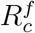 denote the sensitive and resistant variants identified for the RE-1 untreated sample). These numbers, together with the known volumes taken from the source tubes to create the input mixtures, are used to calculate the RE-1–enrichment factor, *E*, and the total number, *W*, of wild-type sequences that were either digested by RE-1 or evaded digestion and were sequenced and identified. The right panel shows an example for the calculation of *E* and *W* based on DNA concentrations in each source tube and volumes taken for the input mixtures that are similar to the DNA concentrations and volumes used in our MEMDS experiments but are rounded for the sake of simplicity here. In this example, the RE-1–enrichment factor equals 120, meaning that *de novo* mutations in the ROI, which block RE-1 similarly to the artificial sequences, are enriched 120-fold in the RE-1–treated sample compared to the RE-1–untreated sample. Using this enrichment factor we find that the total number of unique wild-type molecules screened by the MEMDS procedure is 24,000,000, which includes the number of wild-type target molecules in the RE-1– treated sample that were digested by RE-1 and the 400,000 RE-1–sensitive variants that escaped digestion and were sequenced. Note that the calculation of *E* and *W* relies only on the number of original target molecules that were sequenced in the computational analysis step and on the volumes used to generate the input mixtures, and therefore the number of genomes and artificial sequences in the source tubes is not needed for it. Yet, by having a rough estimate of the actual amount of DNA transferred from the source tube, one can assess the number of target DNA molecules (either genomic or artificial ROI-including molecules) that were lost during the MEMDS procedure (not due to RE-1 digestion but due to general loss of material involving the efficiencies of labeling, amplifying, purifying, capturing and sequencing all target sequences).

**Figure S3:**
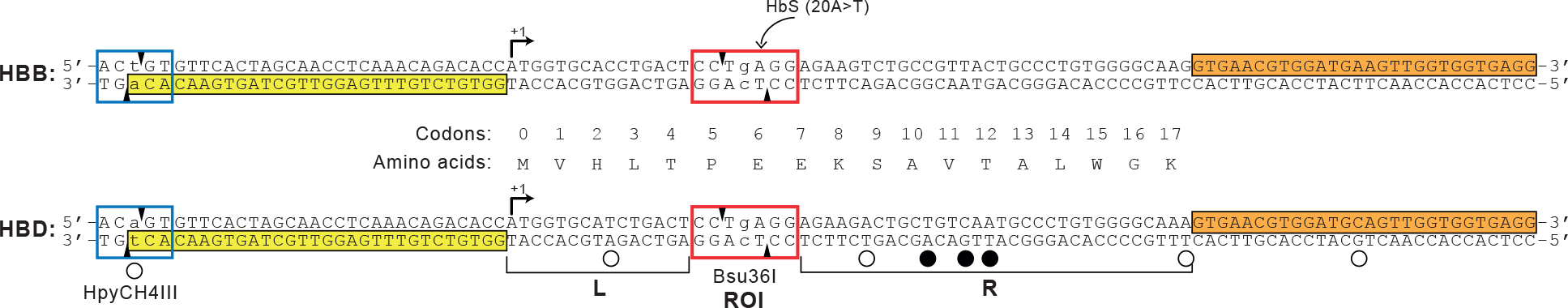
*HBB* and *HBD* sequence features. The double-stranded 114 bp DNA segments from the first exon of *HBB* (upper sequences) and the homologous region of *HBD* (lower sequences) are shown. The mRNA-translation start sites (ATG) are marked by black arrows. For each gene, the upper sequence is in the sense orientation and the lower, antisense, complementary sequence served as the target DNA strand, which was barcoded and subsequently amplified by the MEMDS protocol. Positions that vary between the two genes are marked by circles below the *HBD* segment. Positions marked by filled circles were used to sort NGS reads from the same sperm-DNA sample to separate *HBB* and *HBD* datasets at the sequence analysis stage, as the two genes were barcoded and amplified simultaneously by the MEMDS procedure. The Bsu36I (RE-1)-recognition sequence is marked by a red frame and its cleavage sites are marked by small black triangles. Position 20, where the HbS (20A T) mutation occurs, is marked by a curved arrow. The base denoted by a lower-case letter in the center of the Bsu36I site can tolerate any substitution without affecting Bsu36I activity. Therefore, the region of interest (ROI) is confined to six of the seven bases in the red frame that constitute the Bsu36I site. The HpyCH4III (RE-2)-recognition sequence is marked by a blue frame and its cleavage sites are marked by small black triangles. The base denoted by a lower-case letter in the HpyCH4III site can tolerate any substitution without affecting HpyCH4III activity. The sequence in the yellow box anneals to oligo A and receives the primary barcode via a single, fill-in reaction (see Figure S1, step 2). Note that the first base that primes this extension, marked by a lower-case letter, differs between *HBB* and *HBD*. Therefore, we used a mixture of oligo A sequences that carry either one of the two complementary bases to minimize any bias due to delayed extension by the Q5 DNA polymerase. The sequence in the orange box anneals to oligo C and receives the secondary barcode via a single extension reaction (see Figure S1, step 5). The sequence between oligo A and oligo C remains untouched by any primer and therefore is suitable for mutation detection analysis. Yet, only mutations at the ROI can be enriched, while mutations in the flanking right (R) and left (L) sequences are unlikely to affect Bsu36I digestion.

**Figure S4:**
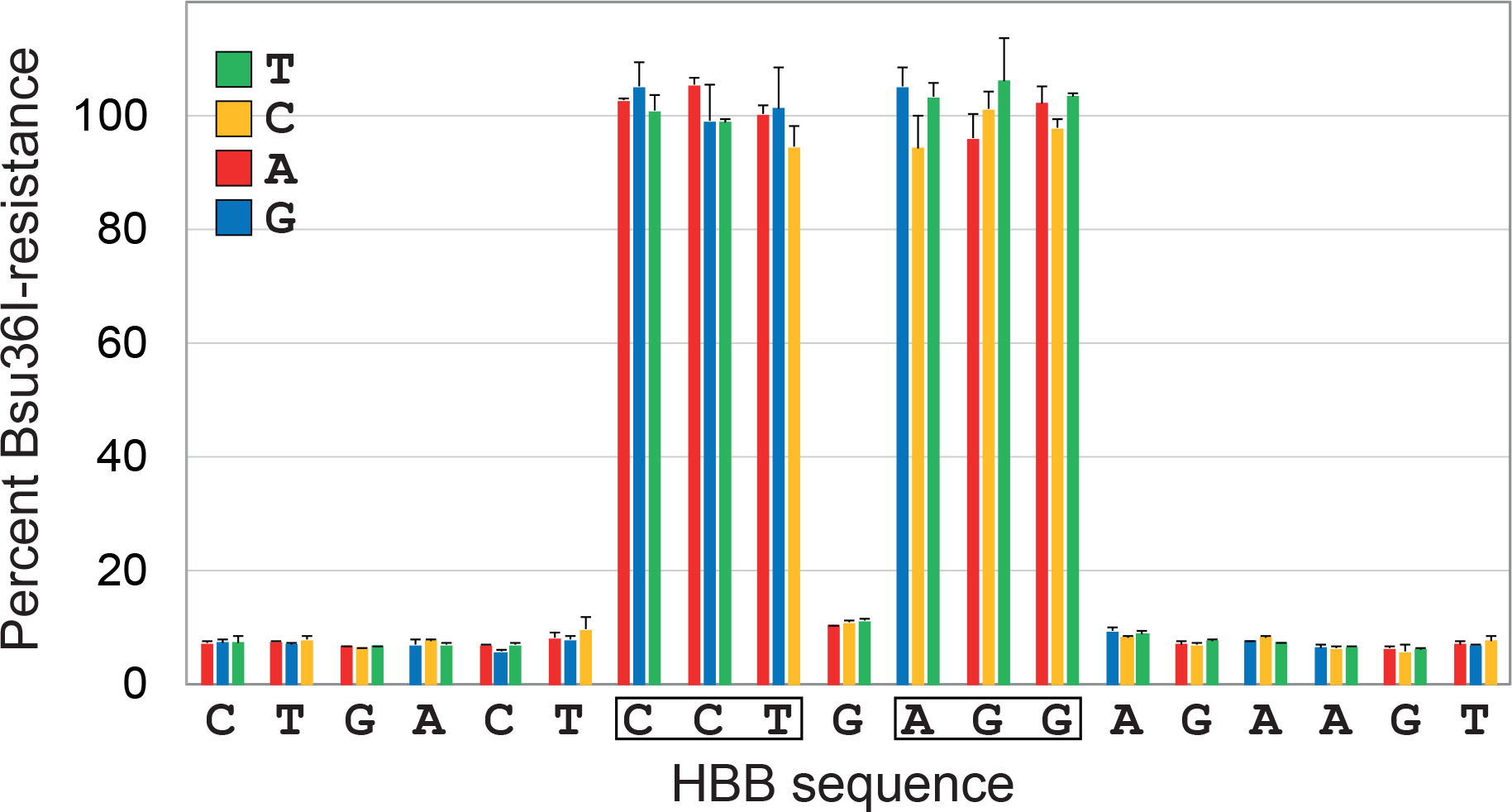
Single-base substitutions protect from Bsu36I digestion similarly to the MEMDS-artificial ROI variant that carries multiple changes in the Bsu36I site. A synthetic dsDNA library of *HBB* gene segments containing the Bsu36I-restriction site with its flanking sequences and a single point mutation per segment was divided into two and sequenced following incubation with or without Bsu36I. As Bsu36I digestion results in the depletion of sequences that are Bsu36I-sensitive, the frequency of each full-length variant in the post-Bsu36I treatment pool compared to its frequency in the pre-Bsu36I treatment pool was used to determine its degree of Bsu36I-resistance. Changes in frequencies were normalized to the change in frequency of an artificial variant with multiple changes in the Bsu36I site, which was set to 100% resistance (n=2). The six bases that constitute the *HBB* ROI are shown in boxes and the identities of the substituting bases are color-coded. Note that Bsu36I-sensitive variants are not completely depleted from the post-Bsu36I treatment pool, probably due to Bsu36I-resistant heteroduplex DNA that carry a Bsu36I-sensitive sequence in one strand and a Bsu36I-resistant sequence in the second strand, formed during the PCR reaction that generated the input dsDNA library.

**Figure S5:**
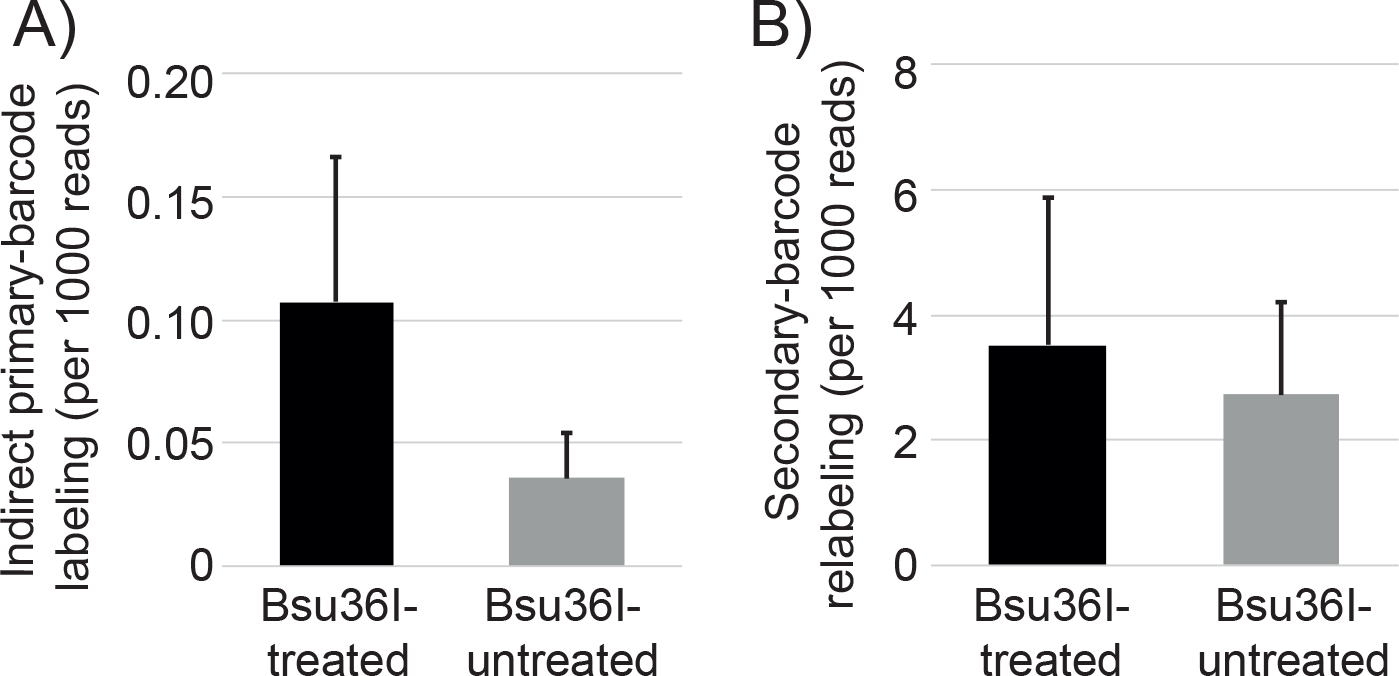
Frequency of erroneous barcode labelings. A) Frequency of indirect labeling by the primary-barcode oligo (oligo A) as measured by the fraction of reads carrying the control-guanine insertion. B) Frequency of secondary-barcode primer (oligo C) relabeling as measured by the relative frequency of reads carrying the sequence signature of the control secondary-barcode relabeling primer (oligo D).

**Figure S6:**
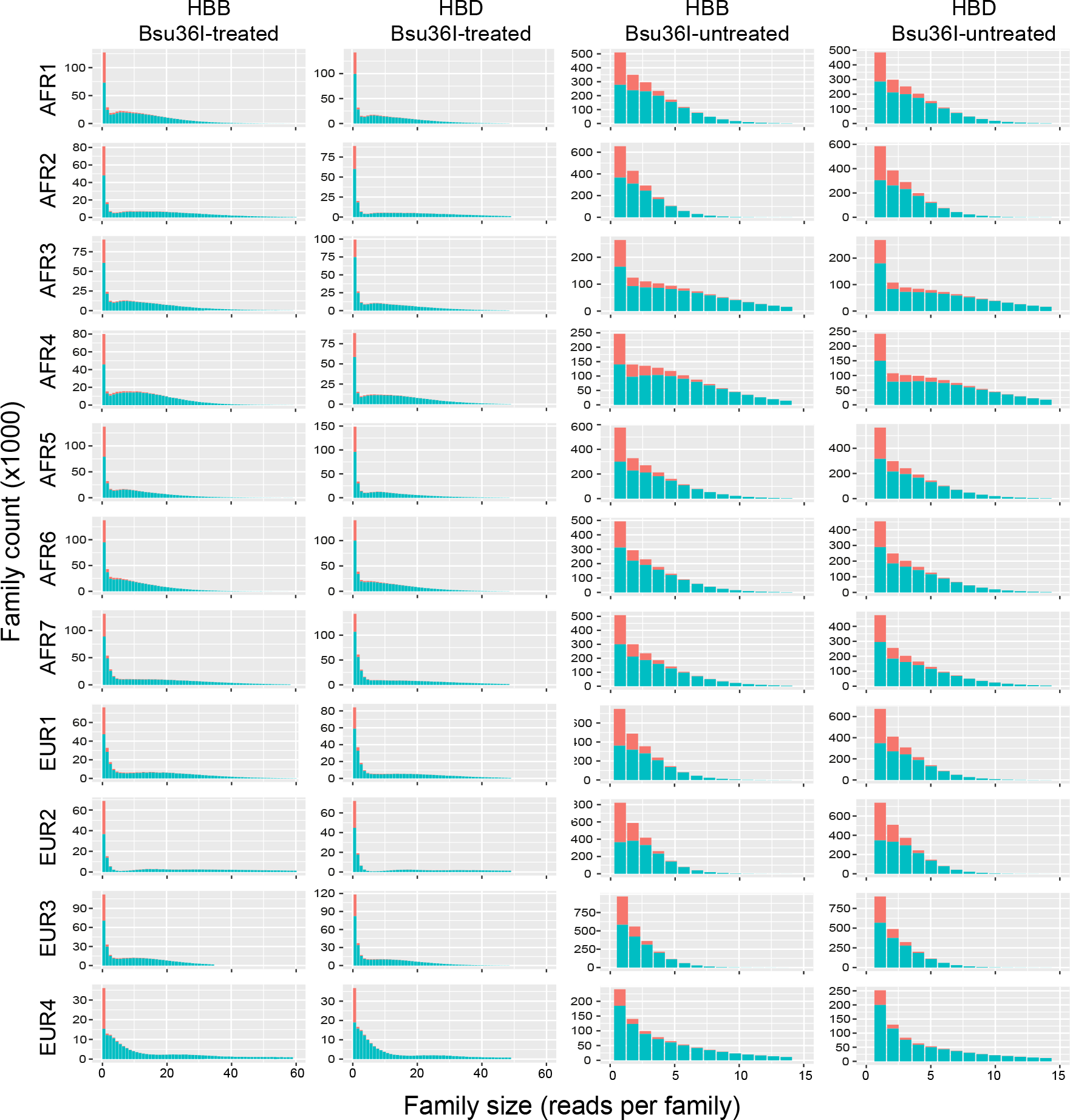
Family-size distributions. Distributions of primary-barcode families based on the number of read in a family (family size). In red: counts of families with primary barcode sequences that deviate by a Hamming distance of one from primary barcode sequences of families with a greater number of reads. In green: counts of families with primary barcode sequences that deviate by a Hamming distance *>* 1 from primary barcode sequences of families with a greater number of reads. Note the different scales used for the Bsu36I-untreated and treated samples. (The differences in family size between the two treatments are merely due to the higher recovery of ROI families in the Bsu36I-untreated samples, which lack depletion of wild-type sequences.)

**Figure S7:**
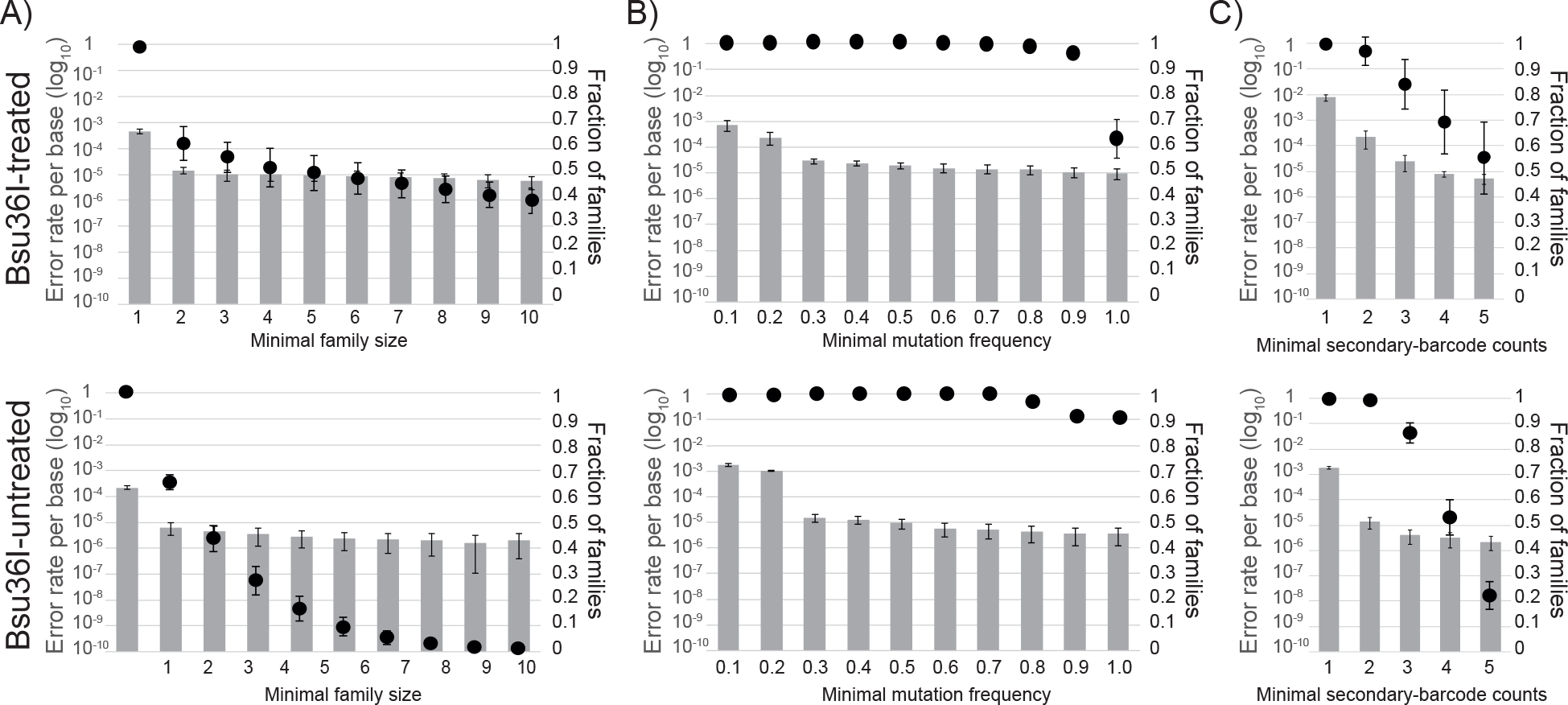
Effects of various cutoff criteria on mutation-calling accuracy. Upper row: average values from the Bsu36I-treated samples of AFR1, AFR2, EUR1, EUR2. Lower row: average values from the Bsu36I-untreated samples of the same donors. Bar graphs: error rate per base in log-10 scale (left axis) while varying each cutoff criterion alone, calculated for the 47 bp that constitute the *HBB* and *HBD* ROI-flanking sequences for the Bsu36I-treated samples and for the 54 bp that constitute the ROI and the flanking sequences for the Bsu36I-untreated samples. Chimeric *HBB*/*HBD* markers (*HBB* 9C→T and *HBD* 9T→C substitutions) were not included in the mutation count (supplementary section 9), nor was the ROI-flanking sequence mutation 14C→A that was found to be enriched by Bsu36I digestion (Fig. S13 and supplementary section 8). Dot plots: Fraction of *HBB* and *HBD* families that meet a cutoff criterion (right axis). A) The effect of increasing the family-size cutoff. Mutations present at 100% of the sequences in a primary-barcode family were selected for the mutation-rate calculation. B) The effect of increasing the mutation-frequency cutoff for families with at least four reads. C) The effect of increasing the secondary-barcode count cutoff for families with at least four reads.

**Figure S8:**
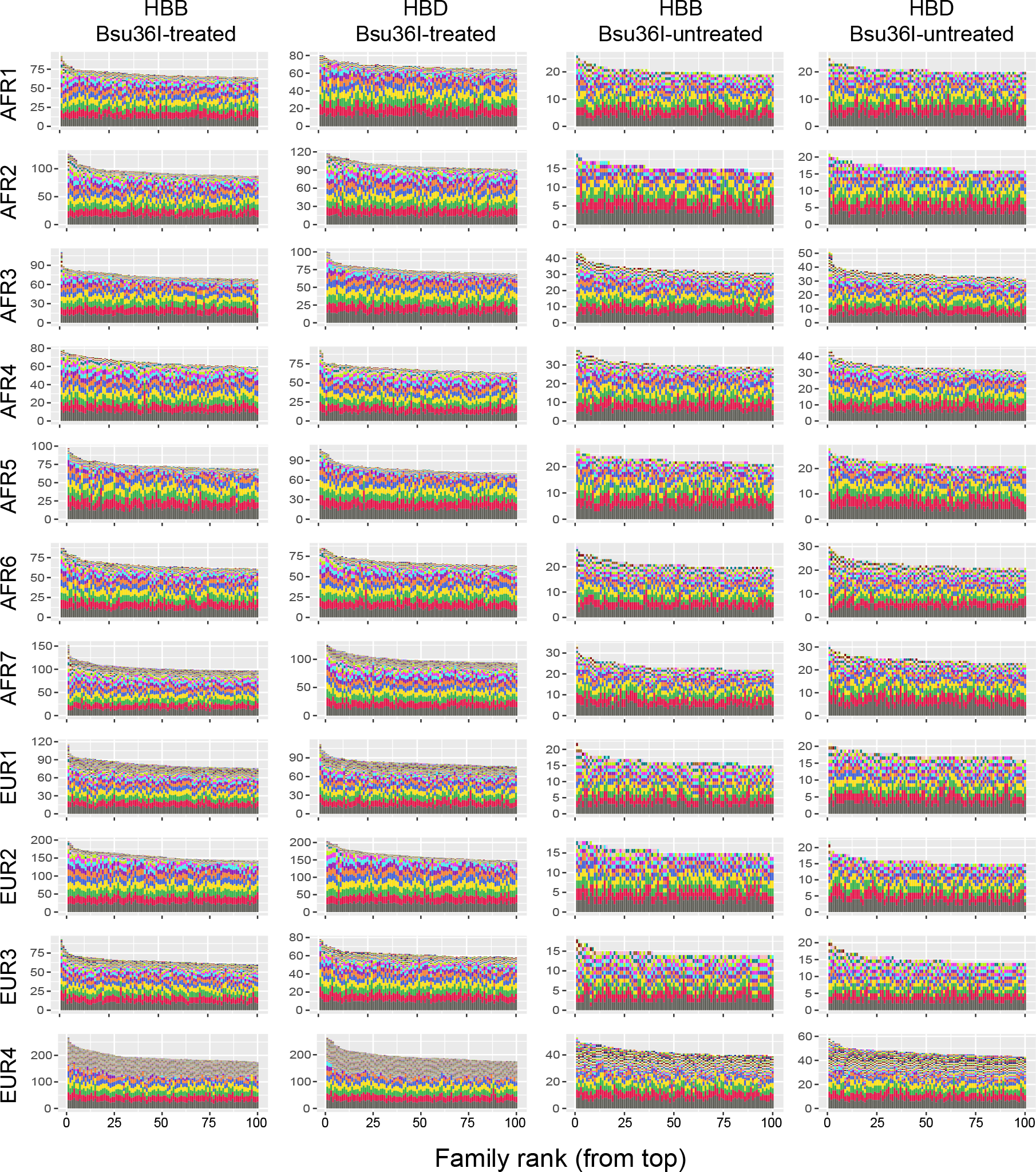
Secondary barcode distribution. Distribution of secondary-barcode counts in the top 100 families with the highest read counts. Unique secondary barcodes are marked by different colors. The family-size axes were adjusted to the families with the highest read counts.

**Figure S9:**
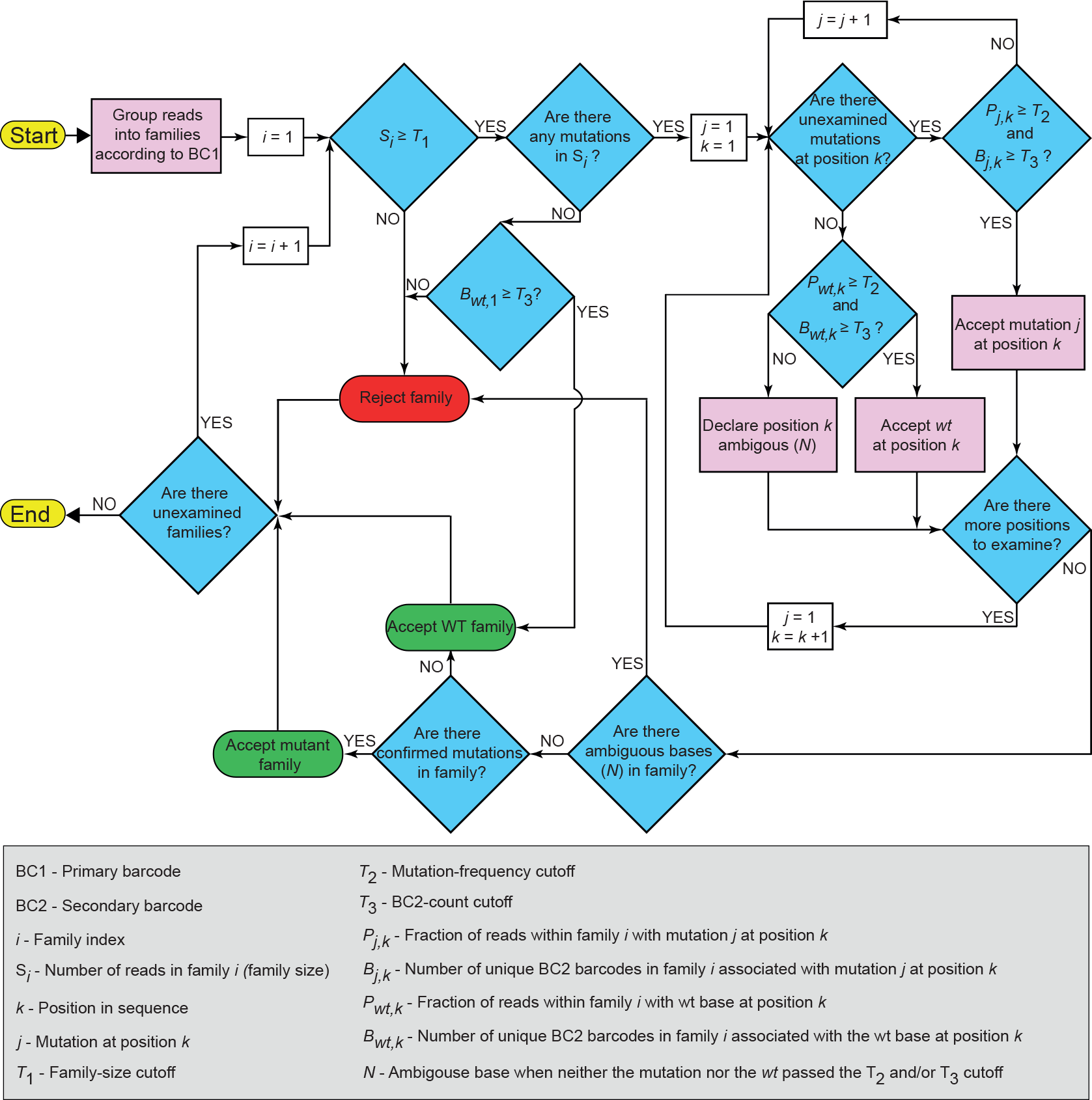
An illustration of the MEMDS computational pipeline. The workflow describes the computational analysis from the point of grouping reads into families by their shared primary barcodes, where each family represent a single target-DNA molecule, to the characterization of each family by its mutations that pass the combined cutoff criteria, if they exist. These criteria include a minimal family size of four reads (*T*_1_), a mutation frequency cutoff of at least 0.7 (*T*_2_) and the association of the particular mutation called with at least two secondary barcodes (*T*_3_). Note that for any given mutated position, in the case of failure to pass the combined cutoff criteria, the wild-type base is tested by the same conditions to validate its authenticity in an unbiased manner. If both the mutation and the wild-type base fail to meet the cutoff criteria, the base identity at that position is declared ambiguous (*N*), and the family is rejected.

**Figure S10:**
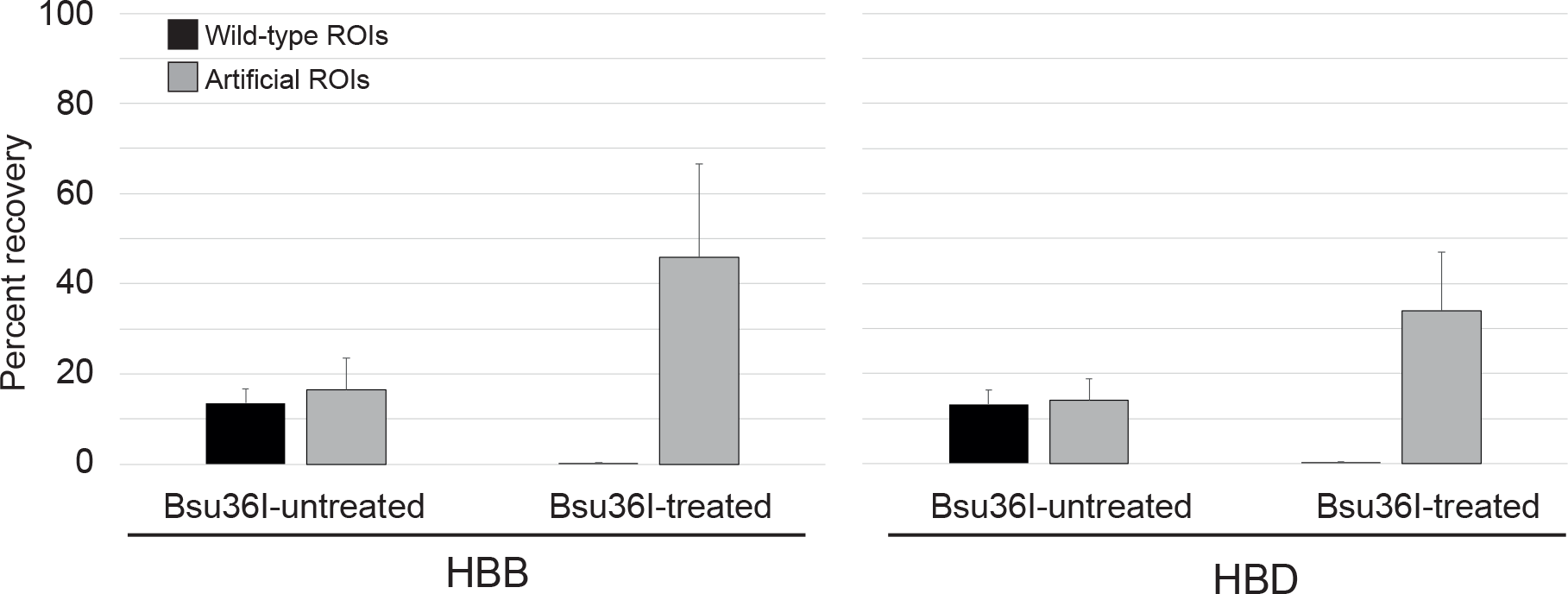
Percent recoveries of WT (genomic) sequences and artificial (plasmid) sequences by the MEMDS method. Percent recoveries of the WT and artificial target molecules were calculated using the ratios between the obtained number of families of each type and the estimated amounts of input families derived from their DNA concentration measurements.

**Figure S11:**
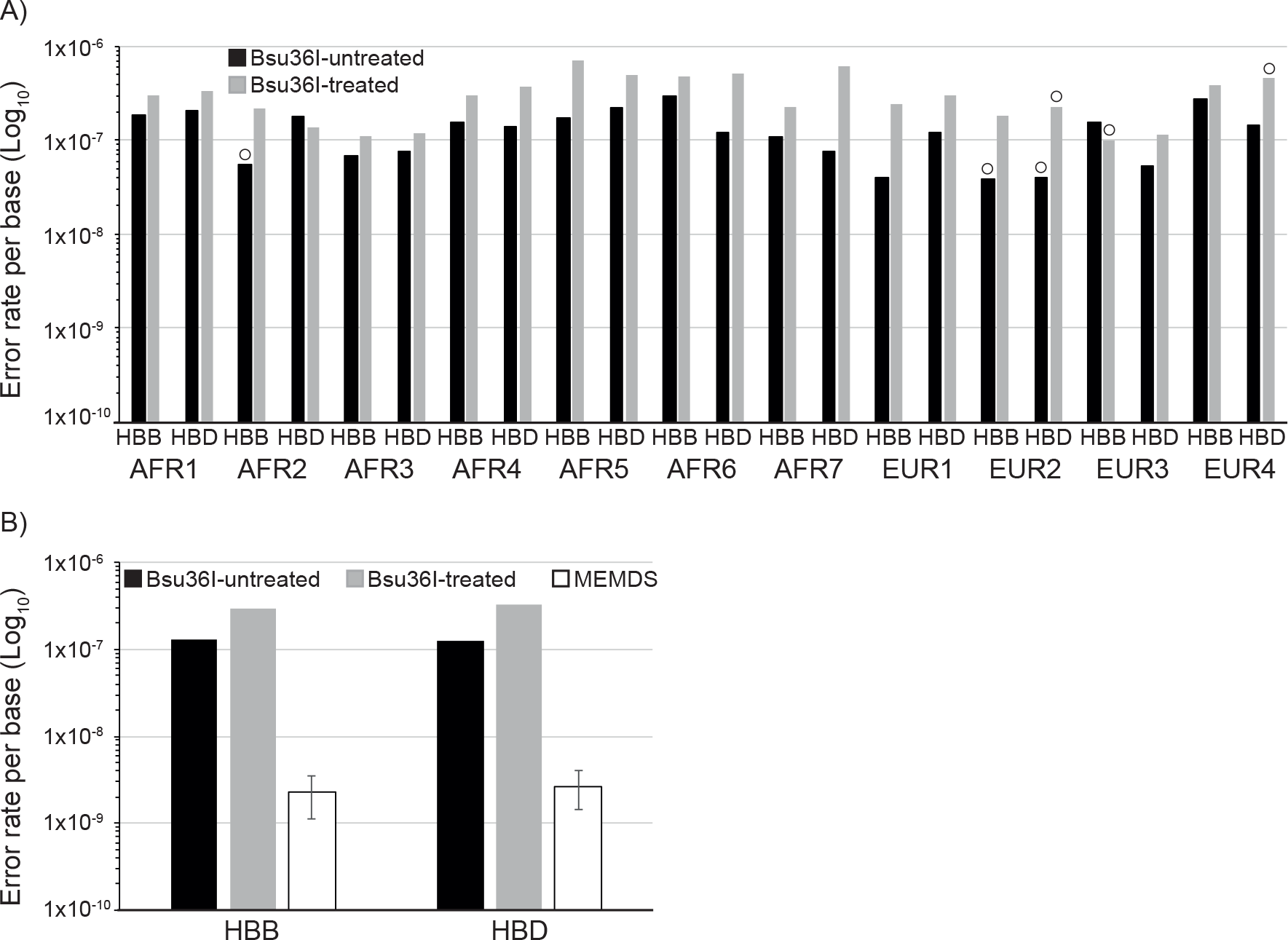
Calculated error rates. A) Per-base error rates for non G→T, C→T and C→A mutations (in the target DNA strand) were calculated for each donor for the 47 bp that include the ROI-flanking sequences in the Bsu36I-untreated samples (black bars) and Bsu36I-treated (gray bars) samples, under the stringent assumption that all mutations observed in these unenriched sequences are errors. Open circles mark samples where no non-G→T, C→T and C→A mutations were observed and the error rate calculation for these samples used a theoretical mutation count value of 1. B) The total error rate for each gene (i.e., the sum of non-G→T, C→T and C→A mutations for that gene across all donors divided by the total number of bases) calculated for the same sequences depicted in S11A. The MEMDS error rates for the 6 bases that constitute the ROI were calculated for each donor by dividing the total error rate achieved for the ROI-flanking sequences of the Bsu36I-treated samples by the relevant Bsu36I-enrichment factor.

**Figure S12:**
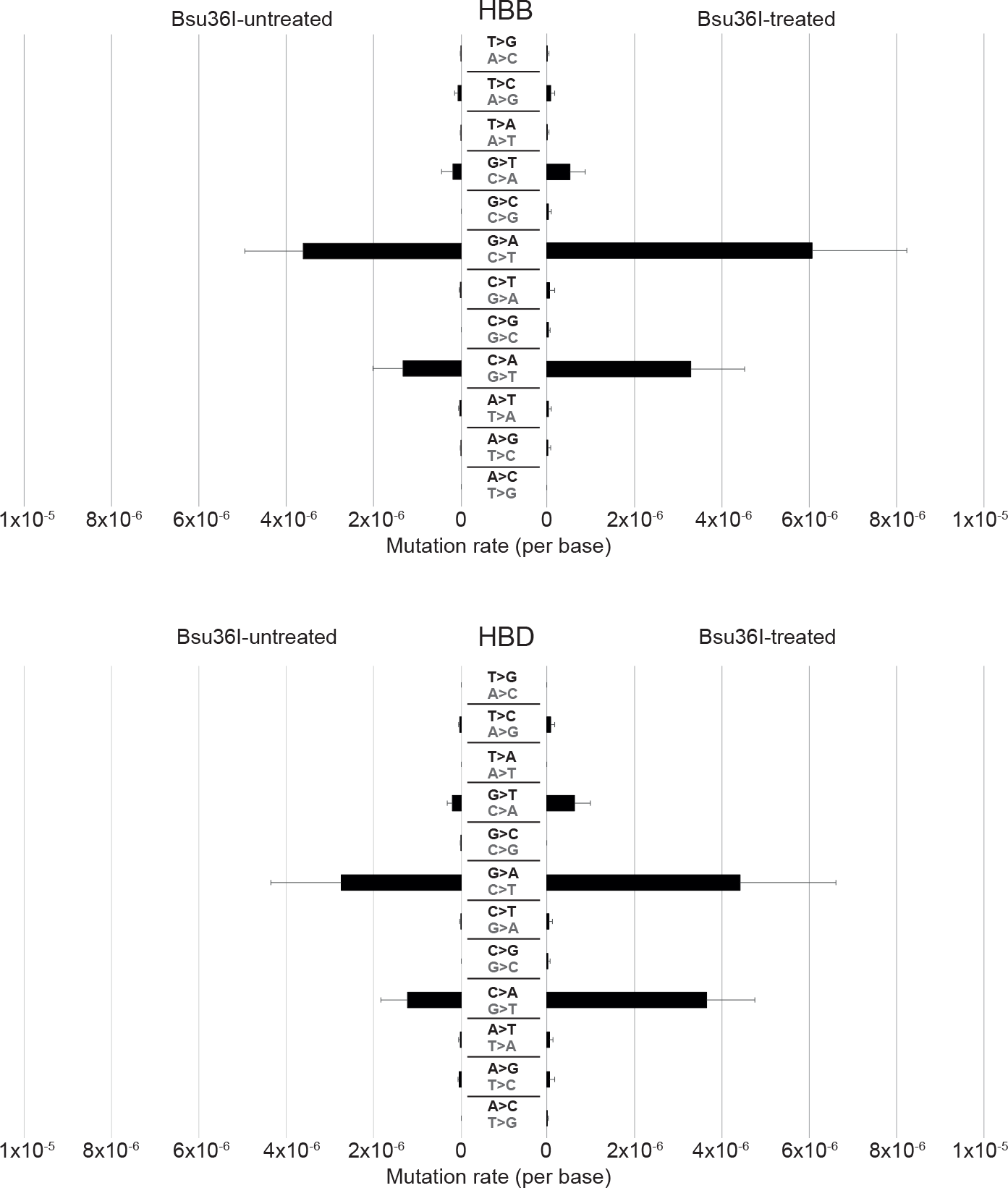
Per-type point mutation frequencies. Average point mutation frequencies were calculated for the 47 base sequence that include the ROI-flanking sequences of the Bsu36I-treated and untreated samples. The chimeric *HBB*/*HBD* markers (*HBB* 9C→T and *HBD* 9T→C substitutions) were not included in the mutation count (Fig. S13 and supplementary section 9), nor was the ROI-flanking sequence mutation 14C→A that was found to be enriched by Bsu36I digestion (Fig. S13 and supplementary section 8). In gray: mutations in the target (antisense) DNA strand. In black: the complementary mutations in the sequenced (sense) strand.

**Figure S13:**
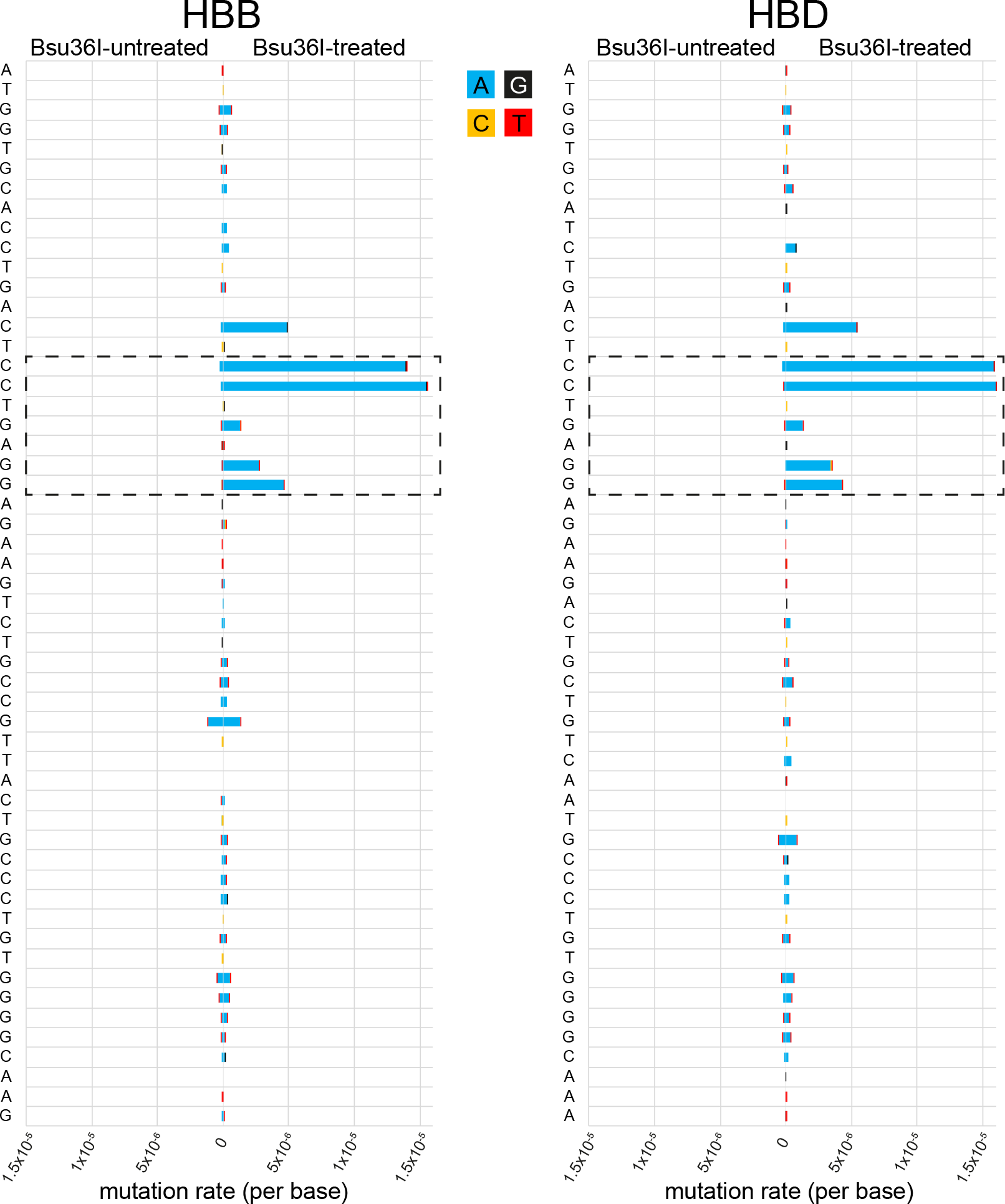
Mutation distribution in *HBB* and *HBD* sequences. Shown are the total mutation frequencies in *HBB* (left) and *HBD* (right) sequences of all 11 donors. Frequencies of mutations from the Bsu36I-treated and untreated samples are displayed in opposite directions. The Bsu36I-restriction site, six of whose seven bases define the ROI, is boxed by dashed lines. Mutation frequencies for the 9C→T substitution in *HBB* and 9T→C in *HBD* were omitted due to their possible chimeric origin. Note that both *HBB* and *HBD* sequences are shown in the sense orientation, which corresponds to the sequencing output data. Since this MEMDS experiment targeted the antisense strand of both genes, the mutations in the target DNA molecules were the reciprocals of the mutations shown here.

**Figure S14:**
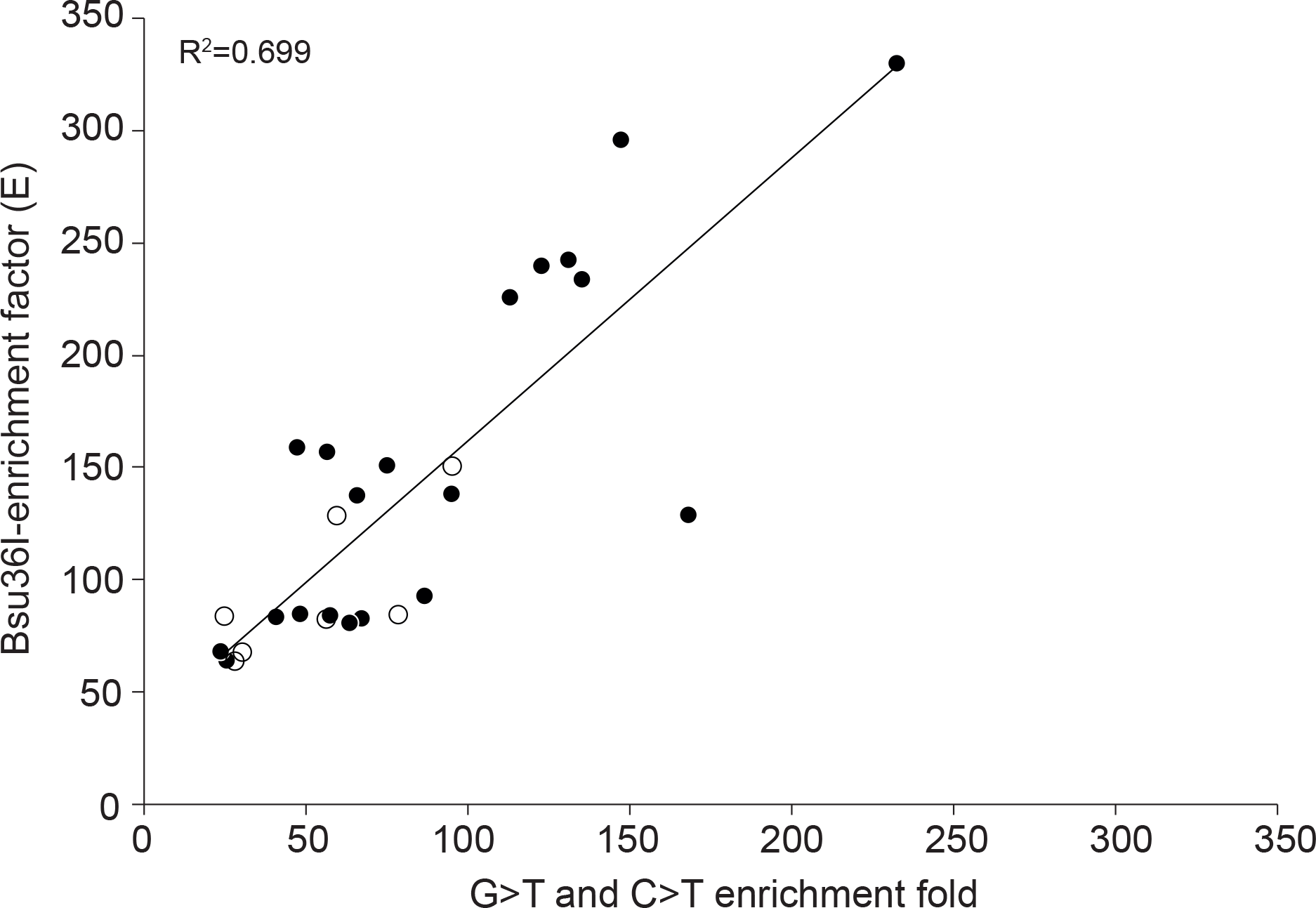
Correlation between the enrichments of target-strand G→T and C→T mutations and the Bsu36I-enrichment factors. The fold enrichment of the ROI G→T (filled circles) and C→T (open circles) mutations (C→A and G→A mutations in the sequence data, respectively) was determined by the ratio between the mutation frequencies in the ROI of the Bsu36I-treated and untreated samples. For each mutation type data is shown only for donors with at least 3 mutation counts in the ROI site.

**Table S1:** General properties of sperm samples. Semen donations were received from 8 African (all Ghanaian) and 4 Northern European donors. AFR3 is a mixture of two samples from two separate donors that were combined into one. For simplicity of analysis, we consider it as one sample of mixed Ghanaian origins.

| Donor | Donation year | Parent-1/Parent-2 origin | Age group | Ethnic group | Sperm count (millions/mL) | % motility |
| --- | --- | --- | --- | --- | --- | --- |
| Afr1 | 2018 | Ghanaian/Ghanaian | 23-28 | Akan | 46.1 | 45 |
| Afr2 | 2018 | Ghanaian/Ghanaian | 35-39 | Ewe | 27.6 | 38 |
| Afr3* | 2018 | Ghanaian/Ghanaian | 23-28 | Ga/Ewe | 24.1 | 43 |
|  | 2018 | Ghanaian/Ghanaian | 23-28 | Dagbani | 20 | 40 |
| Afr4 | 2018 | Ghanaian/Ghanaian | 18-22 | Akan | 30.5 | 46 |
| Afr5 | 2019 | Ghanaian/Ghanaian | 23-28 | Akan | 51.1 | 42 |
| Afr6 | 2019 | Ghanaian/Ghanaian | 29-34 | Akan | 42.3 | 50 |
| Afr7 | 2019 | Ghanaian/Ghanaian | 23-28 | Ewe | 45.3 | 36 |
| Eur1 | 1994 | English/German | 23-28 | Caucasian | 68.0 | 50 |
| Eur2 | 2001 | English/English-Scandinavian | 29-34 | Caucasian | 68.7 | 40 |
| Eur3 | 2005/6 | German/French-Italian | 23-28 | Caucasian | 57.2 | 35 |
| Eur4 | 1995 | English/Norwegian | 29-34 | Caucasian | 77.8 | 43 |

\* Sample contains mixed cells from two donors

**Table S2:** Numbers of rejected and approved gene families following filtration by the combined cutoff criteria. 1. Total number of families subjected to the combined cutoff criteria. 2. Families failing to meet the family size cutoff of ≤4. 3. Wild-type families with at least 4 reads that fail to meet a secondary-barcode count cutoff of ≤2. 4. Families that share their primary-barcode sequences between HBB and HBD genes of the same donor and treatment (chimeric artifacts). 5. Mutation-containing families that fail to meet a mutation-frequency cutoff of ≤0.7 or a secondary-barcode count cutoff of ≤2 for either a mutation or a wild-type base at least in one position in sequence. The combined cutoff criteria include a family size cutoff of →4, a mutation frequency cutoff of →0.7 and a secondary-barcode count cutoff of →2. Note that the numbers of rejected and approved families sum up to the total number of families.

| Donor | Treatment | Gene | Total Families | Rejected due to low counts | Rejected due to WT-secondary barcode failure | Rejected due to shared primary barcodes | Rejected due to ambiguous bases (N) | Approved by the combined cutoff criteria |
| --- | --- | --- | --- | --- | --- | --- | --- | --- |
| AFR1 | Bsu36I-treated | HBB | 577,003 | 174,577 | 33,474 | 2,865 | 6,985 | 359,102 |
|  |  | HBD | 492,594 | 189,270 | 31,969 | 2,865 | 7,390 | 261,110 |
|  | Bsu36I-untreated | HBB | 1,876,444 | 1,154,913 | 5,796 | 4,569 | 3,052 | 708,114 |
|  |  | HBD | 1,698,297 | 1,035,754 | 6,657 | 4,569 | 3,222 | 648,095 |
| AFR2 | Bsu36I-treated | HBB | 326,152 | 105,610 | 12,819 | 2,326 | 2,910 | 202,487 |
|  |  | HBD | 295,836 | 115,683 | 11,330 | 2,326 | 2,862 | 163,635 |
|  | Bsu36I-untreated | HBB | 1,784,865 | 1,374,897 | 6,232 | 2,583 | 2,136 | 399,017 |
|  |  | HBD | 1,753,711 | 1,259,296 | 5,899 | 2,583 | 2,344 | 483,589 |
| AFR3 | Bsu36I-treated | HBB | 386,700 | 126,712 | 16,860 | 3,104 | 8,990 | 231,034 |
|  |  | HBD | 349,216 | 137,830 | 18,259 | 3,104 | 5,435 | 184,588 |
|  | Bsu36I-untreated | HBB | 1,148,389 | 497,865 | 9,729 | 3,725 | 5,900 | 631,170 |
|  |  | HBD | 1,051,314 | 464,336 | 10,926 | 3,725 | 3,520 | 568,807 |
| AFR4 | Bsu36I-treated | HBB | 426,278 | 107,727 | 19,883 | 987 | 3,122 | 294,559 |
|  |  | HBD | 372,518 | 113,201 | 9,855 | 987 | 14,838 | 233,637 |
|  | Bsu36I-untreated | HBB | 1,264,724 | 521,564 | 21801 | 4,290 | 4,563 | 712,506 |
|  |  | HBD | 1,120,079 | 450,195 | 11,035 | 4,290 | 22,693 | 631,866 |
| AFR5 | Bsu36I-treated | HBB | 463,684 | 185,935 | 15,604 | 4,054 | 29,396 | 228,695 |
|  |  | HBD | 420,559 | 196,822 | 28,311 | 4,054 | 9,221 | 182,151 |
|  | Bsu36I-untreated | HBB | 1,891,157 | 1,174,471 | 7,638 | 4,031 | 19,437 | 685,580 |
|  |  | HBD | 1,746,131 | 1,103,189 | 14,760 | 4,031 | 5,622 | 618,529 |
| AFR6 | Bsu36I-treated | HBB | 573,258 | 208,356 | 29,491 | 3,613 | 7,612 | 324,186 |
|  |  | HBD | 531,781 | 200,664 | 22,825 | 3,613 | 5,782 | 298,897 |
|  | Bsu36I-untreated | HBB | 1,575,956 | 1,014,909 | 3,497 | 3,292 | 3,864 | 550,394 |
|  |  | HBD | 1,481,666 | 903,810 | 3,176 | 3,292 | 3,305 | 568,083 |
| AFR7 | Bsu36I-treated | HBB | 574,043 | 212,633 | 10,969 | 32,028 | 3,850 | 314,563 |
|  |  | HBD | 541,285 | 234,389 | 11,208 | 32,028 | 3,758 | 259,902 |
|  | Bsu36I-untreated | HBB | 1,699,015 | 1,043,765 | 10,971 | 4,513 | 5,811 | 633,955 |
|  |  | HBD | 1,555,197 | 932,933 | 9,599 | 4,513 | 5,004 | 603,148 |
| EUR1 | Bsu36I-treated | HBB | 331,608 | 126,256 | 3,737 | 19,141 | 1,229 | 181,245 |
|  |  | HBD | 311,712 | 139,262 | 3,252 | 19,141 | 1,422 | 148,635 |
|  | Bsu36I-untreated | HBB | 2,144,051 | 1,589,762 | 2,462 | 4,199 | 2,161 | 545,467 |
|  |  | HBD | 1,936,890 | 1,388,844 | 2,327 | 4,199 | 2,317 | 539,203 |
| EUR2 | Bsu36I-treated | HBB | 228,770 | 89,463 | 7,512 | 4,692 | 2,623 | 124,480 |
|  |  | HBD | 210,577 | 97,700 | 6,413 | 4,692 | 2,734 | 99,038 |
|  | Bsu36I-untreated | HBB | 2,392,244 | 1,827,090 | 3,029 | 4,056 | 1,565 | 556,504 |
|  |  | HBD | 2,172,098 | 1,619,546 | 3,392 | 4,056 | 1,725 | 543,379 |
| EUR3 | Bsu36I-treated | HBB | 418,458 | 161,339 | 3236 | 7,226 | 7,428 | 239,229 |
|  |  | HBD | 393,302 | 172,746 | 7,036 | 7,226 | 3,384 | 202,910 |
|  | Bsu36I-untreated | HBB | 2,335,295 | 1,888,649 | 422 | 2,698 | 11,427 | 432,099 |
|  |  | HBD | 2,143,059 | 1,712,093 | 778 | 2,698 | 3,942 | 423,548 |
| EUR4 | Bsu36I-treated | HBB | 144,833 | 43,533 | 20,359 | 36 | 13,548 | 67,357 |
|  |  | HBD | 122,042 | 38,710 | 18,175 | 36 | 11,712 | 53,409 |
|  | Bsu36I-untreated | HBB | 731,846 | 242,928 | 122,442 | 1,102 | 26,355 | 339,019 |
|  |  | HBD | 649,808 | 215,711 | 98,264 | 1,102 | 19,597 | 315,134 |

**Table S3:** Values for the calculation of Bsu36I-enrichment factors and numbers of scanned target DNA sequences. Number of families with no mutations at the ROI that passed the combined-cutoff criteria. 2. Number of families of plasmid DNA (two plasmids per a tested gene) that passed the combined-cutoff criteria. 3. Bsu36I enrichment factor, calculated by the volumes and family counts depicted in the table using the formula shown in supplementary section 2. 4. For the Bsu36I-treated samples, includes both the sequenced and the Bsu36I-digested WT target sequences, which are computed by multiplying the number of WT families by the enrichment factor. For the Bsu36I-untreated samples, this number includes only the sequenced families.

| Donor | Treatment | Donor DNA Volume taken | Plasmid DNA Volume taken | Gene | WT families passing cutoff<sup>1</sup> | Plasmid families passing cutoff<sup>2</sup> | Enrichment factor<sup>3</sup> | Scanned target-sequences<sup>4</sup> |
| --- | --- | --- | --- | --- | --- | --- | --- | --- |
| **Afr1**<br>380 ng/μl | Bsu36I-treated | 700 μl ($V_{Se}$) | 20 μl ($V_{Re}$) | HBB | 350,937 ($S_e^f$) | 3,657; 3,899 ($R_e^f$) | 63.60 | 22,319,593 |
| | | | | HBD | 254,507 ($S_e^f$) | 2,939; 3,158 ($R_e^f$) | 67.04 | 17,062,149 |
| | Bsu36I-untreated | 35 μl ($V_{Sc}$) | 120 μl ($V_{Rc}$) | HBB | 680,429 ($S_c^f$) | 12,275; 15,367 ($R_c^f$) | 1.00 | 680,429 |
| | | | | HBD | 621,399 ($S_c^f$) | 12,156; 14,492 ($R_c^f$) | 1.00 | 621,399 |
| **Afr2**<br>591 ng/μl | Bsu36I-treated | 338 μl ($V_{Se}$) | 20 μl ($V_{Re}$) | HBB | 194,704 ($S_e^f$) | 3,483; 3,803 ($R_e^f$) | 158.56 | 30,872,266 |
| | | | | HBD | 157,003 ($S_e^f$) | 3,057; 3,200 ($R_e^f$) | 156.05 | 24,500,318 |
| | Bsu36I-untreated | 16.9 μl ($V_{Sc}$) | 120 μl ($V_{Rc}$) | HBB | 388,011 ($S_c^f$) | 4,682; 6,307 ($R_c^f$) | 1.00 | 388,011 |
| | | | | HBD | 469,182 ($S_c^f$) | 6,077; 8,302 ($R_c^f$) | 1.00 | 469,182 |
| **Afr3**<br>355 ng/μl | Bsu36I-treated | 744 μl ($V_{Se}$) | 20 μl ($V_{Re}$) | HBB | 194,315 ($S_e^f$) | 4,179; 3917 ($R_e^f$) | 150.09 | 29,164,431 |
| | | | | HBD | 177,422 ($S_e^f$) | 3,463; 3,227 ($R_e^f$) | 128.13 | 22,696,345 |
| | Bsu36I-untreated | 37.2 μl ($V_{Sc}$) | 120 μl ($V_{Rc}$) | HBB | 610,803 ($S_c^f$) | 10,760; 9,587 ($R_c^f$) | 1.00 | 610,803 |
| | | | | HBD | 549,390 ($S_c^f$) | 10,267; 9,135 ($R_c^f$) | 1.00 | 549,390 |
| **Afr4**<br>355 ng/μl | Bsu36I-treated | 720 μl ($V_{Se}$) | 20 μl ($V_{Re}$) | HBB | 286,939 ($S_e^f$) | 3,561; 3,341 ($R_e^f$) | 82.85 | 23,773,196 |
| | | | | HBD | 227,197 ($S_e^f$) | 2,931; 2,908 ($R_e^f$) | 83.65 | 19,005,979 |
| | Bsu36I-untreated | 36 μl ($V_{Sc}$) | 120 μl ($V_{Rc}$) | HBB | 688,478 ($S_c^f$) | 12,538; 11,448 ($R_c^f$) | 1.00 | 688,478 |
| | | | | HBD | 609,365 ($S_c^f$) | 11,616; 10,849 ($R_c^f$) | 1.00 | 609,365 |
| **Afr5**<br>445 ng/μl | Bsu36I-treated | 600 μl ($V_{Se}$) | 20 μl ($V_{Re}$) | HBB | 211,556 ($S_e^f$) | 9,274; 7,321 ($R_e^f$) | 91.87 | 19,436,031 |
| | | | | HBD | 172,778 ($S_e^f$) | 5,007; 3,830 ($R_e^f$) | 83.92 | 14,500,198 |
| | Bsu36I-untreated | 30 μl ($V_{Sc}$) | 120 μl ($V_{Rc}$) | HBB | 621,848 ($S_c^f$) | 37,351; 26,363 ($R_c^f$) | 1.00 | 621,848 |
| | | | | HBD | 576,349 ($S_c^f$) | 25,761; 16,389 ($R_c^f$) | 1.00 | 576,349 |
| **Afr6**<br>894 ng/μl | Bsu36I-treated | 320 μl ($V_{Se}$) | 20 μl ($V_{Re}$) | HBB | 305,108 ($S_e^f$) | 9,731; 8,726 ($R_e^f$) | 82.10 | 25,048,983 |
| | | | | HBD | 285,210 ($S_e^f$) | 7,152; 5,863 ($R_e^f$) | 80.18 | 22,867,577 |
| | Bsu36I-untreated | 16 μl ($V_{Sc}$) | 120 μl ($V_{Rc}$) | HBB | 505,664 ($S_c^f$) | 25,122; 19,589 ($R_c^f$) | 1.00 | 505,664 |
| | | | | HBD | 531,747 ($S_c^f$) | 20,863; 15,454 ($R_c^f$) | 1.00 | 531,747 |
| **Afr7**<br>694 ng/μl | Bsu36I-treated | 360 μl ($V_{Se}$) | 20 μl ($V_{Re}$) | HBB | 284,362 ($S_e^f$) | 15,848; 13,647 ($R_e^f$) | 137.55 | 39,114,652 |
| | | | | HBD | 239,830 ($S_e^f$) | 10,693; 8,789 ($R_e^f$) | 136.97 | 32,848,562 |
| | Bsu36I-untreated | 18 μl ($V_{Sc}$) | 120 μl ($V_{Rc}$) | HBB | 581,338 ($S_c^f$) | 29,950; 22,654 ($R_c^f$) | 1.00 | 581,338 |
| | | | | HBD | 563,058 ($S_c^f$) | 22,790; 17,283 ($R_c^f$) | 1.00 | 563,058 |
| **Eur1**<br>446 ng/μl | Bsu36I-treated | 592 μl ($V_{Se}$) | 20 μl ($V_{Re}$) | HBB | 173,119 ($S_e^f$) | 3,696; 4,081 ($R_e^f$) | 225.50 | 39,038,335 |
| | | | | HBD | 141,078 ($S_e^f$) | 3,460; 3,731 ($R_e^f$) | 239.46 | 33,782,538 |
| | Bsu36I-untreated | 29.6 μl ($V_{Sc}$) | 120 μl ($V_{Rc}$) | HBB | 532,724 ($S_c^f$) | 5,431; 7,304 ($R_c^f$) | 1.00 | 532,724 |
| | | | | HBD | 525,762 ($S_c^f$) | 5,951; 7,479 ($R_c^f$) | 1.00 | 525,762 |
| **Eur2**<br>339 ng/μl | Bsu36I-treated | 780 μl ($V_{Se}$) | 20 μl ($V_{Re}$) | HBB | 118,015 ($S_e^f$) | 3,020; 3,231 ($R_e^f$) | 329.27 | 38,858,799 |
| | | | | HBD | 93,487 ($S_e^f$) | 2,669; 2,712 ($R_e^f$) | 340.41 | 31,823,910 |
| | Bsu36I-untreated | 39 μl ($V_{Sc}$) | 120 μl ($V_{Rc}$) | HBB | 545,961 ($S_c^f$) | 4,499; 6,040 ($R_c^f$) | 1.00 | 545,961 |
| | | | | HBD | 532,567 ($S_c^f$) | 4,788; 6,018 ($R_c^f$) | 1.00 | 532,567 |
| **Eur3**<br>633 ng/μl | Bsu36I-treated | 440 μl ($V_{Se}$) | 20 μl ($V_{Re}$) | HBB | 210,009 ($S_e^f$) | 15,676; 13,036 ($R_e^f$) | 233.12 | 48,956,676 |
| | | | | HBD | 183,054 ($S_e^f$) | 10,589; 8,828 ($R_e^f$) | 242.05 | 44,308,361 |
| | Bsu36I-untreated | 22 μl ($V_{Sc}$) | 120 μl ($V_{Rc}$) | HBB | 403,681 ($S_c^f$) | 17,836; 10,574 ($R_c^f$) | 1.00 | 403,681 |
| | | | | HBD | 402,381 ($S_c^f$) | 13,357; 7,803 ($R_c^f$) | 1.00 | 402,381 |
| **Eur4**<br>376 ng/μl | Bsu36I-treated | 720 μl ($V_{Se}$) | 20 μl ($V_{Re}$) | HBB | 55,898 ($S_e^f$) | 6,135; 5,215 ($R_e^f$) | 295.65 | 16,526,517 |
| | | | | HBD | 45,832 ($S_e^f$) | 4,077; 3,403 ($R_e^f$) | 318.23 | 14,585,464 |
| | Bsu36I-untreated | 36 μl ($V_{Sc}$) | 120 μl ($V_{Rc}$) | HBB | 313,203 ($S_c^f$) | 13,410; 12,402 ($R_c^f$) | 1.00 | 313,203 |
| | | | | HBD | 296,860 ($S_c^f$) | 9,937; 8,332 ($R_c^f$) | 1.00 | 296,860 |

**Table S4:** Fold change of observed *de novo* rates from the genome-wide average (GWA) rates for point mutations in the African *HBB* ROI. Names of clinically known variants were taken from the Globin Gene Server database (*25,26*). GWA rates were calculated as described in section 10 based on a subset of *de novo* mutations with phasing information taken from genome-wide family sequencing studies (*55*) as well as based on relative frequencies of Extremely Rare Variants (ERVs) for the 3-mer, 5-mer and 7-mer genetic contexts (*61*). The significance of the deviations of the mutation-specific origination rates observed here from the GWA rates is not affected by taking into account the local genetic context in three out of four cases, in accord with the fact that adjustments to the GWA rates based on context are minor compared to the variation in mutation-specific origination rates observed here. Out of 12 point mutation types studied here, the mutation-specific origination rate of the HbS mutation deviates by far the most from its GWA rate, even when taking into account the local genetic context. This effect is only strengthened in the larger contexts, as the origination rate of the HbS mutation deviates by ∼35 from its GWA calculated based on *de novo* mutation studies and by ∼38, ∼40 and ∼51 from its GWA calculated based on ERVs for the 3-mer, 5-mer and 7-mer contexts, respectively.

| Change description |  | *De novo* mutation-based |  | ERV-based, 3-mer |  | ERV-based, 5-mer |  | ERV-based, 7-mer |  |
| --- | --- | --- | --- | --- | --- | --- | --- | --- | --- |
| Mutation | Name (HbVar) | GWA rate (x10<sup>-8</sup>) | Fold change | GWA rate (x10<sup>-8</sup>) | Fold change | GWA rate (x10<sup>-8</sup>) | Fold change | GWA rate (x10<sup>-8</sup>) | Fold change |
| 16C>G | Hb Gorwihl | 0.21 | *7.4 | 0.27 | 5.8 | 0.28 | 5.7 | 0.36 | 4.4 |
| 16C>T | Hb Tyne | 0.73 | 2.2 | 0.78 | 2.0 | 0.97 | 1.6 | 0.79 | 2.0 |
| 17C>G | Hb Warwickshire | 0.21 | ***12.3 | 0.27 | ***9.7 | 0.17 | ***15.1 | 0.16 | ***16.4 |
| 17C>T | Hb Aix-les-Bains | 0.73 | 2.2 | 0.84 | 1.9 | 0.66 | 2.4 | 0.60 | 2.6 |
| 18T>G |  | 0.17 | 3.1 | 0.17 | 3.1 | 0.16 | 3.3 | 0.16 | 3.2 |
| 18T>A |  | 0.14 | 3.8 | 0.13 | 4.0 | 0.13 | 4.1 | 0.13 | 3.9 |
| 18T>C |  | 0.59 | 1.8 | 0.55 | 1.9 | 0.59 | 1.8 | 0.76 | 1.4 |
| 20A>G | Hb Lavagna | 0.59 | **4.4 | 0.37 | **7.0 | 0.29 | ***9.1 | 0.28 | ***9.5 |
| 20A>T | HbS | 0.14 | ****34.4 | 0.13 | ****37.6 | 0.12 | ****40.4 | 0.09 | ****50.7 |
| 20A>C | Hb G-Makassar | 0.17 | 3.1 | 0.13 | 4.1 | 0.12 | 4.4 | 0.12 | 4.5 |
| 21G>C |  | 0.21 | 2.5 | 0.27 | 1.9 | 0.22 | 2.4 | 0.18 | 3.0 |
| 22G>C | Hb Bellevue III | 0.21 | 0 | 0.27 | 0.0 | 0.28 | 0.0 | 0.20 | 0.0 |

\* p = 0.0082
\*\* p < 0.0060
\*\*\* p < 3x10<sup>-4</sup>
\*\*\*\* p < 10<sup>-10</sup>

## References

1. Hodgkinson A, Eyre-Walker A (2011) Variation in the mutation rate across mammalian genomes. Nat Rev Genet 12:756–766.

2. Rahbari R, et al. (2016) Timing, rates and spectra of human germline mutation. Nat Genet 48(2):126.

3. Carlson J, et al. (2018) Extremely rare variants reveal patterns of germline mutation rate heterogeneity in humans. Nat Commun 9(1):3753.

4. Gojobori T, Li WH, Graur D (1982) Patterns of nucleotide substitution in pseudogenes and functional genes. Journal of Molecular Evolution 18(5):360–369.

5. Bulmer M (1986) Neighboring base effects on substitution rates in pseudogenes. Molecular Biology and Evolution 3(4):322–329.

6. Blake R, Hess ST, Nicholson-Tuell J (1992) The influence of nearest neighbors on the rate and pattern of spontaneous point mutations. Journal of Molecular Evolution 34(3):189– 200.

7. Hwang DG, Green P (2004) Bayesian markov chain monte carlo sequence analysis reveals varying neutral substitution patterns in mammalian evolution. Proceedings of the National Academy of Sciences 101(39):13994–14001.

8. Wolfe KH, Sharp PM, Li WH (1989) Mutation rates differ among regions of the mammalian genome. Nature 337(6204):283–285.

9. Matassi G, Sharp PM, Gautier C (1999) Chromosomal location effects on gene sequence evolution in mammals. Current Biology 9(15):786–791.

10. Lercher MJ, Williams EJ, Hurst LD (2001) Local similarity in evolutionary rates extends over whole chromosomes in human-rodent and mouse-rat comparisons: implications for understanding the mechanistic basis of the male mutation bias. Molecular Biology and Evolution 18(11):2032–2039.

11. Ellegren H, Smith NG, Webster MT (2003) Mutation rate variation in the mammalian genome. Current Opinion in Genetics & Development 13(6):562–568.

12. Campbell CD, et al. (2012) Estimating the human mutation rate using autozygosity in a founder population. Nature Genetics 44(11):1277–1281.

13. Michaelson JJ, et al. (2012) Whole-genome sequencing in autism identifies hot spots for de novo germline mutation. Cell 151(7):1431–1442.

14. Francioli LC, et al. (2015) Genome-wide patterns and properties of *de novo* mutations in humans. Nat Genet 47(7):822.

15. Nachman MW, Crowell SL (2000) Estimate of the mutation rate per nucleotide in humans. Genetics 156(1):297–304.

16. Haldane JBS (1949) The rate of mutation of human genes. Hereditas 35(S1):267–273.

17. Vogel F, Motulsky A (1997) Human genetics: problems and approaches. (Springer-Verlag, Berlin).

18. Kondrashov AS (2003) Direct estimates of human per nucleotide mutation rates at 20 loci causing Mendelian diseases. Hum Mutat 21(1):12–27.

19. Lek M, et al. (2016) Analysis of protein-coding genetic variation in 60,706 humans. Nature 536(7616):285–291.

20. Harpak A, Bhaskar A, Pritchard J (2016) Mutation rate variation is a primary determinant of the distribution of allele frequencies in humans. PLoS Genet 12(12):e1006489.

21. Mathieson I, Reich D (2017) Differences in the rare variant spectrum among human populations. PLoS Genet 13(2):e1006581.

22. Inoue K, et al. (2001) The 1.4-Mb CMT1A duplication/HNPP deletion genomic region reveals unique genome architectural features and provides insights into the recent evolution of new genes. Genome Res 11(6):1018–1033.

23. Crow KD, Amemiya CT, Roth J, Wagner GP (2009) Hypermutability of *HoxA13A* and functional divergence from its paralog are associated with the origin of a novel developmental feature in zebrafish and related taxa (Cypriniformes). Evolution 63(6):1574–1592.

24. Dumas LJ, et al. (2012) Duf1220-domain copy number implicated in human brain-size pathology and evolution. Am J Hum Genet 91(3):444–454.

25. Losos JB (2017) Improbable Destinies: Fate, Chance, and the Future of Evolution. (Penguin).

26. Xie KT, et al. (2019) DNA fragility in the parallel evolution of pelvic reduction in stickle-back fish. Science 363(6422):81–84.

27. Kratochwil CF, Liang Y, Urban S, Torres-Dowdall J, Meyer A (2019) Evolutionary dynamics of structural variation at a key locus for color pattern diversification in cichlid fishes. Genome Biol Evol 11(12):3452–3465.

28. Kratochwil CF, Meyer A (2019) Fragile DNA contributes to repeated evolution. Genome Biol 20(1):39.

29. Lind PA (2019) Repeatability and predictability in experimental evolution in Evolution, Origin of Life, Concepts and Methods, ed. Pontarotti P. (Springer), pp. 57–83.

30. Lupski JR (1998) Genomic disorders: structural features of the genome can lead to DNA rearrangements and human disease traits. Trends Genet 14(10):417–422.

31. McClellan J, King MC (2010) Genetic heterogeneity in human disease. Cell 141(2):210– 217.

32. Veltman JA, Brunner HG (2012) *de novo* mutations in human genetic disease. Nat Rev Genet 13(8):565–575.

33. Shendure J, Akey JM (2015) The origins, determinants, and consequences of human mutations. Science 349(6255):1478–1483.

34. Goldmann JM, et al. (2016) Parent-of-origin-specific signatures of *de novo* mutations. Nat Genet 48(8):935.

35. Roach JC, et al. (2010) Analysis of genetic inheritance in a family quartet by whole-genome sequencing. Science 328(5978):636–639.

36. Conrad DF, et al. (2011) Variation in genome-wide mutation rates within and between human families. Nat Genet 43(7):712.

37. Salk JJ, Schmitt MW, Loeb LA (2018) Enhancing the accuracy of Next-Generation Sequencing for detecting rare and subclonal mutations. Nature Reviews Genetics 19(5):269.

38. Pauling L, Itano HA, Singer SJ, Wells IC (1949) Sickle-cell anemia, a molecular disease. Science 110:543–548.

39. Ingram V (1957) Gene mutations in human hemoglobin: The chemical difference between normal and sickle hemoglobin. Nature 180:326–328.

40. Allison AC (1954) Protection afforded by sickle-cell trait against subtertian malarial infection. BMJ Brit Med J 1:290–294.

41. Hartl DL, Clark AG (2007) Principles of Population Genetics. (Sinauer Associates), 4th edition.

42. Cavalli-Sforza LL, Feldman MW (2003) The application of molecular genetic approaches to the study of human evolution. Nat Genet 33(3):266–275.

43. Feng Z, Smith D, McKenzie F, Levin S (2004) Coupling ecology and evolution: malaria and the S-gene across time scales. Math Biosci 189(1):1–19.

44. Piel FB, et al. (2010) Global distribution of the sickle cell gene and geographical confirmation of the malaria hypothesis. Nat Commun 1:104.

45. Flint J, Harding RM, Boyce AJ, Clegg JB (1998) The population genetics of the haemoglobinopathies. Baillière’s Clin Haem 11:1–51.

46. Kwiatkowski DP (2005) How malaria has affected the human genome and what human genetics can teach us about malaria. Am J Hum Genet 77(2):171–192.

47. Carter R, Mendis K (2002) Evolutionary and historical aspects of the burden of malaria. Clin Microbiol Rev 15(4):564–94.

48. Hardison RC, et al. (2002) HbVar: a relational database of human hemoglobin variants and thalassemia mutations at the globin gene server. Hum Mutat 19(3):225–233.

49. Hardison R, Miller W (2002) “Globin Gene Server,” http://globin.cse.psu.edu/ Accessed 10/5/2019.

50. Steinberg M, Adams JI (1991) Hemoglobin A2: origin, evolution, and aftermath. Blood 78(9):2165–2177.

51. Jee J, et al. (2016) Rates and mechanisms of bacterial mutagenesis from maximum-depth sequencing. Nature 534(7609):693.

52. Arbeithuber B, Makova KD, Tiemann-Boege I (2016) Artifactual mutations resulting from DNA lesions limit detection levels in ultrasensitive sequencing applications. DNA Res 23(6):547–559.

53. Harris K (2015) Evidence for recent, population-specific evolution of the human mutation rate. Proceedings of the National Academy of Sciences 112(11):3439–3444.

54. Harris K, Pritchard JK (2017) Rapid evolution of the human mutation spectrum. Elife 6.

55. Gu X, Li WH (1995) The size distribution of insertions and deletions in human and rodent pseudogenes suggests the logarithmic gap penalty for sequence alignment. Journal of Molecular Evolution 40(4):464–473.

56. Lynch M (2010) Rate, molecular spectrum, and consequences of human mutation. P Natl Acad Sci USA 107(3):961–968.

57. Gu W, Zhang F, Lupski JR (2008) Mechanisms for human genomic rearrangements. Patho-Genetics 1:4.

58. Zhang F, Carvalho CMB, Lupski JR (2009) Complex human chromosomal and genomic rearrangements. Trends Genet 25:298–307.

59. Livnat A (2013) Interaction-based evolution: how natural selection and nonrandom mutation work together. Biol Direct 8(1):24.

60. Livnat A (2017) Simplification, innateness, and the absorption of meaning from context: how novelty arises from gradual network evolution. Evol Biol 44(2):145–189.

61. Zhang Z, Gerstein M (2003) Patterns of nucleotide substitution, insertion and deletion in the human genome inferred from pseudogenes. Nucleic Acids Res 31(18):5338–5348.

62. Hodgkinson A, Ladoukakis E, Eyre-Walker A (2009) Cryptic variation in the human mutation rate. PLoS Biol 7(2).

63. Leigh Jr EG (1970) Natural selection and mutability. Am Nat 104:301–305.

64. Martincorena I, Luscombe NM (2013) Non-random mutation: the evolution of targeted hypermutation and hypomutation. Bioessays 35(2):123–130.

65. Feldman MW, Liberman U (1986) An evolutionary reduction principle for genetic modifiers. P Natl Acad Sci USA 83:4824–4827.

66. Altenberg L, Liberman U, Feldman MW (2017) Unified reduction principle for the evolution of mutation, migration, and recombination. Proceedings of the National Academy of Sciences 114(12):E2392–E2400.

67. Cairns J, Overbaugh J, Miller S (1988) The origin of mutants. Nature 335(6186):142–145.

68. Moxon ER, Rainey PB, Nowak MA, Lenski RE (1994) Adaptive evolution of highly mutable loci in pathogenic bacteria. Current biology 4(1):24–33.

69. Nishikura K (2010) Functions and regulation of RNA editing by ADAR deaminases. Annual Review of Biochemistry 79:321–349.

70. Klose RJ, Bird AP (2006) Genomic DNA methylation: the mark and its mediators. Trends Biochem Sci 31(2):89–97.

71. Xu G, Zhang J (2014) Human coding RNA editing is generally nonadaptive. Proceedings of the National Academy of Sciences USA 111(10):3769–3774.

72. Popitsch N, et al. (2020) A-to-I RNA editing uncovers hidden signals of adaptive genome evolution in animals. Genome Biology and Evolution 12(4):345–357.

73. Luria SE, Delbrück M (1943) Mutations of bacteria from virus sensitivity to virus resistance. Genetics 28(6):491.

74. CDC Division of Parasitic Diseases & Malaria (2019) “Where Malaria Occurs,” http://www.cdc.gov/malaria/about/distribution.html Accessed 1/24/2019.

75. Kennedy SR, et al. (2014) Detecting ultralow-frequency mutations by Duplex Sequencing. Nat Protoc 9(11):2586.

## References

1. Hestand MS, Van Houdt J, Cristofoli F, Vermeesch JR (2016) Polymerase specific error rates and profiles identified by single molecule sequencing. Mutat Res 784:39–45.

2. Lee DF, Lu J, Chang S, Loparo JJ, Xie XS (2016) Mapping DNA polymerase errors by single-molecule sequencing. Nucleic Acids Res 44(13):e118–e118.

3. Potapov V, Ong JL (2017) Examining sources of error in PCR by single-molecule sequencing. PloS One 12(1).

4. Fox EJ, Reid-Bayliss KS, Emond MJ, Loeb LA (2014) Accuracy of Next Generation Sequencing platforms. Next Gener Seq Appl 1.

5. Ma X, et al. (2019) Analysis of error profiles in deep Next-Generation Sequencing data. Genome Biol 20(1):50.

6. Cibulskis K, et al. (2013) Sensitive detection of somatic point mutations in impure and heterogeneous cancer samples. Nat Biotechnol 31(3):213.

7. Casbon JA, Osborne RJ, Brenner S, Lichtenstein CP (2011) A method for counting PCR template molecules with application to next-generation sequencing. Nucleic Acids Res 39(12):e81–e81.

8. Kinde I, Wu J, Papadopoulos N, Kinzler KW, Vogelstein B (2011) Detection and quantification of rare mutations with massively parallel sequencing. P Natl Acad Sci USA 108(23):9530–9535.

9. Hiatt JB, Pritchard CC, Salipante SJ, O’Roak BJ, Shendure J (2013) Single molecule molecular inversion probes for targeted, high-accuracy detection of low-frequency variation. Genome Res 23(5):843–854.

10. Lou DI, et al. (2013) High-throughput DNA sequencing errors are reduced by orders of magnitude using circle sequencing. P Natl Acad Sci USA 110(49):19872–19877.

11. Hong J, Gresham D (2017) Incorporation of unique molecular identifiers in truseq adapters improves the accuracy of quantitative sequencing. BioTechniques 63(5):221–226.

12. Gregory MT, et al. (2016) Targeted single molecule mutation detection with massively parallel sequencing. Nucleic Acids Res 44(3):e22–e22.

13. Schmitt MW, et al. (2012) Detection of ultra-rare mutations by Next-Generation Sequencing. P Natl Acad Sci USA 109(36):14508–14513.

14. Wang K, et al. (2016) Ultra-precise detection of mutations by droplet-based amplification of circularized DNA. BMC Genomics 17(1):214.

15. Hoang ML, et al. (2016) Genome-wide quantification of rare somatic mutations in normal human tissues using massively parallel sequencing. P Natl Acad Sci USA 113(35):9846– 9851.

16. Jee J, et al. (2016) Rates and mechanisms of bacterial mutagenesis from maximum-depth sequencing. Nature 534(7609):693.

17. Kennedy SR, et al. (2014) Detecting ultralow-frequency mutations by Duplex Sequencing. Nat Protoc 9(11):2586.

18. Jinek M, et al. (2012) A programmable dual-RNA–guided DNA endonuclease in adaptive bacterial immunity. Science 337(6096):816–821.

19. Tsai SQ, Joung JK (2016) Defining and improving the genome-wide specificities of CRISPR–Cas9 nucleases. Nat Rev Genet 17(5):300.

20. Flint J, Harding RM, Boyce AJ, Clegg JB (1998) The population genetics of the haemoglobinopathies. Baillière’s Clin Haem 11:1–51.

21. Borg J, Georgitsi M, Aleporou-Marinou V, Kollia P, Patrinos GP (2009) Genetic recombination as a major cause of mutagenesis in the human globin gene clusters. Clin Biochem 42(18):1839–1850.

22. Allison AC (1964) Polymorphisms and natural selection in human populations. Cold Spring Harb Symp Quant Biol 29:137–149.

23. Hill AV, et al. (1991) Common West African HLA antigens are associated with protection from severe malaria. Nature 352:595–600.

24. Serjeant G, Serjeant B (2001) Sickle Cell Disease. (Oxford University Press).

25. Hardison RC, et al. (2002) HbVar: a relational database of human hemoglobin variants and thalassemia mutations at the globin gene server. Hum Mutat 19(3):225–233.

26. Hardison R, Miller W (2002) “Globin Gene Server,” http://globin.cse.psu.edu/ Accessed 10/5/2019.

27. Orkin SH (1990) Globin gene regulation and switching: circa 1990. Cell 63(4):665–672.

28. Steinberg M, Adams JI (1991) Hemoglobin A2: origin, evolution, and aftermath. Blood 78(9):2165–2177.

29. Fowler DM, Fields S (2014) Deep mutational scanning: a new style of protein science. Nat Methods 11(8):801.

30. Melamed D, Young DL, Gamble CE, Miller CR, Fields S (2013) Deep mutational scanning of an RRM domain of the *Saccharomyces cerevisiae* poly(A)-binding protein. RNA 19(11):1537–1551.

31. Tubbs A, Nussenzweig A (2017) Endogenous DNA damage as a source of genomic instability in cancer. Cell 168(4):644–656.

32. Ohno M, et al. (2014) 8-oxoguanine causes spontaneous *de novo* germline mutations in mice. Sci Rep 4(1):1–9.

33. Arbeithuber B, Makova KD, Tiemann-Boege I (2016) Artifactual mutations resulting from DNA lesions limit detection levels in ultrasensitive sequencing applications. DNA Res 23(6):547–559.

34. Costello M, et al. (2013) Discovery and characterization of artifactual mutations in deep coverage targeted capture sequencing data due to oxidative DNA damage during sample preparation. Nucleic Acids Res 41(6):e67–e67.

35. Bruskov VI, Malakhova LV, Masalimov ZK, Chernikov AV (2002) Heat-induced formation of reactive oxygen species and 8-oxoguanine, a biomarker of damage to DNA. Nucleic Acids Res 30(6):1354–1363.

36. Cheng KC, Cahill DS, Kasai H, Nishimura S, Loeb LA (1992) 8-hydroxyguanine, an abundant form of oxidative DNA damage, causes G*>*T and A*>*C substitutions. J Biol Chem 267(1):166–172.

37. Chen G, Mosier S, Gocke CD, Lin MT, Eshleman JR (2014) Cytosine deamination is a major cause of baseline noise in next-generation sequencing. Mol Diagn Ther 18(5):587– 593.

38. Wang RYH, Kuo KC, Gehrke CW, Huang LH, Ehrlich M (1982) Heat and alkali-induced deamination of 5-methylcytosine and cytosine residues in DNA. BBA Gene Struct Expr 697(3):371–377.

39. Duncan BK, Miller JH (1980) Mutagenic deamination of cytosine residues in DNA. Nature 287(5782):560–561.

40. Allinson SL, Dianova II, Dianov GL (2001) DNA polymerase *β* is the major dRP lyase involved in repair of oxidative base lesions in DNA by mammalian cell extracts. EMBO J 20(23):6919–6926.

41. Turk PW, Weitzman SA (1995) Free radical DNA adduct 8-oh-deoxyguanosine affects activity of Hp a II and Msp I restriction endonucleases. Free Radical Res 23(3):255–258.

42. Wood ML, Dizdaroglu M, Gajewski E, Essigmann JM (1990) Mechanistic studies of ionizing radiation and oxidative mutagenesis: genetic effects of a single 8-hydroxyguanine (7-hydro-8-oxoguanine) residue inserted at a unique site in a viral genome. Biochemistry 29(30):7024–7032.

43. Le Page F, et al. (2000) BRCA1 and BRCA2 are necessary for the transcription-coupled repair of the oxidative 8-oxoguanine lesion in human cells. Cancer Res 60(19):5548–5552.

44. Hoppins JJ, et al. (2016) 8-Oxoguanine affects DNA backbone conformation in the EcoRI recognition site and inhibits its cleavage by the enzyme. PloS One 11(10).

45. Lu AL, Clark S, Modrich P (1983) Methyl-directed repair of DNA base-pair mismatches in vitro. P Natl Acad Sci USA 80(15):4639–4643.

46. Glenn T, Waller D, Braun MJ (1994) Increasing proportions of uracil in DNA substrates increases inhibition of restriction enzyme digests. BioTechniques 17(6):1086–1090.

47. Koizume S, Inoue H, Kamiya H, Ohtsuka E (1998) Neighboring base damage induced by permanganate oxidation of 8-oxoguanine in DNA. Nucleic Acids Res 26(15):3599–3607.

48. Haas BJ, et al. (2011) Chimeric 16S rRNA sequence formation and detection in Sanger and 454-pyrosequenced PCR amplicons. Genome Res 21(3):494–504.

49. Bradley RD, Hillis DM (1997) Recombinant DNA sequences generated by PCR amplification. Mol Biol Evol 14(5):592–593.

50. Holcomb C, et al. (2014) Next-generation sequencing can reveal in vitro-generated PCR crossover products: some artifactual sequences correspond to HLA alleles in the IMGT/HLA database. Tissue Antigens 83(1):32–40.

51. Shendure J, Akey JM (2015) The origins, determinants, and consequences of human mutations. Science 349(6255):1478–1483.

52. Kong A, et al. (2012) Rate of *de novo* mutations and the importance of father’s age to disease risk. Nature 488(7412):471.

53. Campbell CD, Eichler EE (2013) Properties and rates of germline mutations in humans. Trends Genet 29(10):575–584.

54. Francioli LC, et al. (2015) Genome-wide patterns and properties of *de novo* mutations in humans. Nat Genet 47(7):822.

55. Rahbari R, et al. (2016) Timing, rates and spectra of human germline mutation. Nat Genet 48(2):126.

56. Goldmann JM, et al. (2016) Parent-of-origin-specific signatures of *de novo* mutations. Nat Genet 48(8):935.

57. Ségurel L, Wyman MJ, Przeworski M (2014) Determinants of mutation rate variation in the human germline. Annual Review of Genomics and Human Genetics 15:47–70.

58. Kondrashov AS (2003) Direct estimates of human per nucleotide mutation rates at 20 loci causing Mendelian diseases. Hum Mutat 21(1):12–27.

59. Lynch M (2010) Rate, molecular spectrum, and consequences of human mutation. P Natl Acad Sci USA 107(3):961–968.

60. Turner TN, et al. (2017) Genomic patterns of *de novo* mutation in simplex autism. Cell 171(3):710–722.

61. Carlson J, et al. (2018) Extremely rare variants reveal patterns of germline mutation rate heterogeneity in humans. Nat Commun 9(1):3753.

62. Lander E, Linton L, Birren B, et al,. (2001) Initial sequencing and analysis of the human genome. Nature 409:860–921.

63. Karczewski K, Francioli L, MacArthur D, et al,. (2020) The mutational constraint spectrum quantified from variation in 141,456 humans. Nature 581:434–443.

64. Yates A, Achuthan P, Akanni W (2020) Ensembl 2020. Nucleic Acids Research 48:D682– D688.

65. Weyrich A (2012) Preparation of genomic DNA from mammalian sperm. Curr Protoc Mol Biol 98(1):2–13.

